# Epigenetic control of S100A8/A9-driven monocytic inflammation licenses anti-leukemic functionality of immature NK cells during hematopoietic stem cell differentiation

**DOI:** 10.64898/2026.03.25.714180

**Authors:** R Schirrmann, D Stowitschek, M Sutter, J-H Lee, B Zhao, S Lee, A Neyazi, BF Broesamle, F Ginsberg, P Krammer, A Kübler, T Vogl, H Wittkowski, S Ahmad, B Krämer, N Peter, M Klimiankou, M Ritter, J Skokowa, D Atar, EM Mace, M Barroso Oquendo, N Casadei, N Guengoermues, R Handgretinger, FC Jones, U Holzer, MC André

## Abstract

Inflammation is a key driver of hematopoietic dysfunction in myeloid malignancies, but its role in the context of hypomethylating therapy remains incompletely understood. Although 5-Azacytidine is used posttransplant in high-risk myelodysplastic syndrome (MDS), only 50% of patients show a clinical response. We provide evidence that inherent inflammatory properties of healthy donor CD34^+^ stem cells exist that are likely to contribute to the “response” seen in MDS patients. These are linked to epigenetic priming of the myeloid niche, resulting in S100A8/A9-driven inflammatory program that promotes functionality of immature NK cells. Using in vitro differentiation systems, multi-omic profiling, and a S100A9^−/−^ mouse model, we find that 5-AzaC modulates inflammatory transcriptional programs through epigenetic rewiring of upstream regulatory elements. Loss of S100A9 disrupts myeloid differentiation, impairs NK cell maturation, and alters key developmental regulators including CEBPB, JUN, and NFIL3. In vivo, 5-AzaC restores these defects and primes NK cells in a time- and context-dependent manner. Re-analysis of the published Australian MDS/CMML cohort shows that “responders” display increased S100A8/A9 expression together with enhanced IFN-γ, IL6-JAK-STAT3, and TNF signaling. These findings suggest that inflammatory myeloid programs may serve as predictive biomarkers and therapeutic targets to enhance NK cell–mediated graft-versus-leukemia activity posttransplant.

**Summary:**

1. We provide compelling evidence that inherent properties of healthy donor CD34^+^ hematopoietic stem cells (SCs) exist that are likely to contribute to the “response” seen upon pre-emptive posttransplant 5-AzaC therapy of patients with high-risk myelodysplastic syndrome (MDS).
2. These properties are linked to a distinct form of epigenetic plasticity at upstream-located transcription factor (TF) binding sites. This may indirectly contribute to acute S100A8/A9-driven inflammation, which is demonstrable in distinct monocyte subsets and, importantly, also in NK cells thereby determining the characteristics of inflammatory monocyte-NK cell crosstalk.
3. Mice with a targeted deletion of S100A9 fail to upregulate CEBPB / JUN and NFIL3 which results in impaired myeloid priming and dysfunctional NK cell maturation, respectively.
4. Re-analysis of the Australian MDS/CMML cohort confirms that MDS patients that “respond” to 5-AzaC exhibit activated IFN-γ, IL6-JAK-STAT3, and TNF-signaling pathways in the context of upregulated S100A8/A9 after six months of treatment.
5. Our study indicates that screening of healthy donors SCs for specific inflammatory markers in early developing monocytes could be used as a marker to predict which donor will have the potential of generating a S100A8/A9-driven inflammatory response. This may help identify patients with MDS as well as AML who are likely to benefit from low-dose, short-term 5-AzaC therapy as early as day 7 after transplantation, potentially resulting in increased graft-versus-leukemia (GvL) activity.

## Introduction

Chronic inflammation is a central driver of myeloid disease progression, spanning clonal hematopoiesis of indeterminate potential (CHIP) to myelodysplastic syndromes (MDS) and acute myeloid leukemia (AML)^1–4^. In MDS, inflammatory remodeling of the bone marrow (BM) niche is characterized by the emergence of cytokine-producing stromal populations and elevated levels of alarmins such as S100A8 and S100A9, myleloid proteins, which have been implicated as key mediators linking extracellular danger signals to inflammasome activation and disease propagation^3,5^.

Therapeutic strategies for high-risk MDS, chronic myelomonocytic leukemia (CMML) and AML commonly aim to modulate aberrant inflammatory and epigenetic programs within the myeloid compartment. The hypomethylating agent 5-azacytidine (5-AzaC) is a cornerstone of treatment and induces epigenetic reprogramming through inhibition of DNA methyltransferases^6,7^ (Christman et al., Stresemann et al.). In addition to its direct effects on malignant cells, 5-AzaC has been reported to modulate innate immune compartments, including natural killer (NK) cells. Specifically, 5-AzaC can induce the expression of killer immunoglobulin-like receptors (KIRs) and alter NK cell differentiation states^8,9^. However, the functional consequences of these changes remain incompletely understood, as both enhanced and impaired NK cell cytotoxicity following 5-AzaC exposure have been reported across experimental systems^10–13^.

Natural killer cells are key mediators of graft-versus-leukemia (GvL) activity following allogeneic stem cell transplantation, and their function is tightly regulated by the balance of activating and inhibitory receptors, including KIRs^14^. Using a donor-specific xenotransplantation model in humanized NOD-SCID IL2Rγc⁻/⁻ (NSG) mice, we previously investigated NK cel-–mediated anti-leukemic responses in the setting of minimal residual disease^15–17^. In this context, low-dose 5-AzaC administration during early hematopoietic stem cell differentiation unexpectedly resulted in the expansion of immature NK cell precursors lacking KIR expression yet exhibiting enhanced anti-leukemic activity^17^. These findings suggested that 5-AzaC may modulate NK cell function at earlier stages of hematopoietic differentiation than previously appreciated. However, the cellular and molecular mechanisms linking epigenetic therapy and NK cell functional programming remain poorly defined.

Here, we show that 5-AzaC reshapes the inflammatory bone marrow niche to reprogram NK cell differentiation at early hematopoietic stages, thereby generating functionally superior immature NK cells with enhanced anti-leukemic activity. Mechanistically, this process is driven by S100A8/A9-mediated inflammatory signaling from monocytes to NK cells. These findings suggest that inflammatory myeloid programs may serve as predictive biomarkers and therapeutic targets to *enhance* NK cell–mediated graft-versus-leukemia activity posttransplant.

## Results

### Low-dose 5-AzaC induces a S100A8/A9-driven inflammatory response during CD34⁺ SC differentiation that promotes anti-leukemic functionality of immature NK cells in “responders”

To explore earlier findings by our group that linked exposure to low-dose 5-AzaC to improved anti-leukemic function of immature NK cells^17^, we established an in vitro system based on a protocol by the Dolstra group^18^ by differentiating healthy CD34⁺ SC into immature NK cells in a medium containing a complex cytokine mixture **(Fig. 1A)**. Based on the analysis of various cell surface molecules^19^ and extensive titration of 5-AzaC (data not shown), we chose day 31 of culture for the analysis of NK cell function to allow the characterization of immature as opposed to mature NK cell function. We observed that 5-AzaC enhanced the cytotoxicity of immature NK cells in a subset but not all SC donors (**Fig. 1B**). Interestingly, immature NK cells from “responders” displayed a somewhat lower baseline cytotoxic activity in the absence of 5-AzaC than “non-responders”. However, upon 5-AzaC exposure, immature “responder” NK cells increased their level of cytotoxicity comparatively more than “non-responder”, an effect that correlated well with increased migratory properties towards K562-derived tumor supernatants **(Fig. 1C).** This effect was TRAIL-dependent, as their ability to lyse Jurkat cells was reduced by blocking Ca2^+^-dependent signaling with EGTA/MgCl_2_ **(Fig. 1D).** Using multicolor flow cytometry, we documented that exposure to 5-AzaC did not alter overall cell proliferation rates (**Suppl. Fig. S1**); however, a significant number of myeloid precursors and myeloid lineage-committed cells existed next to lymphoid precursors and lineage-committed NK cells in our culture suspensions on day 17 and day 31 (**Suppl. Fig. S2**). Interestingly, the inclusion of adherently growing cells which probably corresponded to few mDCs and pDCs during medium change was crucial to achieve the 5-AzaC-induced effect on immature day 31 NK cell function (data not shown). As crucial epigenetically controlled expression of EOMES and T-bet (TBX21) determines early NK cell-lineage-specification^20^, we next quantified the expression of key transcription factors (TF) involved in early NK development (**Suppl. Fig. S3**).

**Fig. 1.**
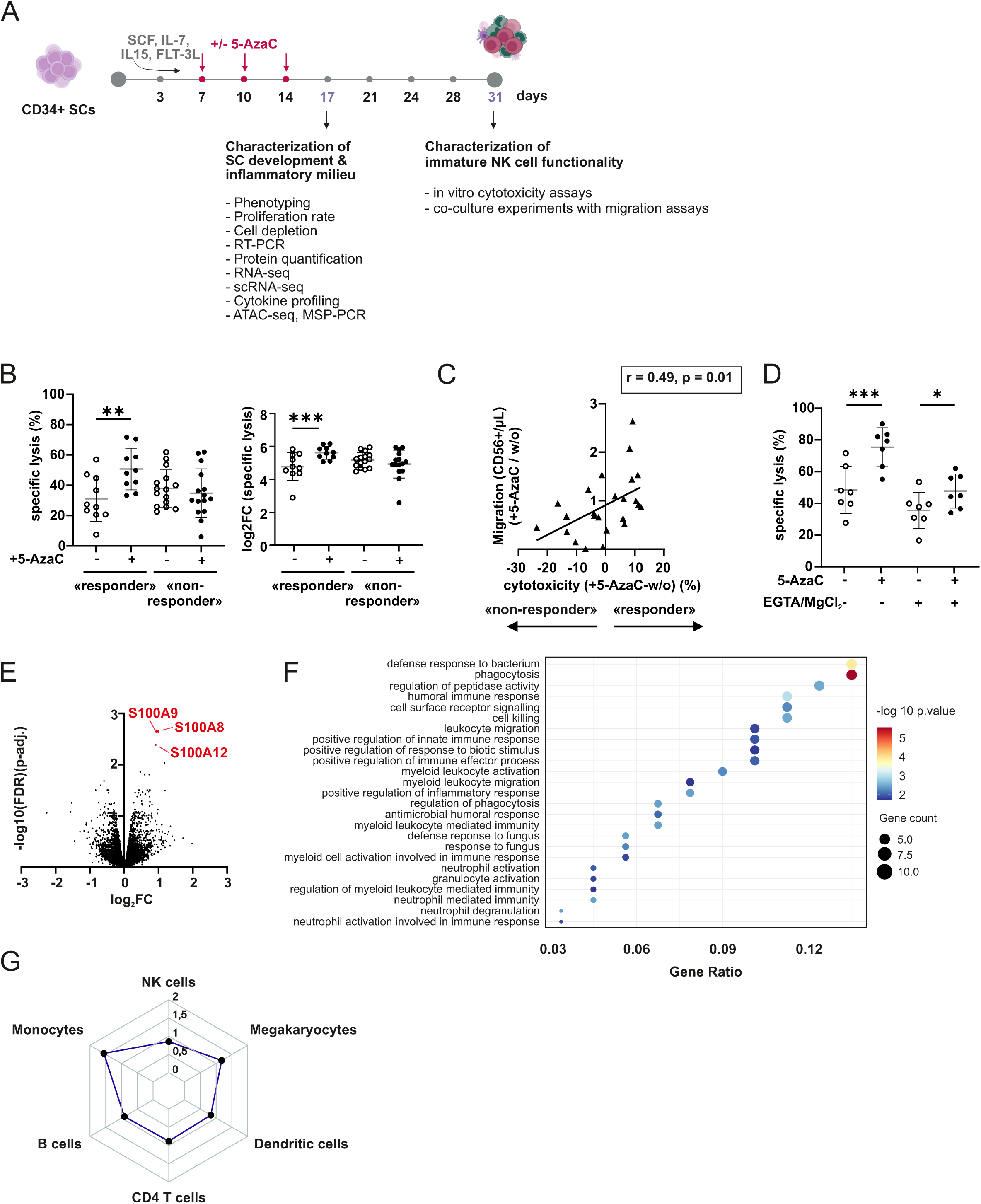
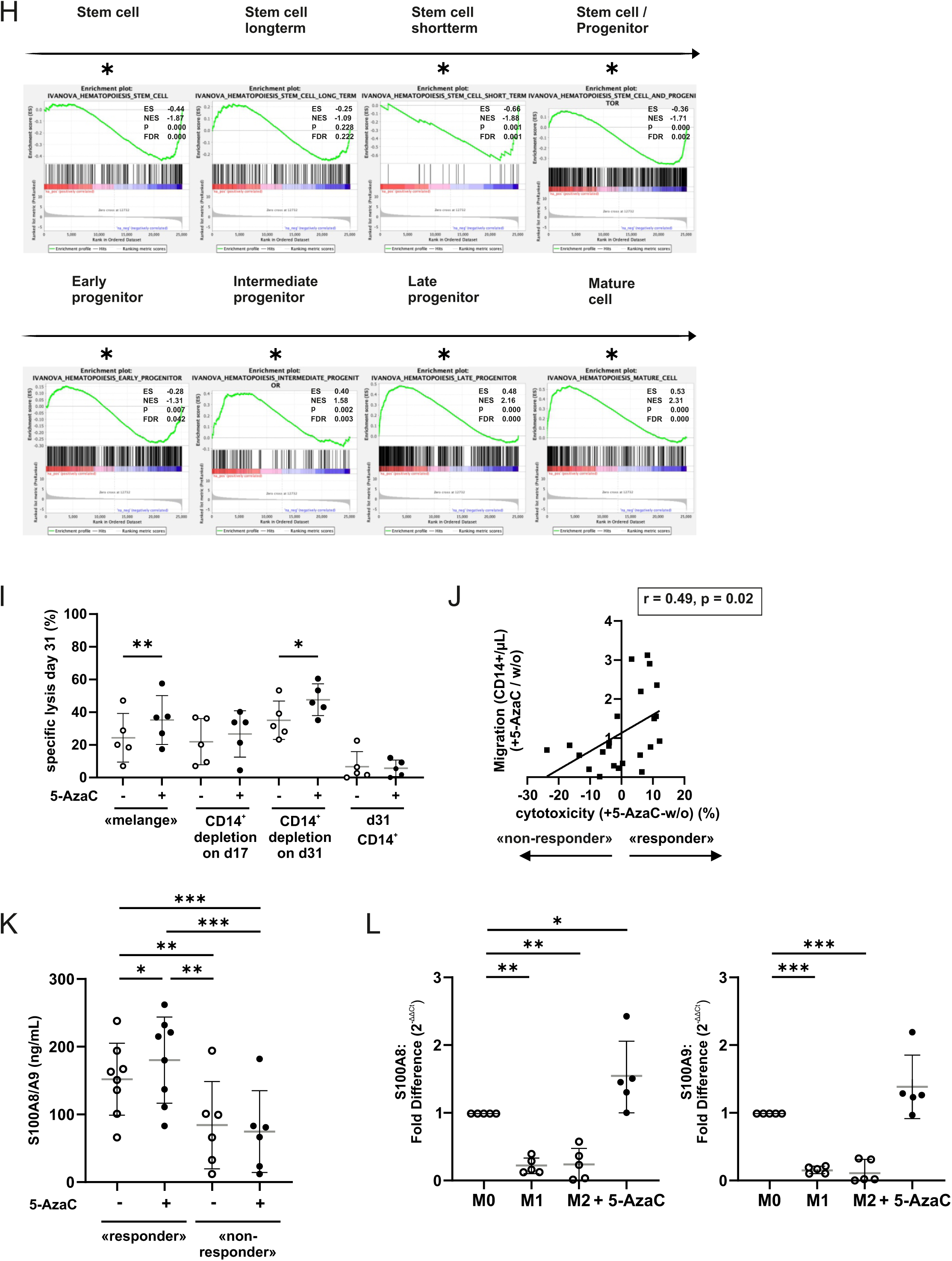
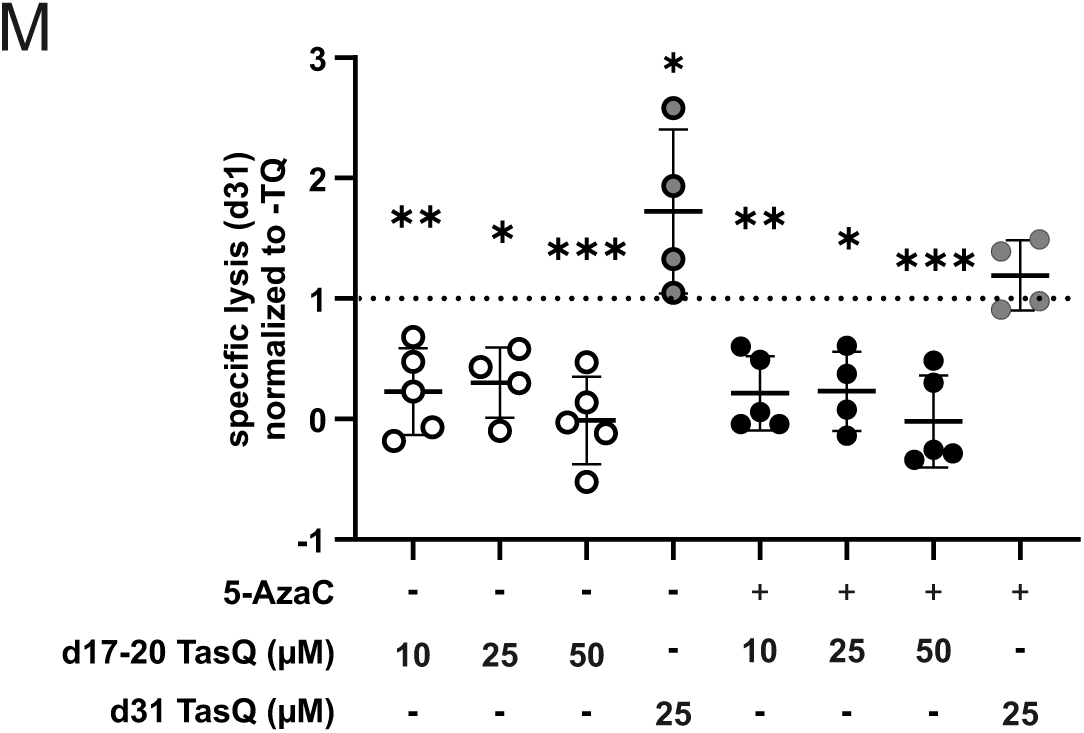
Low-dose 5-AzaC during early CD34^+^ SC development elicits a S100A8/A9-driven inflammatory response that supports anti-leukemic functionality of immature NK cells. **(A**) Experimental design of in vitro differentiation of human CD34^+^ developing stem cells (SCs). See also Supplemental Methods and Figures section. **(B)** Immature NK cells on day 31 of SC culture were tested for cytotoxicity against K562 targets (E:T 50:1). Shown are specific lysis (%) and log2 fold change (FC) upon 5-AzaC treatment in “responders” (n = 10) and “non-responders” (n = 15). Statistical significance was assessed by two-way repeated-measures ANOVA with Sidak’s multiple-comparisons test. Data were log2-transformed prior to analysis. Mature NK cell functionality typically arises later during culture (days 35–42) (Spanholtz et al.). **(C)** Immature NK cell migration toward K562-derived supernatant was assessed in a transwell migration assay. Donors were classified as “responders” or “non-responders” based on their capacity to increase NK cell cytotoxicity upon 5-AzaC treatment. The y-axis shows the fold change (FC) in CD56^+^ cell migration in 5-AzaC-treated vs. untreated conditions. Migration was positively correlated with responder status (Pearson correlation, r = 0.49, n = 24, p = 0.01). **(D)** Immature NK cells from on day 31 of “responder” cultures were tested for cytotoxicity against Jurkat target cells (E:T 10:1). Shown is specific lysis (%) in the presence or absence of Ca²⁺ chelation with EGTA/MgCl₂ to assess TRAIL-dependent killing. Shown is specific lysis (%) in the presence or absence of Ca²⁺ chelation with EGTA/MgCl₂. Statistical significance was determined by one-way ANOVA with Šídák’s multiple-comparisons test (n = 7). **(E–H)** RNAseq analysis of 5-AzaC-treated CD34⁺ SCs (day 17) from “responders” (n = 4). **(E)** Volcano plot of differential transcript expression between 5-AzaC-treated vs. control samples. Each point represents a transcribed gene; the x-axis indicates log₂ fold change (FC) (5-AzaC/control) and the y-axis indicates -log_10_(FDR)(p-adj). Note that S100A8, S100A9 and S100A12 are highlighted as among the most significantly differentially expressed transcripts. Differential expression was assessed using DESeq2 (Wald test with Benjamini–Hochberg FDR correction). **(F)** Gene Ontology (GO) enrichment analysis of Biological Process (BP) terms comparing 5-AzaC treated vs. untreated controls. Dot color indicates −log10(p-value), dot size indicates gene count (number of overlapping genes), and gene ratio indicates the proportion of input genes that overlap with the GO term. The top 25 significantly enriched terms are shown. Enrichment significance was assessed using a one-sided hypergeometric (Fisher’s exact) over-representation test with Benjamini–Hochberg correction. **(G)** Spider plot showing the FC (5-AzaC/control) in inferred cell-type fractions estimated by digital cytometry (CIBERSORTx) in treated samples relative to matched untreated controls. **(H)** Pre-ranked gene set enrichment analysis (GSEA) of IVANOVA hematopoietic gene sets comparing 5-AzaC–treated and control samples. Normalized enrichment scores (NES) and FDR-adjusted q values for late progenitor- and mature cell–associated programs are shown. Significance was assessed using 10.000 gene set permutations. **(I)** CD14^+^ monocytes were depleted from SC differentiation cultures (“mélange”) either on day 17 or day 31, and NK cell cytotoxicity against K562 target cells was assessed on day 31 as in Fig. 1B (n = 5; one-way ANOVA with Šídák’s multiple-comparisons test). **(J)** Transwell migration of CD14^+^ monocytes toward K562-derived supernatant, shown as fold change (5-AzaC/control) in “responder” and “non-responder” donors (Pearson correlation, r = 0.49, n = 24, p = 0.02). **(K)** Protein concentrations of S100A8 and S100A9 in day 17 SC cultures from 5-AzaC-treated and uncontrolled samples, quantified by ELISA in “responders” (n = 8) and “non-responders” (n = 6) (mixed-effects analysis with Tukey’s multiple-comparisons test). **(L)** S100A8 and S100A9 mRNA expression in 5-AzaC-treated THP-1 cells, with M1- and M2-polarized macrophages (see also Methods section) included for comparison, n = 5. Mixed-effects analysis with Tukey’s multiple comparisons test. **(M)** Cytotoxicity of immature NK cells on day 31 in the presence or absence of S100A8/A9 blockade with Tasquinimod (TasQ). TasQ was added either during differentiation (days 17–20) or during the cytotoxicity assay on day 31 (n = 4–5, one-way ANOVA with Tukey’s multiple comparisons test). Ratio of TasQ-treated to control conditions. Significance is indicated as *p ≤ 0.05, **p ≤ 0.01, ***p ≤ 0.001. n values denote independent experiments using different human SC donors.

However, apart from *SPI1* (PU.1), which was significantly upregulated, none of the other TFs showed any change. As the role of *SPI1* seems to be more permissive and less instructive for NK cell lineage-commitment than the other TFs determined by us^21^, we asked whether the 5-AzaC-induced control of immature NK cell function might be an indirect one. Bulk RNA sequencing of 5-AzaC-treated day 17 “responder” SC cultures documented that few (< 100) genes that were significantly upregulated in our culture system but unexpectedly none of these were related to NK cell differentiation. On the contrary, *S100A8, S100A9,* and *S100A12* genes were the genes with the lowest false discovery rates (FDR) clustering among the highest log fold changes (FC) of all differentially expressed genes **(Fig. 1E, Suppl. Fig. S4, Suppl. Tables 1-4).** Although a limited number of additional monocyte-associated genes (i.e., *MNDA, MARCO, MEFV, CD14, CD300C, FPR2,* and *LILRB2*) were also upregulated, their FC were substantially smaller and associated with higher FDRs. We next analyzed DNA methylation profiles in different blood cell types^22^ and confirmed existing knowledge (www.uniprot.org) that the promotor region and the gene body of S100A8 and S100A9 are demethylated in CD16^+^ neutrophils, neutrophils in general, and CD14^+^ monocytes, but not in NK cells or CD34^+^ SCs (**Suppl. Fig. S5**).

Gene ontology (GO) analysis d17 responder bulk RNA demonstrated a strong enrichment of inflammatory and innate immune response pathways **(Fig. 1F)**. Digital cytometry analysis using CIBERSORTx^23^ evidenced an abundance of myeloid cells in the 5-AzaC-treated cultures on day 17 **(Fig. 1G).** Gene set enrichment analysis (GSEA) revealed a decrease in transcriptional programs associated with SCs, progenitors and early progenitor-like states in 5-AzaC treated “responder” cultures (**Fig. 1H**). Importantly, the depletion of CD14^+^ monocytes on day 17 but not day 31 from our cell culture suspension (“mélange”) completely abrogated any beneficial effect of 5-AzaC on immature NK cell function (**Fig. 1I**), indicating that monocyte-to-NK cell crosstalk was required to instruct emerging immature NK cells. In line with the superior migratory properties of immature NK cells (Fig. 1C), emerging CD14^+^ monocytes from 5-AzaC-treated “responders” displayed higher migratory function in towards K562-derived supernatants (**Fig. 1J**). Quantification of *S100A8* and *S100A9* protein expression confirmed significantly higher concentrations in supernatants of 5-AzaC-treated cultures in “responders” **(Fig. 1K).** Interestingly, S100A8 mRNA upregulation was also demonstrable when treating the human immortalized monocyte-like THP-1 cell line with 5-AzaC (**Fig. 1L**). Blocking of S100A8/A9 with Tasquinimod during SC differentiation (day 17-20) but importantly not during NK cell-to-target cell (K562) recognition (day 31) abrogated the earlier described beneficial effect of 5-AzaC exposure on immature NK cell anti-leukemic functionality (**Fig. 1M, Suppl. Fig. S6A-C**). Collectively, these data indicate that early S100A8/A9-driven monocytic inflammation in 5-AzaC treated SC cultures may enhance immature NK cell function on day 31 in “responder” SC donors.

### 5-AzaC responders exhibit an expanded S100A8/A9⁺ monocytic cluster (InflMo) associated with enhanced inflammatory signaling towards immature NK cells

We next performed single-cell RNA (scRNA)-sequencing on day 17 of 5-AzaC treated SC cultures from “responders” and “non-responders”. Based on canonical marker gene expression (**Suppl. Fig. S7**), unsupervised clustering allowed the identification of 14 transcriptionally distinct immune subsets (**Fig. 2A**). Interestingly, the analysis of the relative abundance of all annotated clusters in day 17 cultures showed a large monocytic compartment comprising of numerous clusters of monocytic cells in varying inflammatory states (**Fig. 2B**) that inherently differed between “responders” and “non-responders” with a large inflMo cluster in “responders” and abundantly represented hyinflMo and IntMo clusters in “non-responders” (**Fig. 2C, Suppl. Fig. 8**). Monocle inferred a trajectory that was manually rooted in GMPs, from which inflMo and hyinflMo emerged as divergent branches suggesting that cluster differences between “responders” and “non-responders” reflect distinct differentiation-associated programs (**Fig. 2D, Suppl. Fig. S9**). Importantly, 5-AzaC treatment did not appreciably change the inferred trajectory topology of both “responders” and “non-responders” **(Fig. 2E**).

**Fig. 2.**
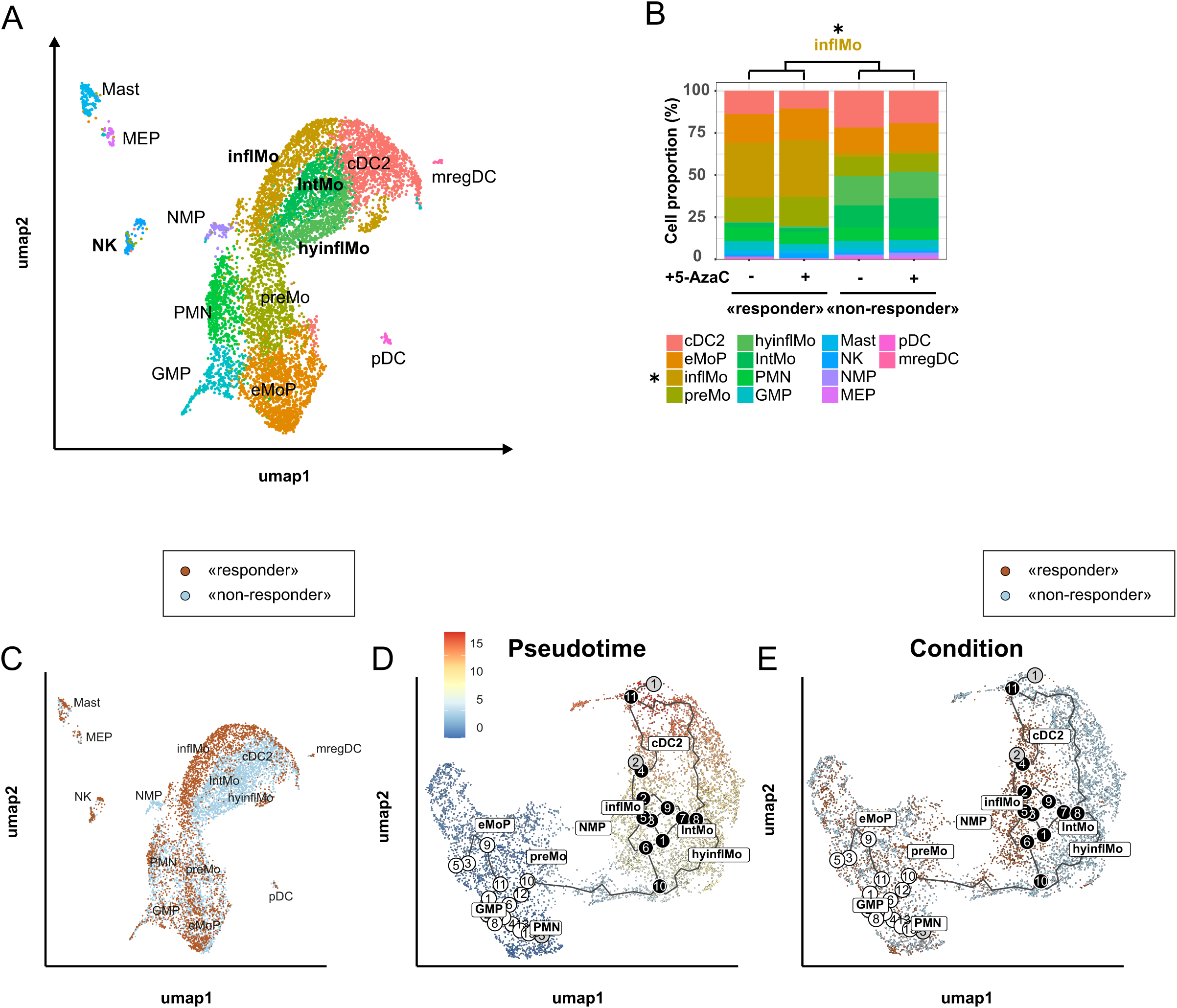
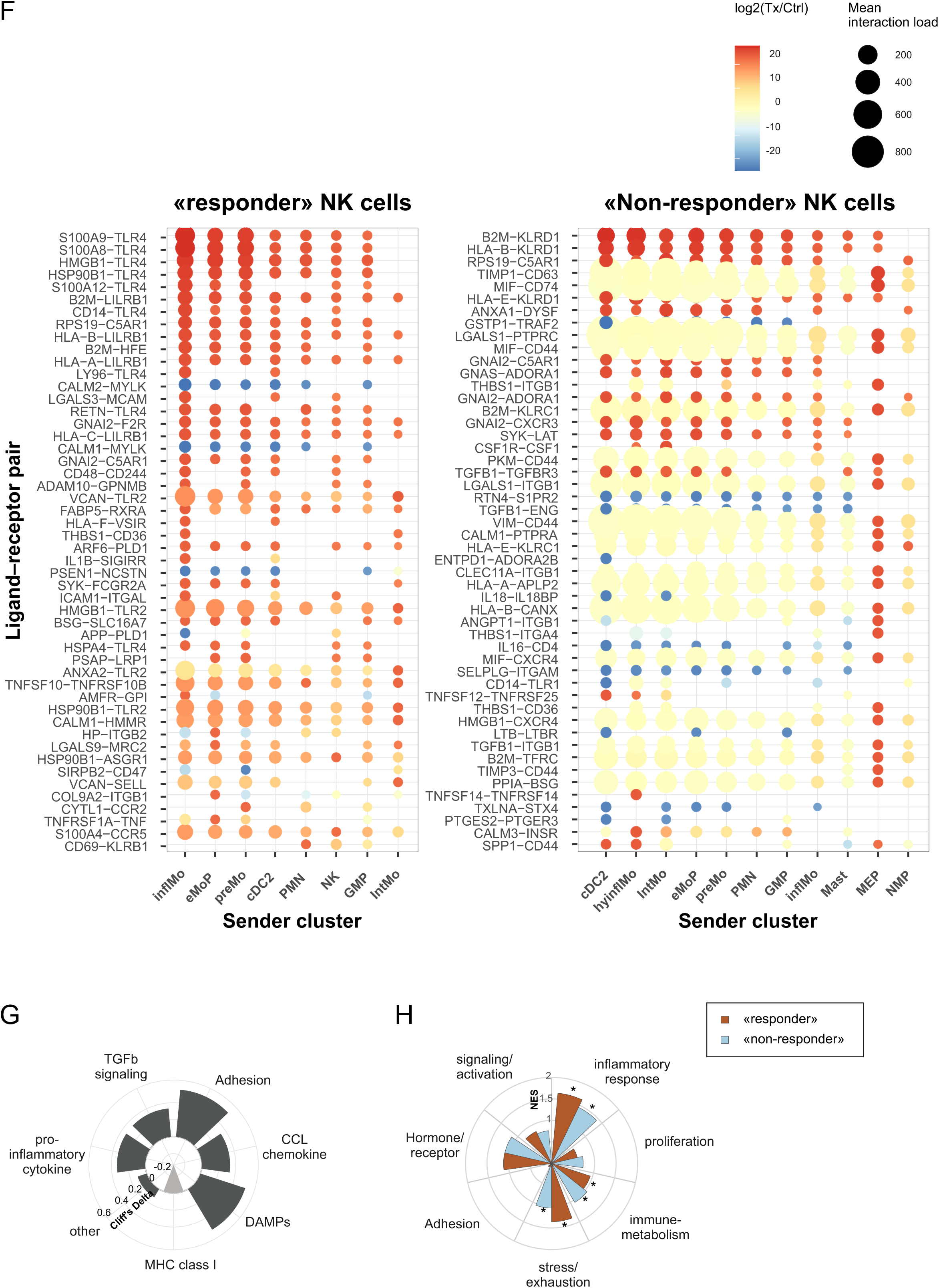
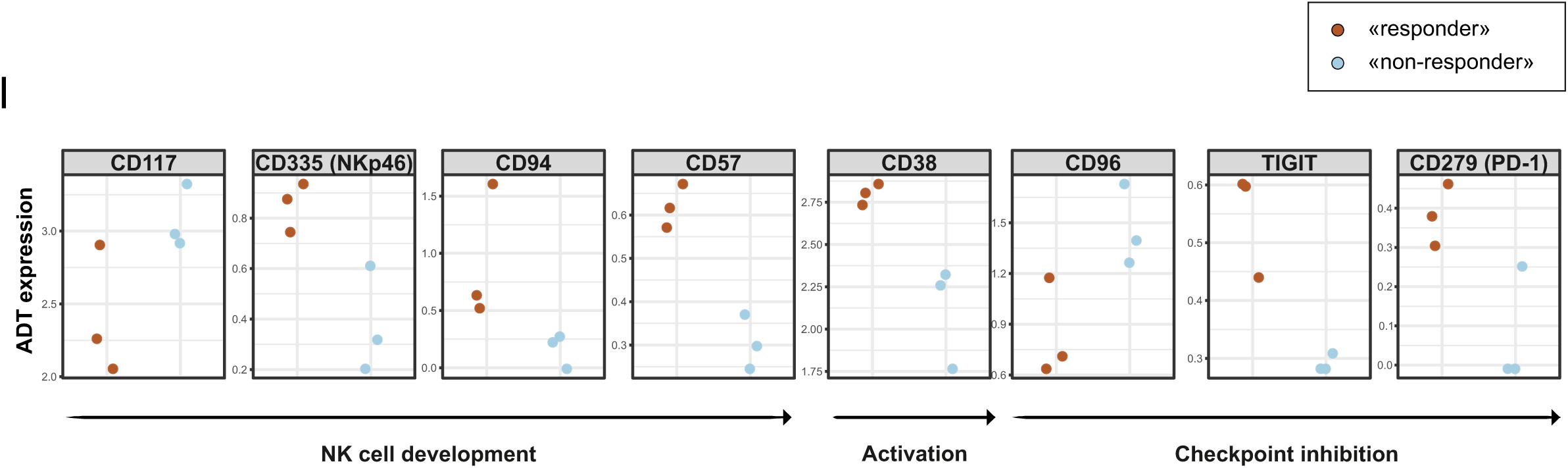
Distinct inflammatory monocyte signatures and NK cell interaction networks in 5-AzaC “responders”. **(A)** Uniform Manifold Approximation and Projection (UMAP) of single-cell transcriptomes from day 17 cultures of “responders” and “non-responders”, pooled across treatment conditions (n=3 “responder”, n=3 “non-responder”), colored by annotated clusters. Cells are colored by unsupervised clusters and annotated as indicated: Polymorphonuclear neutrophils (PMN), conventional dendritic cells type 2 (cDC2), plasmacytoid dendritic cells (pDC), mature-regulatory dendritic cells (mregDC), granulocyte-monocyte progenitor (GMP), early monocyte progenitor (eMoP), pre-monocyte (preMo), neutrophil-monocyte progenitor (NMP), mast cells (Mast), NK cells (NK) and megakaryocyte-erythroid progenitor (MEP), inflMo = inflammatory monocyte, IntMo = intermediate monocyte, hyinflMo = hyperinflammatory monocyte. **(B)** Stacked bar plot showing the relative abundance of annotated clusters across donor groups and treatment conditions (one-sided Welch’s t-test on donor-level mean proportions, *p < 0.05, unadjusted). **(C)** UMAP colored by donor group. **(D, E)** Monocle3 trajectory analysis of myeloid differentiation toward inflammatory monocyte subsets and cDC2s, shown by **(D)** Pseudotime and **(E)** Condition. **(F)** Differentially regulated ligand–receptor pairs between treatment and control conditions across sender clusters engaging NK cells. Bubble size represents the interaction strength, the color the log2 fold change. Included are only LR pairs with detectable LR expression ≥ 0.05, an interaction score ≥ 0.05, that is present in at least 2 donors, and sender clusters with ≥ 10 cells per donor and condition. **(G)** Effect sizes of 5-AzaC–induced changes in inferred ligand–receptor interaction categories between monocyte subsets and NK cells, summarized as Cliff’s delta comparing “responders” and “non-responders”. For each donor, culture group (“responder” vs. “non-responder”), and category, the 5-AzaC-induced change in weighted interaction load was calculated as Δ = treatment − control, using the summed weighted scores across all ligand–receptor pairs within that category. The circle plot summarizes effect sizes as Cliff’s delta, comparing Δ values between “responders” and “non-responders” for each category. Effect sizes were interpreted as small for |δ| > 0.14, medium for |δ| > 0.33, and large for |δ| > 0.47. **(H)** Gene set enrichment analysis (GSEA) of Hallmark pathways summarized by functional super-categories in NK cells (Wilcoxon signed-rank test with Benjamini–Hochberg correction; *FDR < 0.05). **(I)** Donor-level ADT expression of selected NK cell markers in “responders” and “non-responders” under 5-AzaC treatment (n=3 per group).

Using GSEA and exploratory differential gene expression (DGEA) in pooled comparisons of inflMo versus hyinflMo and IntMo, we showed significant enrichment of interferon (IFN)-α/γ responses as well as KRAS signaling in inflMo. InflMo showed a significant negative enrichment of coagulation, TGF-β signaling, mTORC1 signaling, angiogenesis, epithelial–mesenchymal transition, and NOTCH signaling relative to intMo. When comparing hyinflMo with IntMo, pathways related to coagulation, adipogenesis, cholesterol homeostasis, glycolysis, mTORC1 signaling, hypoxia, and NOTCH signaling were significantly negatively enriched in hyinflMo (**Suppl. Fig. S10, Suppl. Tables 5-10**).

To understand differences in the individual transcriptional response to 5-AzaC, we performed GSEA as well as exploratory DGEA (treated/untreated; unadjusted p-value–based) (**Suppl. Fig. S11-12, Suppl. Tables 11-13, Suppl. Tables 14-16**). In the dominant inflMo cluster in “responders”, GSEA revealed downregulation of interferon-α/-γ signaling (**Suppl. Fig. S11**) which was accompanied by reduced expression of representative IFN- and inflammasome-associated genes in DGEA (**Suppl. Fig. 12**), including FCER1A, IFIT3, IFI44L, MX1, GPANK1, and NEK7. In contrast, the large IntMo cluster in “non-responders” displayed a decrease in CLEC10A, IL1B, MRC1, CNST, and IGF1, alongside an increase in TNNT1, ACSL1, PTX3, KRCC1, THBS1, and SPP1, suggesting a transformation from inflammatory signaling toward metabolic and extracellular matrix–associated pathways. The large hyinflMo cluster in “non-responders” also showed a downregulation of apoptosis- and KRAS signaling, accompanied by reduced expression of genes associated with activation, proliferation, and migratory properties, including *TMPO, SLC7A5, CD69, CX3CR1, EMP1, IL1B, CDIPT, CCND2, HOXB6, PLEK, ATIC*, and *ZNF160*. Collectively, these findings indicate that inflMo, IntMo, and hyinflMo represent functionally distinct monocytic clusters that undergo divergent transcriptional remodeling following 5-AzaC treatment.

We next scrutinized all clusters for their S100A8/A9 expression using donor-paired visualization (**Suppl. Fig. S13**). Interestingly, a modest increase of S100A8/A9 expression was demonstrable in all three monocytic clusters of 5-AzaC treated “responders”, i.e. in the large inflMo cluster and the minor hyinflMo and IntMo clusters. In line with our hypothesis that regulation of S100A8/A9 was inherently different in “non-responders”, we did not document an increase of S100A8/A9 expression in any of the three monocytic clusters in “non-responders”.

This selective remodeling of the monocytic compartment suggests that only inflMo retain an inflammatory capacity to instruct NK cell function, whereas IntMo and hyinflMo adopt transitional or regulatory monocytic programs that may be insufficient to support NK cell functional maturation.

NK cells from “responders” and “non-responders” also exhibited fundamentally different responses to 5-AzaC in both GSEA (**Suppl. Fig S14, Tables 17-18**) as well as exploratory DGEA (unadjusted p-value–based) (**Suppl. Fig. S15, Tables 19-20**). Both “responders” and “non-responders” NK cells displayed shared enrichment of inflammatory pathways, including allograft rejection, complement, IL-6/JAK–STAT signaling, general inflammatory response, and IFN-α/γ signatures. “Responders” uniquely engaged oxidative and stress-adaptive programs, these included oxidative phosphorylation, hypoxia, p53 signaling, unfolded protein response, and mTORC1 and PI3K-AKT-mTOR pathways, indicating successful coupling of inflammatory cues to metabolic reprogramming and cellular stress tolerance. “Responders” showed upregulation of *LGALS1/LGALS3, IFI30, AKIRIN1, NAIP, ATP6VOC, ATP9B*, and *CCNB1*, and downregulated expressions of *SPINK2, SNRNP200*, and *CDC6,* consistent with enhanced cytotoxic readiness and metabolic activation. “Non-responders”, by contrast, predominantly showed upregulated proliferation- and chromatin-associated genes including HIST1H4C, HIST1H1B/H1D, HIST1H3G, TYMS, TUBB, and CDK1, together with downregulated antigen-processing and lysosomal mediators (HLA-DQB1, HLA-DRA, CTSS, CST3, CTSH), suggesting a cycling rather than effector-maturation phenotype. Cluster-weighted LR-analysis (**Fig. 2F, Suppl. Fig. S17**) identified inflMo as the dominant interaction partner of “responder” NK cells, independent of treatment, whereas interactions in “non-responders” were distributed across multiple subsets. Delta-based comparisons showed that 5-AzaC promoted interactions from inflMo toward NK cells in “responders” via S100A8/A9–TLR4 signaling, whereas the overall modest increase in the interaction to NK cells in “non-responders” was conferred by B2M- or HLA-B–CD94 (KLRD1) interactions. In line with this, the effect size of 5-AzaC–induced changes in inferred ligand–receptor interaction categories between monocyte subsets and NK cells (Cliff’s delta) and GSEA of Hallmark pathwayssummarized by functional super-categories in NK cells further indicated that 5-AzaC supported adhesion and DAMP-related LR programs in “responders” and a relative reduction in MHC class I–mediated interactions (**Fig. 2G, H**). CITEseq (antibody-derived tag, ADT) profiling of NK cells (**Fig. 2I**) with donor-level aggregation of centered log ratio (CLR)-normalized ADT expression revealed that “responder” NK cells exhibited a consistently more mature and activated protein phenotype, with higher expression of CD94, CD57, CD38, and the activating receptor NKp46, alongside reduced expression of the immature marker CD117. In parallel, “responders” displayed increased TIGIT and PD-1 expression and reduced CD96 levels. Collectively, these data indicate that 5-AzaC elicits a broadly shared inflammatory activation in NK cells but only “responders” successfully integrate this signal into metabolic reprogramming, stress tolerance, and effector maturation. In contrast, “non-responders” remain confined to a proliferative yet functionally ineffective state, providing a mechanistic explanation for divergent functionality.

### 5-AzaC reshapes myeloid chromatin accessibility at upstream transcription factor regulatory elements while sparing the S100A8/A9 locus

*To* characterize S100A8/A9 epigenetic regulation, we first evaluated global DNA methylation levels by 5-methylcytosine staining, showing an overall reduction in methylation following treatment which was maximal on day 17 of culture **(Suppl. Fig. S18A).** As global measurements do not allow the distinction between functional demethylation of a gene and random demethylation of repetitive non-coding DNA, we analyzed locus-specific DNA methylation at *S100A8* and *S100A9* **(Suppl. Fig. S18B).** In “responders”, 5-AzaC treatment resulted in a modest but reproducible reduction in DNA methylation at selected CpG-rich regions of *S100A8*, with the most pronounced and statistically significant changes detected within CpG rich regions of the gene body **(Suppl. Fig. S18C)** but not in the promoter-proximal regions. In “non-responders” no significant demethylation of *S100A8* CpG islands or gene body was observed **(Suppl. Fig. S18D).** In addition, we found that the *S100A9* gene displayed an overall high level of demethylation particularly at promoter region in both “responders” and “non-responders” irrespective of 5-AzaC treatment **(Suppl. Fig. S18E-G).**

To characterize chromatin accessibility, we performed an Assay for Transposase-Accessible Chromatin with sequencing (ATAC-seq) on day 17 cultures from “responders”, analyzing flow cytometry-sorted CD14⁺ monocytes. In line with the notion that a large-scale loss of DNA methylation is a critical step in myeloid lineage fate decisions^24^, we found that 5-AzaC treatment induced a global increase in chromatin accessibility in CD14⁺ monocytes (**Fig. 3A, Suppl. Tables 21, 22**). Gene Ontology analysis of differentially accessible regions in CD14⁺ monocytes revealed significant enrichment of pathways related to cell adhesion, migration, phagocytosis, and inflammatory responses **(Fig. 3B, Suppl. Tables 23-25**), which are consistent with our functional data that correlate increased cytotoxic activity with enhanced migratory capacity (Fig. 1C). Notably, manual inspection showed no detectable differences in chromatin accessibility at the *S100A8 and S100A9* locus between 5-AzaC-treated samples or untreated controls on day 17 **(Suppl. Fig. S19**). ATAC-seq motif enrichment analysis revealed increased accessibility of motifs associated with developmental, plasticity (i.e., epithelial-mesenchymal transition (EMT)-like) and lineage-remodeling transcription factors, as well as additional basic helix–loop–helix factors. In contrast, motifs corresponding to canonical acute inflammatory drivers, including C/EBP family members, AP-1 components, and canonical NF-κB were reduced in accessibility (**Fig. 3C**). STRING network analysis supported a modular organization of these motif shifts (**Fig. 3D**): TFs with increased accessibility formed a SMAD-centered plasticity/lineage-remodeling network (SMAD3/4/5 coupled to SNAI2/3–ZEB1–TWIST1 and TCF/bHLH factors, with RUNX1/2 as bridging nodes), whereas motifs reduced in accessibility comprised a densely interconnected canonical inflammatory network centered on NF-κB (RELA/REL), AP-1 (FOS/JUN), and C/EBPs. Notably, this inflammatory module also included established stress/feedback constraint regulators (e.g., ATF3/JDP2 and NRF2–MAF factors, with PPARA), consistent with an adaptation-like state in which acute inflammatory enhancer accessibility is attenuated while stress-adaptive circuitry is engaged. TOBIAS digital genomic footprinting analysis showed detectable NF-κB, AP-1, and C/EBP footprints in both treated and untreated samples, as illustrated by aggregate footprint profiles for JUN and CEBPG. Footprint signals were more pronounced in untreated cells, indicating reduced occupancy of canonical inflammatory transcription factors after 5-AzaC treatment rather than complete loss of binding (**Fig. 3E**). Consistent with prior studies in high-risk MDS suggesting that 5-AzaC can preferentially reshape broader transcriptional regulatory networks rather than promoter regions at individual loci^25^, our data support a model in which 5-AzaC primarily remodels myeloid chromatin accessibility at the level of transcription factor networks.

**Fig. 3.**
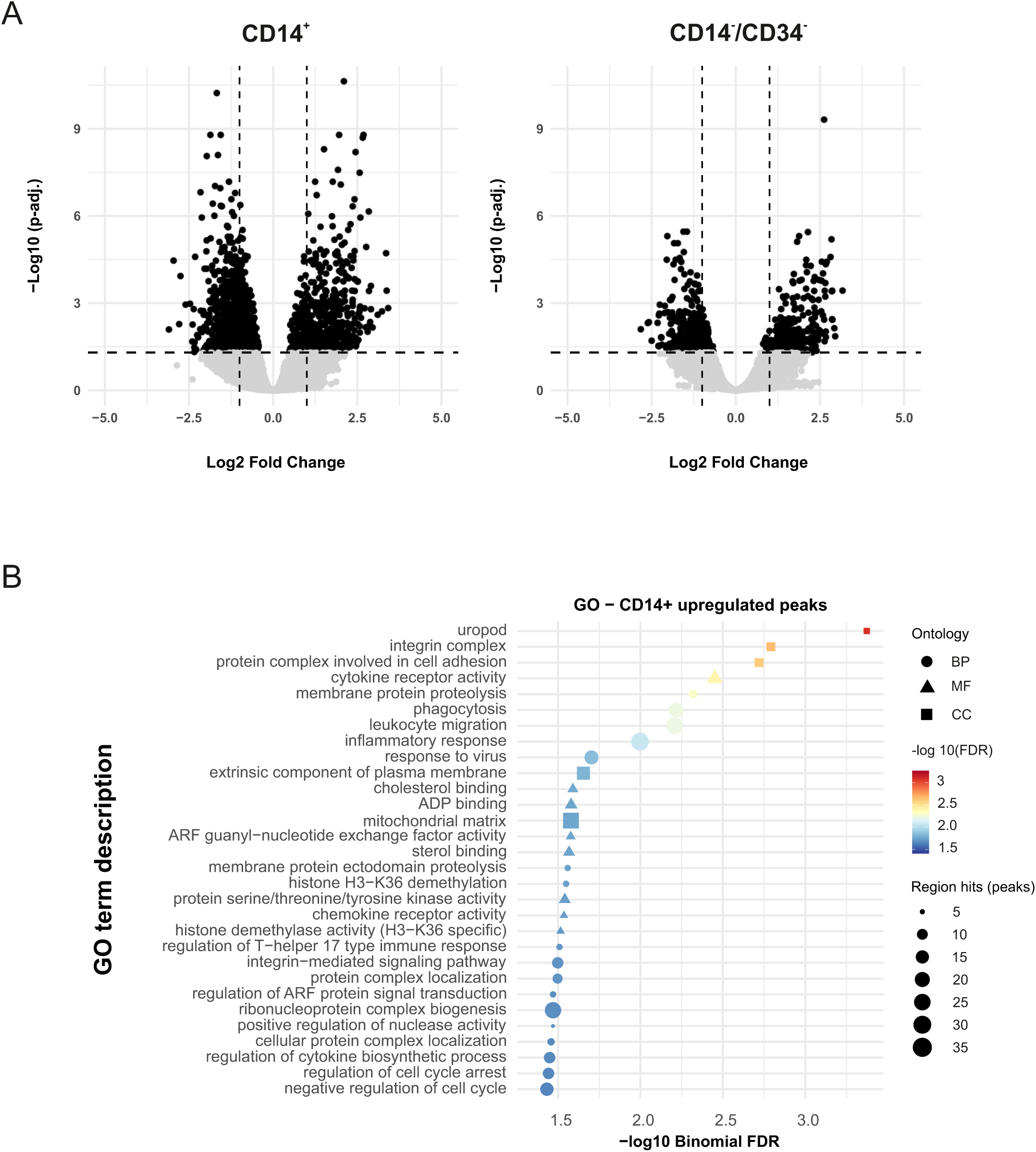
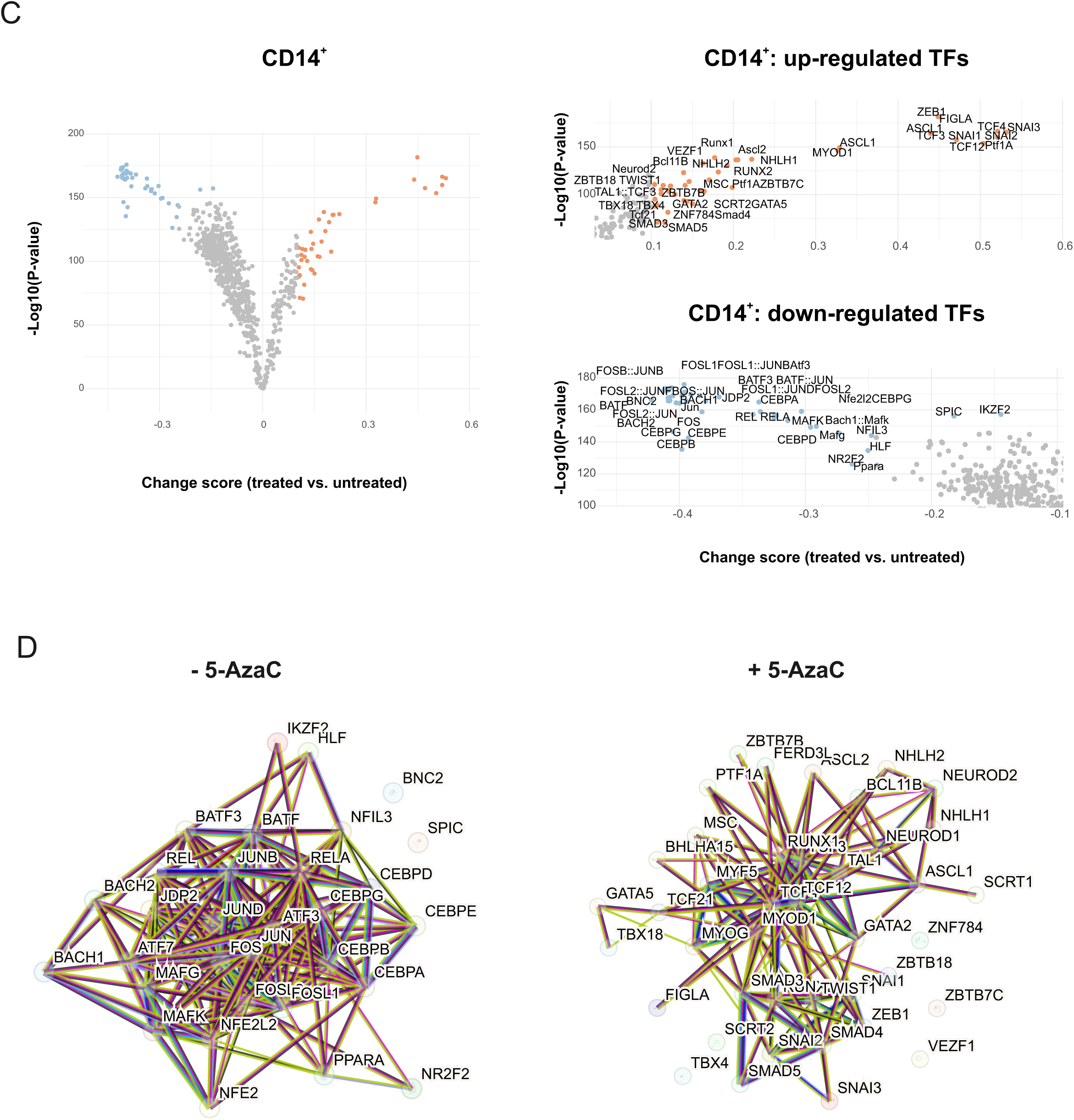
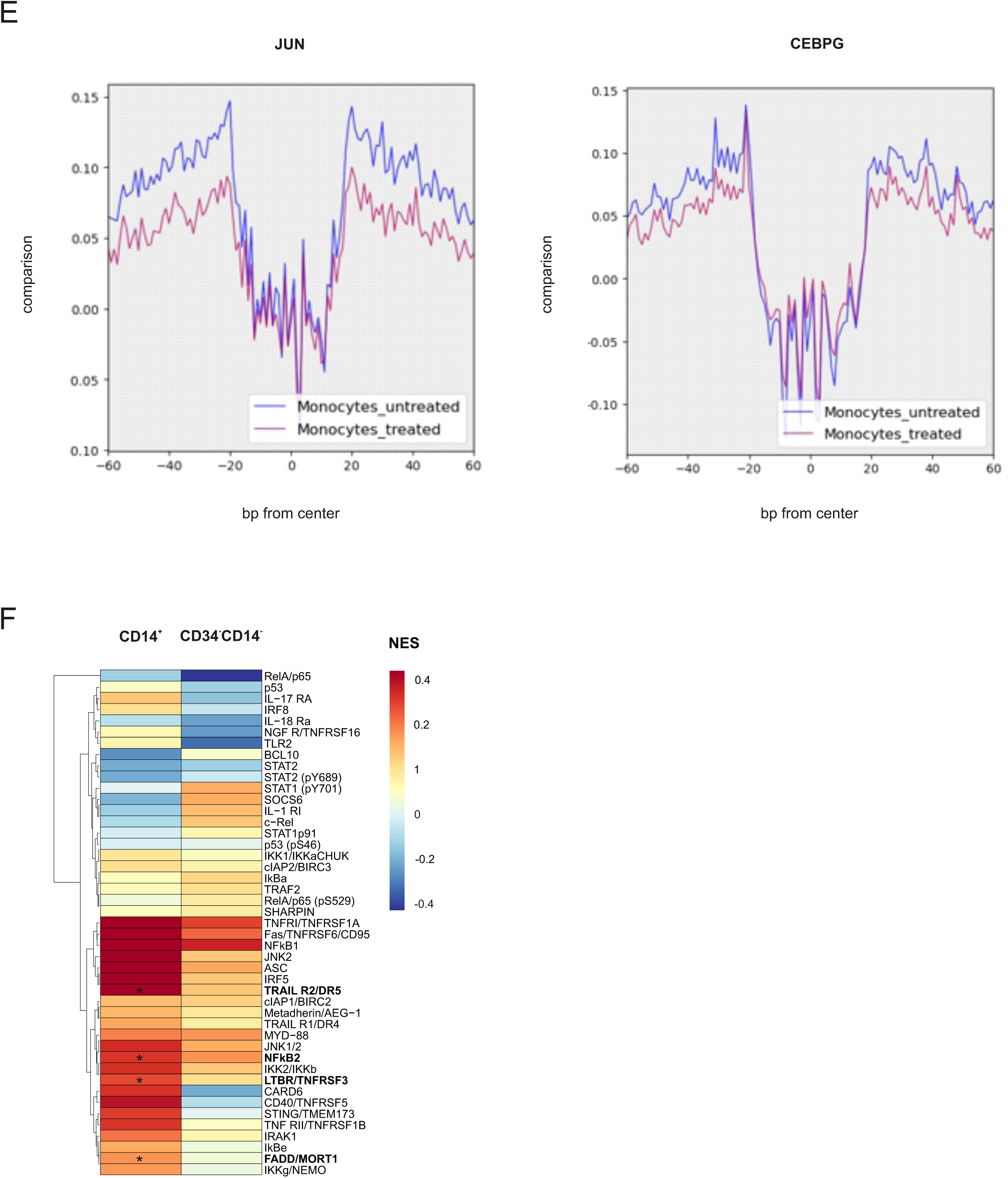
Upstream regulatory demethylation links 5-AzaC exposure to S100A8/A9 induction. **(A-E)** ATAC-seq performed on sorted day 17 CD14^+^ cell fractions or “responders”. **(A)** Volcano plot of differentially accessible chromatin peaks comparing +5-AzaC treated and untreated controls. (n = 3, significance defined as log_2_FC ≥ 1, adjusted p ≤ 0.05). **(B)** GREAT Gene Ontology enrichment analysis of differentially accessible regions in CD14⁺ cells (top 30 terms shown). Dot color indicates −log_10_(FDR) and dot size indicates peak count. Enrichment was assessed using GREAT’s region-based binomial test with basal plus extension regulatory domains. Dot color indicates −log10(Binomial FDR q value) and dot size indicates peak count; terms were filtered to Binomial FDR ≤ 0.05. **(C)** Predicted transcription factor motif binding differences inferred by TOBIAS BINDetect between treated and control conditions. Each point represents one motif (x-axis, differential binding score; y-axis, −log_10_(p value)). Statistical significance was assessed using TOBIAS’ background-model enrichment framework comparing target regions to background sequences. **(D)** STRING protein–protein association networks for transcription factors showing differential footprint activity in CD14^+^ cells inferred by TOBIAS from ATAC-seq data, shown for 5-AzaC–treated and untreated samples. Networks were generated using the full STRING network (functional and physical associations). Edges are shown in “evidence” mode, where edge color indicates the supporting evidence channel (text mining, experiments, curated databases, co-expression, neighborhood, gene fusion, co-occurrence). Associations were filtered at medium confidence (minimum interaction score = 0.400). (E) Exemplified aggregate footprints of JUN and CEBPG using TOBIAS digital genomic footprinting framework. CD14^+^ monocyte footprint scores are compared between condition 5-AzaC treated vs. untreated to define differentially bound TFs. Typical TSS enrichment plot shows that nucleosome-free fragments are enriched at SS, while mono-nucleosome fragments are depleted at TSS but enriched at flanking regions. **(F)** NF-κB pathway protein profiling in sorted CD14⁺ and CD14⁻/CD34⁻ fractions on day 21, shown as log_2_ fold change (5-AzaC/control) in “responder” cultures (n = 3 donors). Significance was assessed using two-sided one-sample t-tests comparing log_2_ fold changes against zero without correction for multiple comparisons. Significance is indicated as *p ≤ 0.05, **p ≤ 0.01, ***p ≤ 0.001. n values denote independent experiments using distinct human SC donors.

Because chromatin accessibility reflects regulatory potential rather than signaling activity, we next scrutinized inflammatory signaling pathways at somewhat later stages of differentiation. Knowing that S100A8/A9 bind to TLR4, we analyzed NF-κB–associated signaling molecules in sorted CD14⁺ and CD34^−^CD14^−^ cells on day 21 by means of protein profiling **(Fig. 3F).** Strikingly, 5-AzaC induced increased the expression of TRAILR2 (DR5), LTBR (TNFRSF3), FADD (MORT1), and NFKB2. This coordinated induction of TNF receptor family members, the adaptor protein FADD, and NFKB2 is consistent with engagement of TNFR/death receptor–linked signaling pathways and suggests a bias towards non-canonical NF-κB–associated programs at day 21. Collectively, we show that 5-AzaC treatment attenuates inflammatory monocyte characteristics and chromatin accessibility of canonical pro-inflammatory transcription factors on day 17 and delays enrichment of non-canonical inflammatory proteins at later time points in “responders”.

### Loss of S100A9 perturbs early and late hematopoietic SC differentiation and impairs functionality of immature (and mature) NK cells

We next analyzed early hematopoietic differentiation in S100A9^−/−^ mice with a targeted deletion of S100A9^26^ (**Fig. 4A**). Differential expression in bulk RNA-seq of BM mononuclear cells evidenced that next to *S100A9*, mRNA expression of *Cxcl3, Areg, Mmrn1, Snca* were significantly upregulated in WT as compared to S100A9^−/−^ mice, indicating that immune responses, inflammation, cell growth, and tissue repair are impaired in S100A9^−/−^ mice. At the same time, downregulated mRNA expression of *Pglyrp4, Cpne4, Hist1h2al, Rps3a2* and *Rps3a3* suggested a state of hyper-inflammation and cell cycle defects that may result in premature growth arrest, uncontrolled cell proliferation and cell death (**Fig. 4B, C, Suppl. Table 26**). GO term analysis in WT as compared to S100A9^−/−^ mice demonstrates upregulation of bacterial/LPS stimuli and inflammatory regulation, alongside myeloid/hematopoietic (including erythroid) differentiation and leukocyte adhesion/chemotaxis programs (**Fig. 4D, Suppl. Tables 27-29**). The unsupervised similarity analysis comparing WT vs. S100A9^−/−^ mice revealed upregulated cluster of transcripts related to acute inflammatory responses, chemotaxis/migration and interestingly also hematopoiesis/ differentiation (**Fig. 4E**). WT mice displayed a significant positive enrichment of stem- and mature-cell signatures together with a significant depletion of early/intermediate progenitor programs in comparison to S100A9^−/−^ mice in the GSEA analysis, indicating that both early and late differentiation processes are impaired in the absence of S100A9 (**Fig. 4F**). Analysis of transcription factors evidenced that members of the SMAD, NFIL3, Jun, GATA3 and CEBPB families were significantly down-regulated in S100A9^−/−^ mice (**Suppl. Fig. 20)**.

**Fig. 4.**
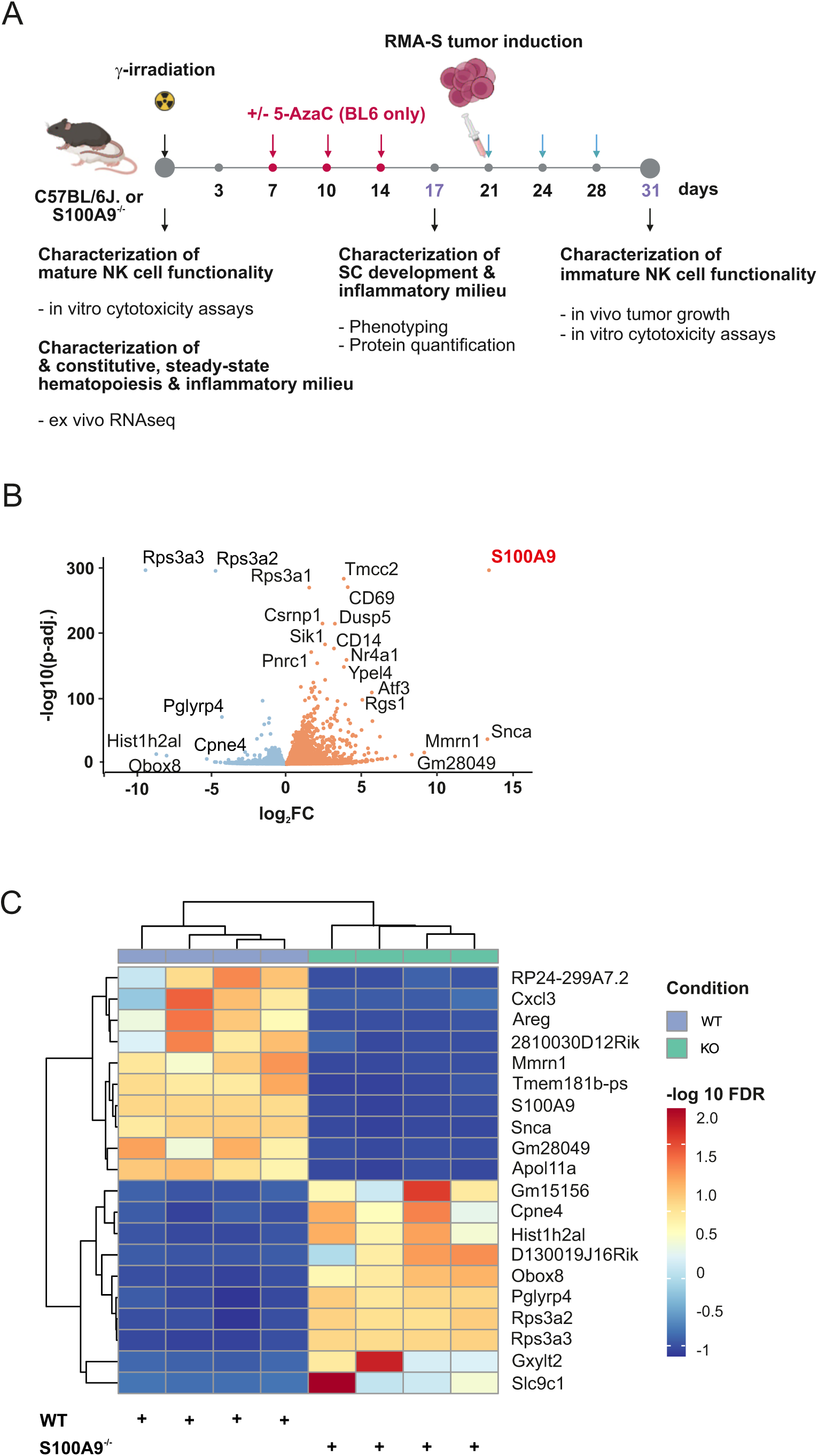
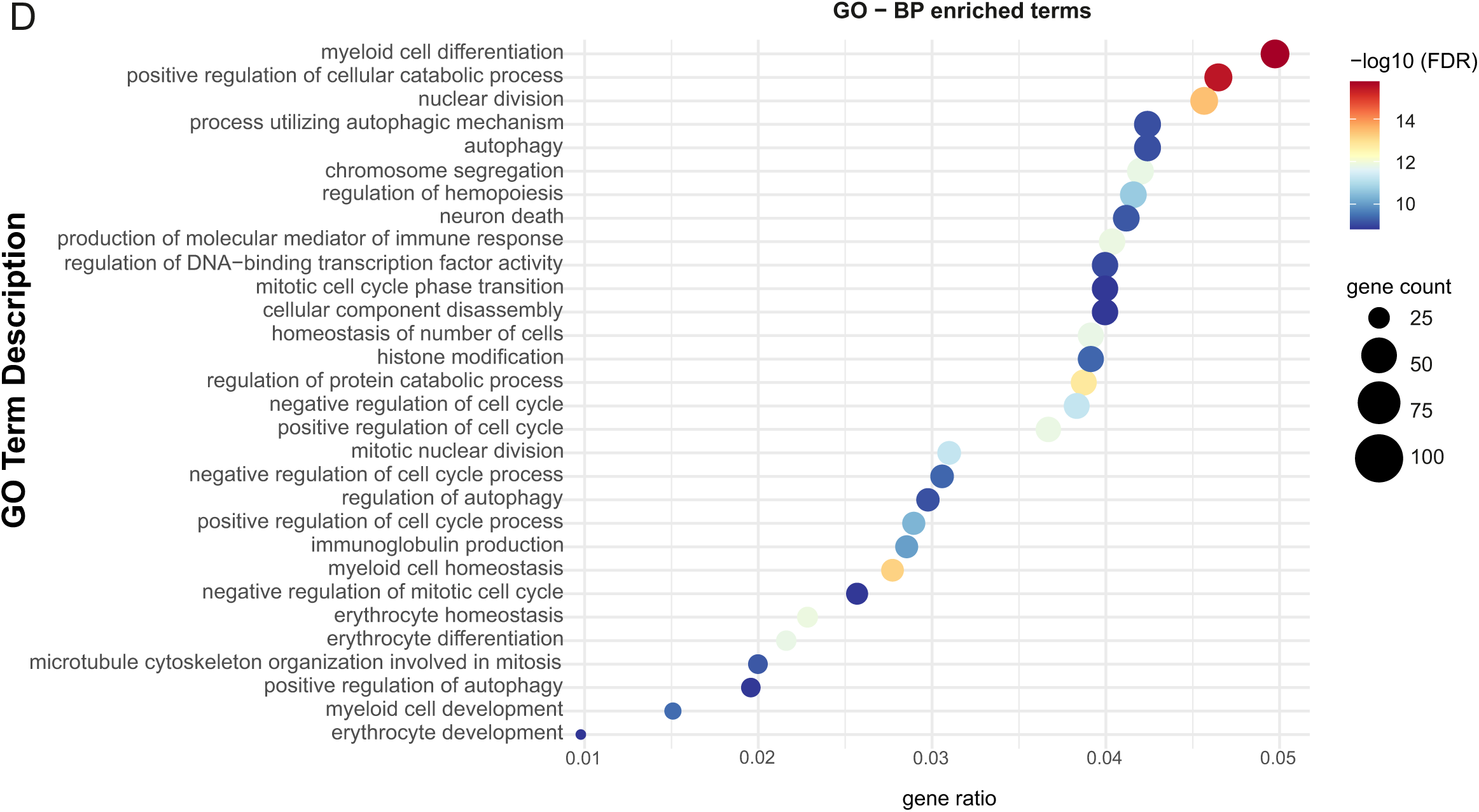
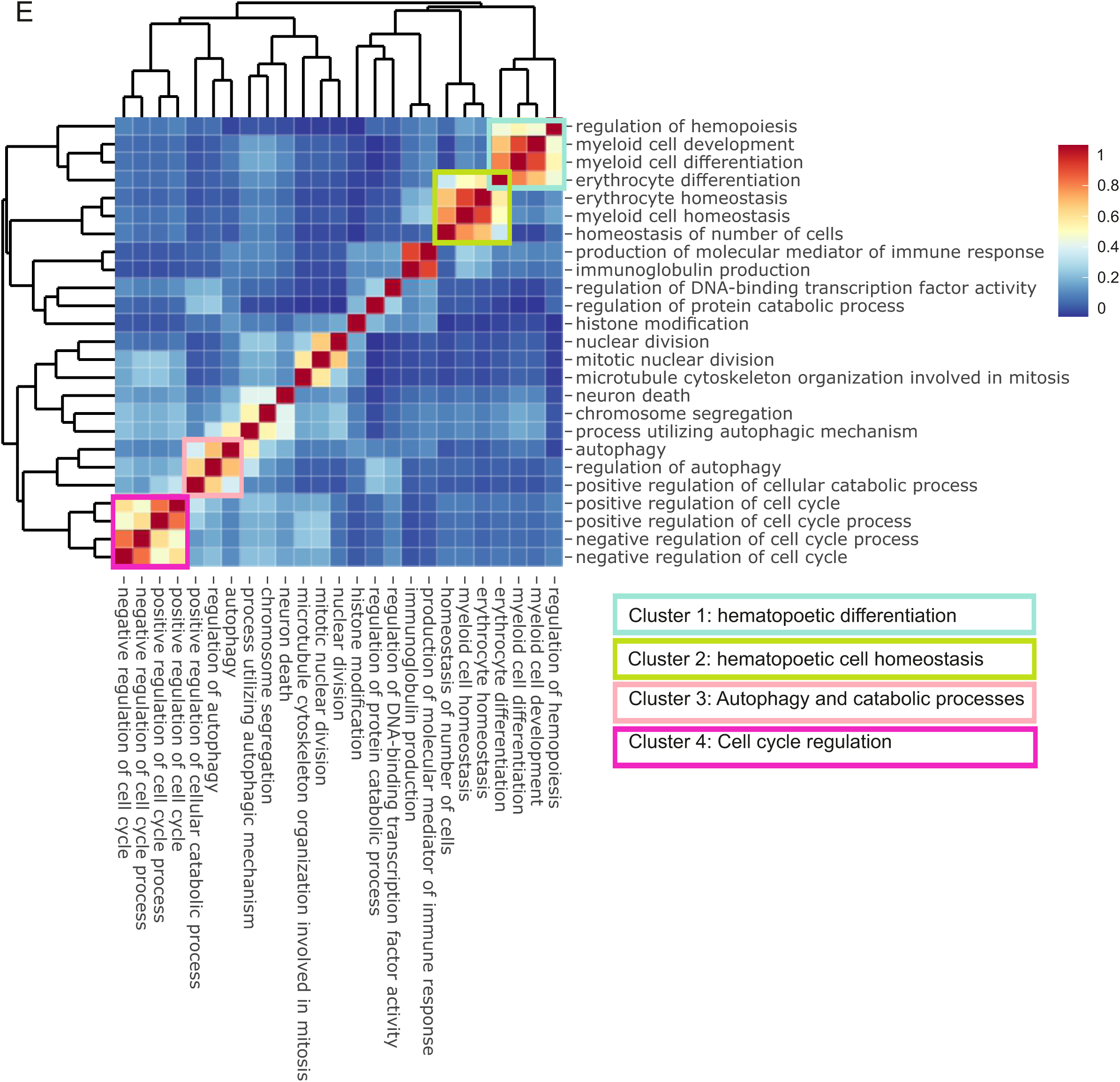
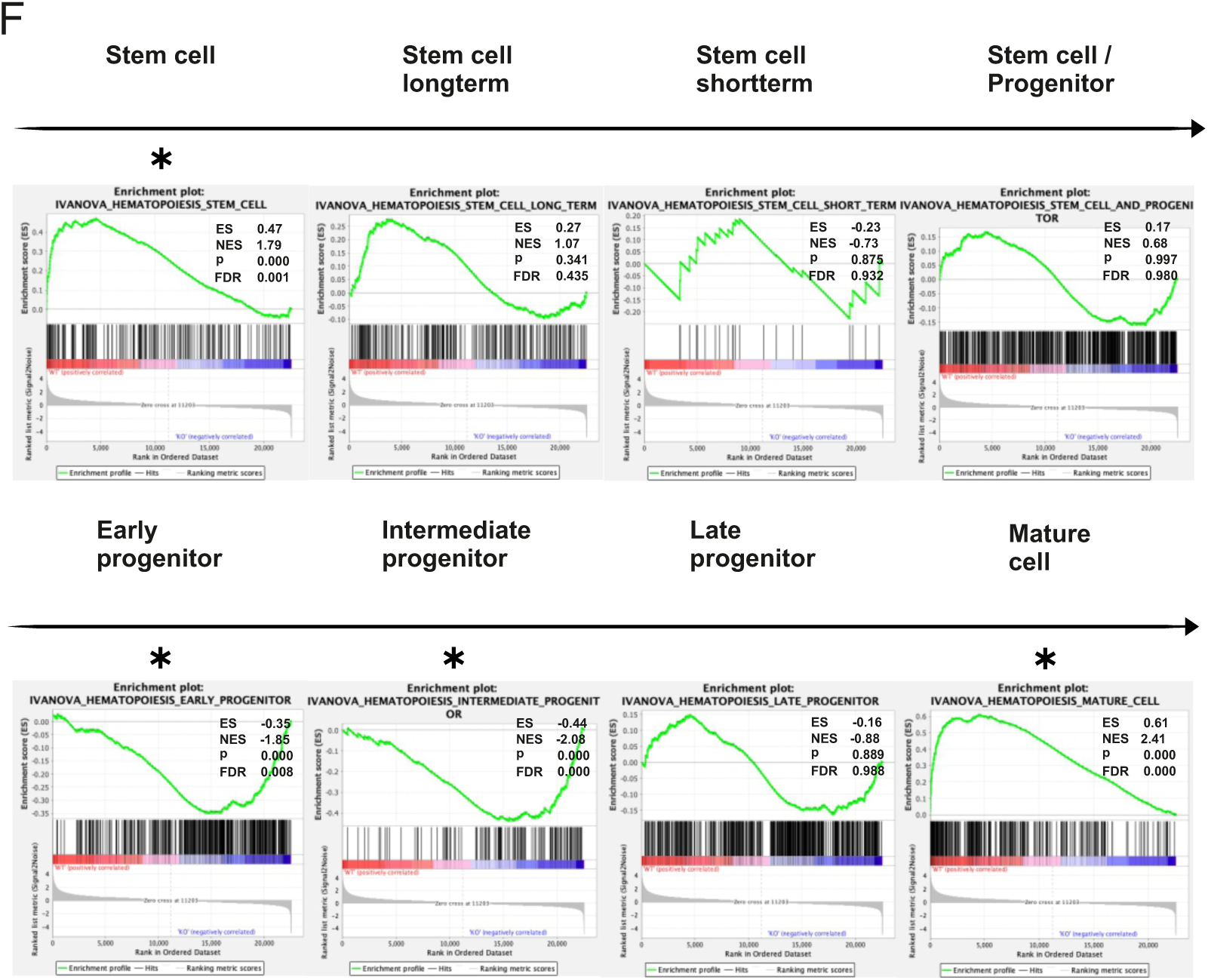
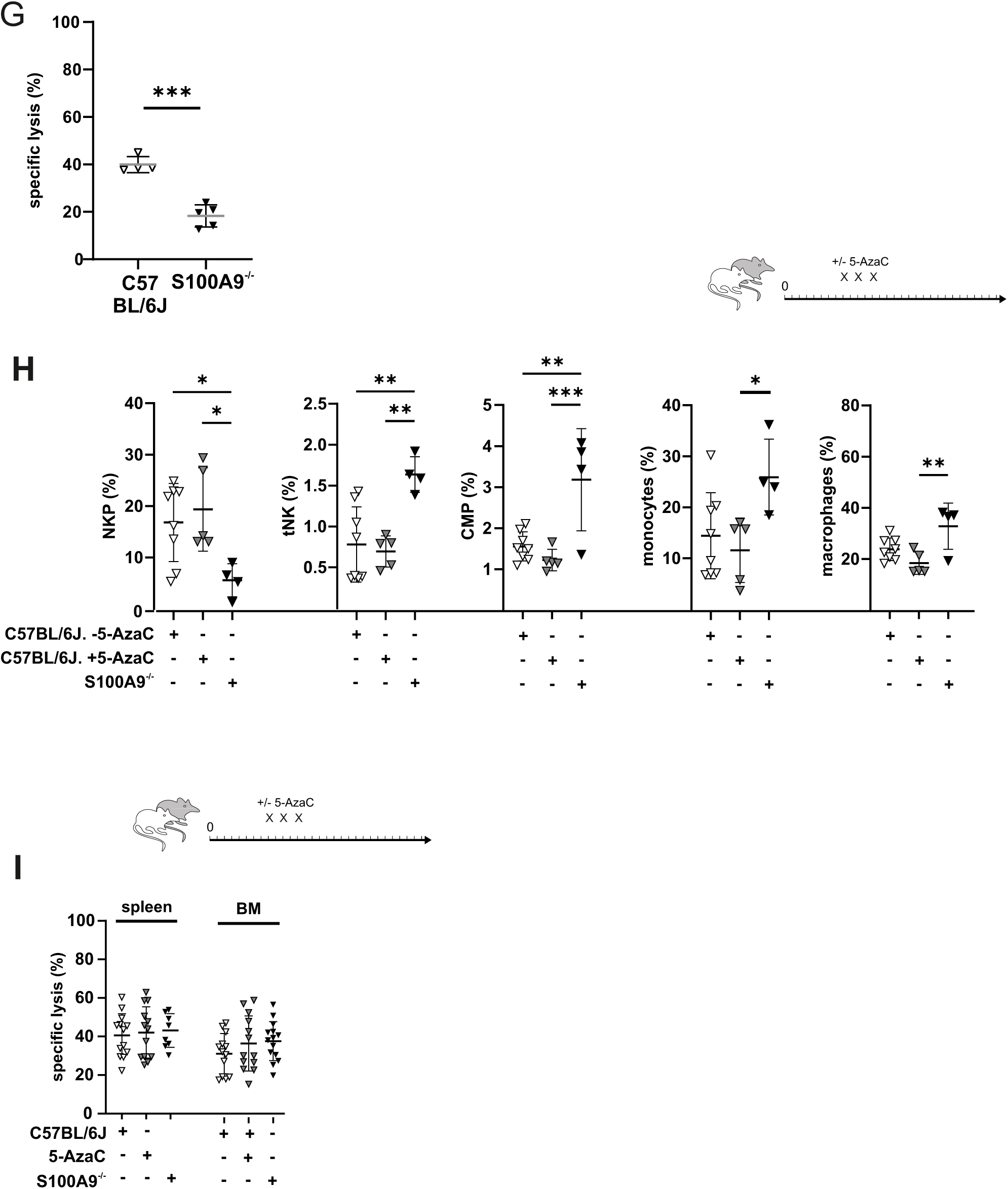
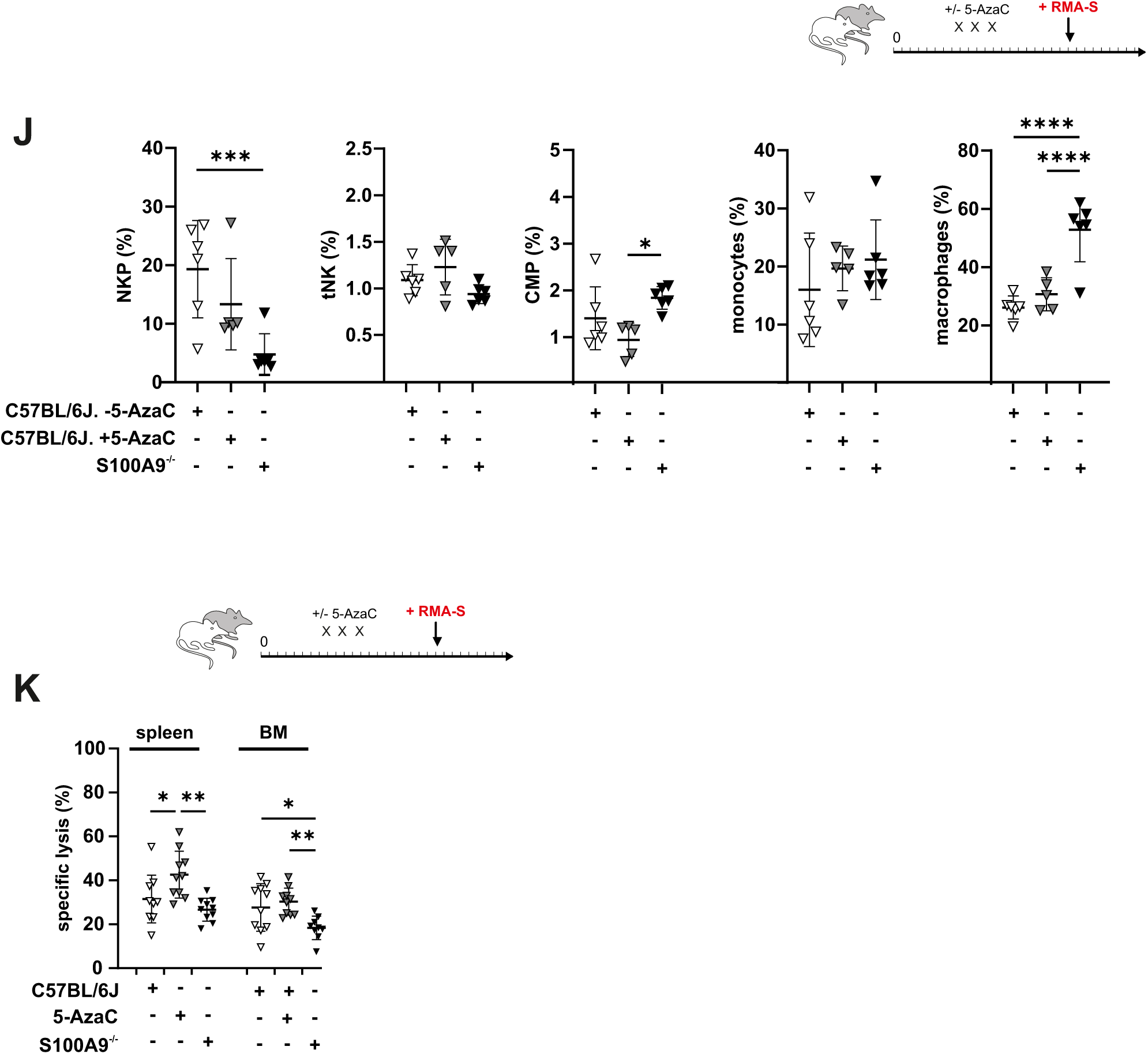
Transcriptomic profiling of C57BL/6J and S100A9^−/−^ bone marrow (BM) mononuclear cells and functional assessment of NK cell activity in vivo. **(A)** Experimental layout of animal experiments. **(B-F)** Bulk RNA-seq was performed on BM mononuclear cells from C57BL/6J. wild-type (WT) and S100A9^−/−^ mice (n = 4 per group). **(B)** Volcano plot of differential expression (WT vs. S100A9^−/−^), showing log₂ fold change versus −log₁₀(adjusted p-value/FDR). **(C)** Heatmap of the top 20 differentially expressed genes ranked by absolute log₂ fold change (row-wise Z-scores of DESeq2-normalized expression; hierarchical clustering). **(D)** Gene Ontology (GO) enrichment analysis of Biological Process (BP) terms for differentially expressed genes upregulated in WT relative to S100A9^−/−^ mice; dot color indicates −log₁₀(FDR), dot size indicates the number of overlapping genes, and gene ratio denotes overlap divided by the size of the input gene list. Enrichment significance was assessed using a one-sided hypergeometric (Fisher’s exact) over-representation test with Benjamini–Hochberg correction. **(E)** Semantic similarity analysis of GO terms comparing WT and S100A9^−/−^ conditions. **(F)** Gene set enrichment analysis (GSEA) using IVANOVA hematopoiesis gene sets; enrichment is reported as enrichment score (ES)/normalized enrichment score (NES) with corresponding p-values and false discovery rate (FDR). Significance was assessed using 1.000 size-matched gene set permutations. (G) *Ex vivo* cytotoxicity of splenic mature NK cells against the RMA-S cell line (E:T 100:1, unpaired t-test). **(H)** Flow cytometric analysis of hematopoietic reconstitution in sublethally irradiated WT mice treated ±5-AzaC and untreated S100A9^−/−^ mice, assessed on day 28–33 post irradiation (n = 4-8; two-way ANOVA with Šídák’s multiple-comparisons test). (I) *Ex vivo* cytotoxicity of immature NK cells isolated from spleen or bone marrow against RMA-S targets (E:T 100:1) in unprimed mice (n=13-15; one-way ANOVA with Tukey’s multiple-comparisons test). **(J)** Hematopoietic reconstitution in irradiated WT ±5-AzaC and S100A9^−/−^ mice following in vivo NK cell priming by subcutaneous RMA-S tumor inoculation (n = 5-6; two-way ANOVA with Šídák’s multiple-comparisons test). (K) *Ex vivo* cytotoxicity of immature NK cells obtained from spleen or bone marrow tested against the RMA-S cell line (E:T 100:1) on day 28-33 post sublethal irradiation. NKP, NK precursor; tNK, transitional NK cells; CMP, common myeloid progenitor. (n = 9-10; one-way ANOVA with Tukey’s multiple-comparisons test). Significance is indicated as *p ≤ 0.05, **p ≤ 0.01, ***p ≤ 0.001, ****p ≤ 0.0001. n values denote independent experiments using distinct number of mice.

Based on this, we tested whether our in vitro observations on healthy human developing SCs could be recapitulated in mice. As an initial functional baseline testing, we quantified ex vivo cytotoxicity of splenic NK cells of naïve S100A9^−/−^ or WT mice and demonstrated a significant functional impairment of mature NK cells in S100A9^−/−^ mice (**Fig. 4G**). We then studied hematopoietic reconstitution in sublethally irradiated +/- 5-AzaC treated C57BL/6J wild-type (WT) mice in comparison to untreated S100A9^−/−^ mice (**Fig. 4H, Suppl. Fig. S21A, B, Suppl. Fig. S22A**). Confirming our findings in the GSEA (Fig. 4E), S100A9^−/−^ mice lacked physiological numbers of NK cell precursors (NKP) but displayed clearly increased numbers of transitional CD49b^−^/NK1.1^+^ NK cells (tNK cells), common myeloid progenitors (CMP), monocytes but also macrophages around day 28-33 post-irradiation. Contrary to our expectations, ex vivo testing of immature NK cell function against the MHC class I-deficient RMA-S tumor cells did not show any differences between WT or S100A9^−/−^ mice and did not show any effect of 5-AzaC treatment (**Fig. 4I**) not even when injecting a complex of IL15 and IL15R in vivo (data not shown).

Knowing that the acquisition of in vivo functionality in immature NK cells requires complex conditions consisting of a first cytokine signal (i.e., complex of IL15 and IL15R), a second signal through metabolic re-reprogramming (improved glycolysis and mitochondrial activity) and a third signal via structural crosstalk priming with dendritic cells (CD62L-mediated homing to lymph nodes)^27^, we decided to subcutaneously (s.c.) inoculate our mice with RMA-S tumor cells to provide sufficiently instructive signals to the immature NK cells. Under these experimental conditions, we observed more pronounced quantitative but not qualitative differences between lineage differentiation in C57BL/6J WT and S100A9^−/−^ mice (**Fig. 4J, Suppl. Fig. S22B**). In line with increased concentrations of serum concentrations of S100A8/A9 in 5-AzaC treated tumor-bearing C57BL/6J mice (**Suppl. Fig. S23**), ex vivo testing of immature NK cell functionality now demonstrated a clear-cut positive effect of in vivo 5-AzaC treatment on immature NK cell functionality (**Fig. 4K**). To exclude the possibility that in vivo priming with RMA-S had elicited an adaptive T cell-mediated immune response, we repeated all cytotoxicity assays with prior CD8 T cell depletion (**Suppl. Fig. S24**). Taken together, these findings support a role for S100A9 in both early and late hematopoietic SC differentiation and document that S100A9 is required during the acquisition of functionality of immature (and mature) NK cells.

### Clinical response to 5-AzaC is associated with skewed myeloid differentiation and signs of acute inflammatory response in MDS/CMML patients

To link our findings to clinically relevant patient outcomes, we next asked whether S100A8/A9 expression was also associated with clinical outcomes in 5-AzaC-treated patients. To this end, we analyzed transcriptomic data from a well-annotated Australian MDS/CMML cohort^28^ (**Fig. 5A**) and compared the ability to elicit an inflammatory response in clinical 5-AzaC “responders” and “non-responders” as defined by IWG 2006 criteria^29^. We first verified that the cohort included patients with a meaningful difference in clinical outcome by comparing overall survival of “responders” and “non-responders” (**Fig. 5B**). We then interrogated CD34⁺ stem/progenitor cell transcriptomes to determine whether “response” was associated with induction of S100A8/A9 and enrichment of innate immune and myeloid differentiation programs. Indeed, S100A8 and S100A9 were significantly upregulated in “responders” following six consecutive cycles of 5-AzaC (**Fig. 5C**). In line with our findings in CD34^+^ SCs of healthy donors (Fig. 1E), MDS “responders” exhibited enrichment of GO biological process (BP) terms related to immune activation, cell surface receptor signaling, leukocyte migration, phagocytosis (**Fig. 5D**), and extracellular matrix and secretory granule lumen (**Suppl. Fig. S25A, B**). Consistently, Hallmark gene sets for inflammatory response, interferon gamma response, IL6-JAK-STAT3 signaling, and TNF alpha signaling were highly enriched in “responders” following 5-AzaC treatment (**Fig. 5E**). Digital cytometry applying CIBERSORTx revealed clear-cut differences between MDS/CMML “responders” and “non-responders” in the myeloid progenitor compartment (**Fig. 5F**). Interestingly, differences between “responders” and “non-responders” in these myeloid compartments (GMP, MEP and MPP) but also in gene sets coding for inflammatory pathways such as Myc targets and E2F targets (**Suppl. Fig. S26**) were in fact inherently demonstrable before initiation of 5-AzaC. GSEA revealed that only MDS/CMML “responders” displayed signs of enhanced myeloid maturation, with reduced stemness-associated signatures and enrichment of early progenitor and mature gene sets (**Fig. 5G**). Consistent with our hypothesis that one aspect of “response” to 5-AzaC is based on the propensity to elicit myeloid skewing and an S100A8/A9–TLR4-mediated inflammatory response, we found in MDS patients that the shared genes mapped to monocyte/macrophage lineage and differentiation programs (**Fig. 5H**) and the shared BP terms (**Fig. 5I**) were related to innate effector functions, including phagocytosis, leukocyte adhesion/recognition, and oxidative burst/reactive oxygen species (ROS) metabolism (**Suppl. Tables 32-34**).

**Fig. 5.**
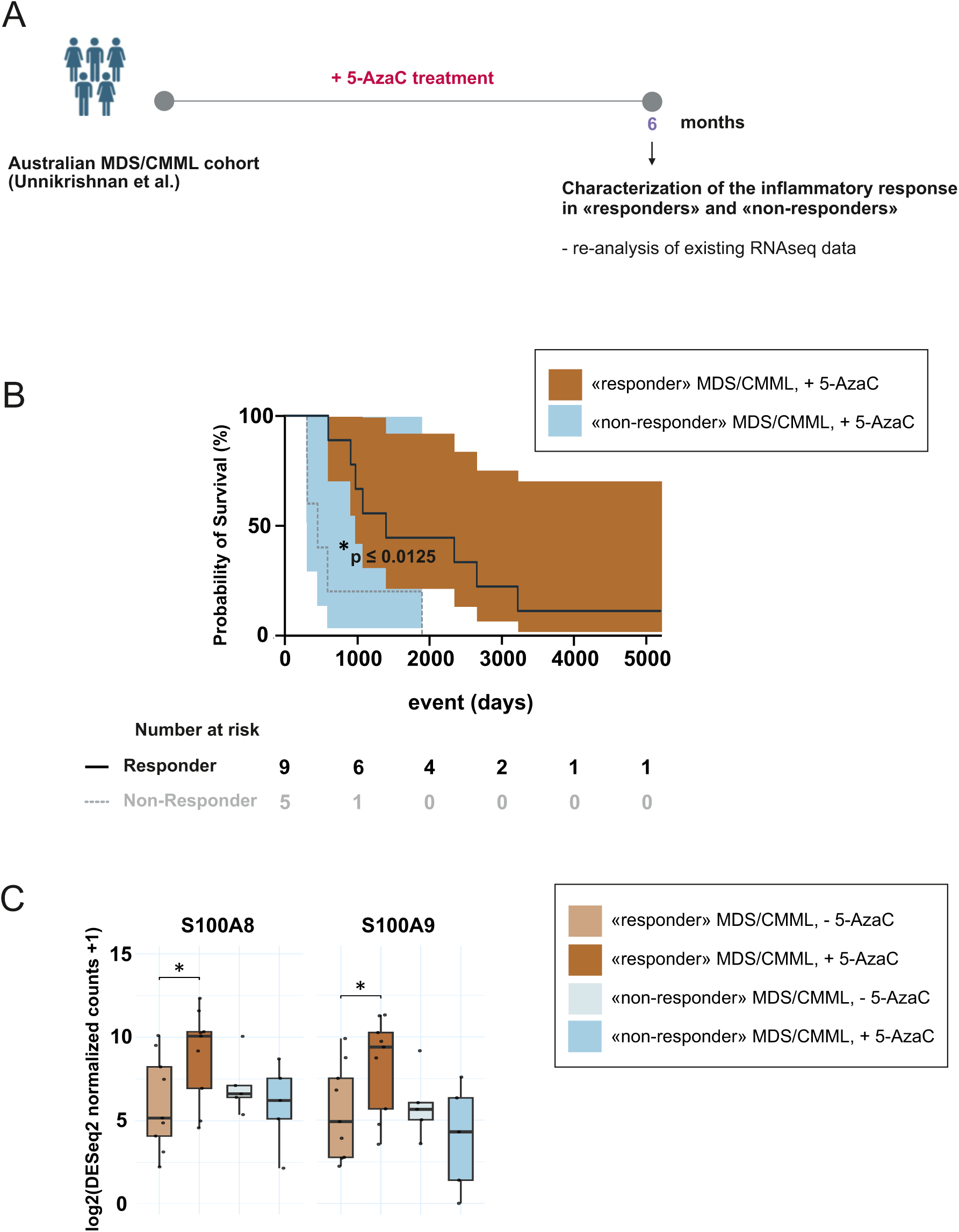
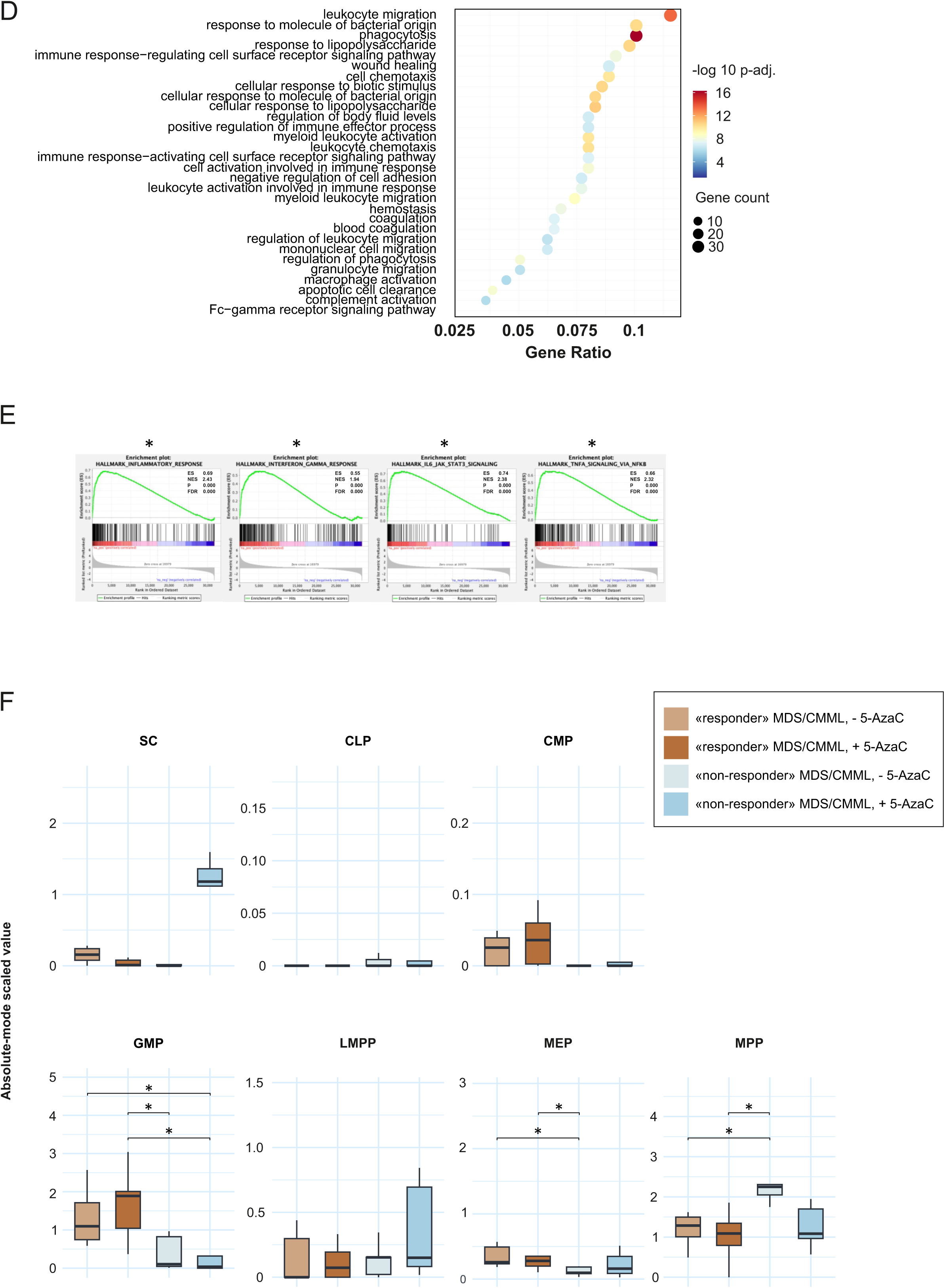
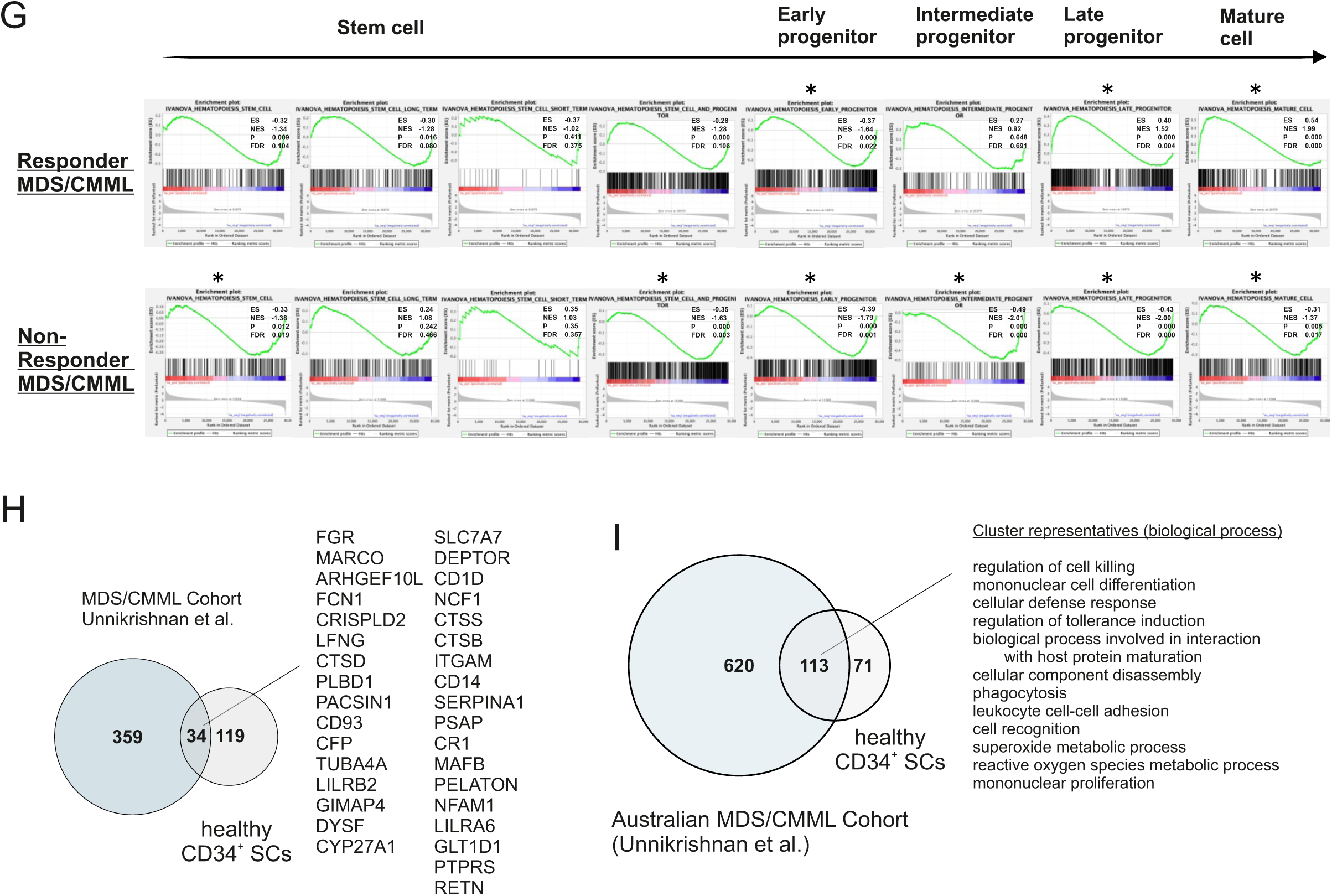
Transcriptional inflammatory signatures associated with clinical response to 5-AzaC in MDS/CMML CD34⁺ cells from the Unnikrishnan cohort. **(A)** Overview of study characteristics. **(B-I)** BulkRNA-seq data were obtained from the Unnikrishnan et al. study (GSE76203) together with additional datasets provided by the same authors. CD34⁺ cells from patients with MDS/CMML were profiled before treatment and after six cycles of 5-AzaC. The cohort included 9 “responders” and 5 “non-responders” (according to IWG 2006 criteria) (Cheson et al.). **(B)** Kaplan–Meier analysis of overall survival stratified by clinical response to 5-AzaC (Mantel–Cox log-rank test, p ≤ 0.0125). **(C)** Box-plot of S100A8 and S100A9 expression grouped by “response” and treatment. Treated vs. untreated comparisons were done using log-transformed DESeq2 size-factor–normalized counts (two-sided) paired t-test was applied for treated/untread comparison (*p value < 0.05). **(D)** Bubble plot of the top 30 enriched GO Biological Process (BP) terms among genes upregulated in treated vs. untreated responders. Over-representation was tested using a one-sided Fisher’s exact (hypergeometric) test with Benjamini–Hochberg correction; dot size indicates gene count, color indicates −log10(p-adj.), and the x-axis shows gene ratio. **(E)** GSEA enrichment plots for selected Hallmark pathways enriched in treated vs. untreated responders. Pre-ranked GSEA was performed using MSigDB Hallmark gene sets with genes ranked by the DESeq2 Wald statistic; enrichment is reported as NES with nominal p and FDR (significance was assessed using 10.000 permutation). **(F)** Boxplots of estimated progenitor cell-type abundances from digital cytometry (CIBERSORTx). Paired treated vs. untreated comparisons within “responders” and “non-responders” used (two-sided) Wilcoxon signed-rank tests; all other between-group comparisons used (two-sided) Wilcoxon rank-sum (Mann–Whitney U) tests. p-values were BH-adjusted across six pairwise comparisons per cell type. **(G)** Pre-ranked GSEA using IVANOVA hematopoietic reference gene sets (analysis performed as in D), indicating enrichment of progenitor- and mature-cell–associated transcriptional programs. **(H, I)** Venn diagrams showing overlap in **(H)** differentially expressed genes and **(I)** enriched GO biological processes (BP) between 5-AzaC “responsive” MDS/CMML CD34⁺ cells and healthy differentiating d17 CD34⁺ cultures.

## Discussion

5-AzaC is used in the posttransplant setting as an off-label prophylactic maintenance treatment strategy for MDS/CMML patients with a high risk of relapse^30^. However, only around 50% of patients show a clinically significant “response” with improved survival outcomes and decreased likelihood of leukemic transformation^31,32^. We here provide compelling evidence that inherent inflammatory properties of healthy donor CD34^+^ SCs exist that are likely to contribute to the “response” seen upon pre-emptive posttransplant 5-AzaC therapy of high-risk MDS patients. We describe distinct “responder” and “non-responder” immune phenotypes of healthy donor SCs that mainly consist of a so-called inflammatory monocytic cluster (inflMo) in “responders”, and intermediate (IntMo) and hyperinflammatory (hyinflMo) monocytic clusters in “non-responders”. The inflMo cluster displays an inflammatory instructive phenotype that strengthens migration/adhesion and DAMP (S100A8/A9–TLR4)-related ligand-receptor interactions in “responders”, whereas the IntMo and hyinflMo clusters represent metabolically and structurally reprogrammed, non-instructive states in “non-responders” that engage primarily in MHC class I (B2M or HLA-B)-CD94 (KLRD1))–mediated interactions. Collectively, these phenotypes are defined by differences in their ability to link epigenetic re-programming of the myeloid niche in the context of S100A8/A9-driven inflammation with acquisition of immature NK cell functionality (**Fig. 6**).

**Fig. 6.**
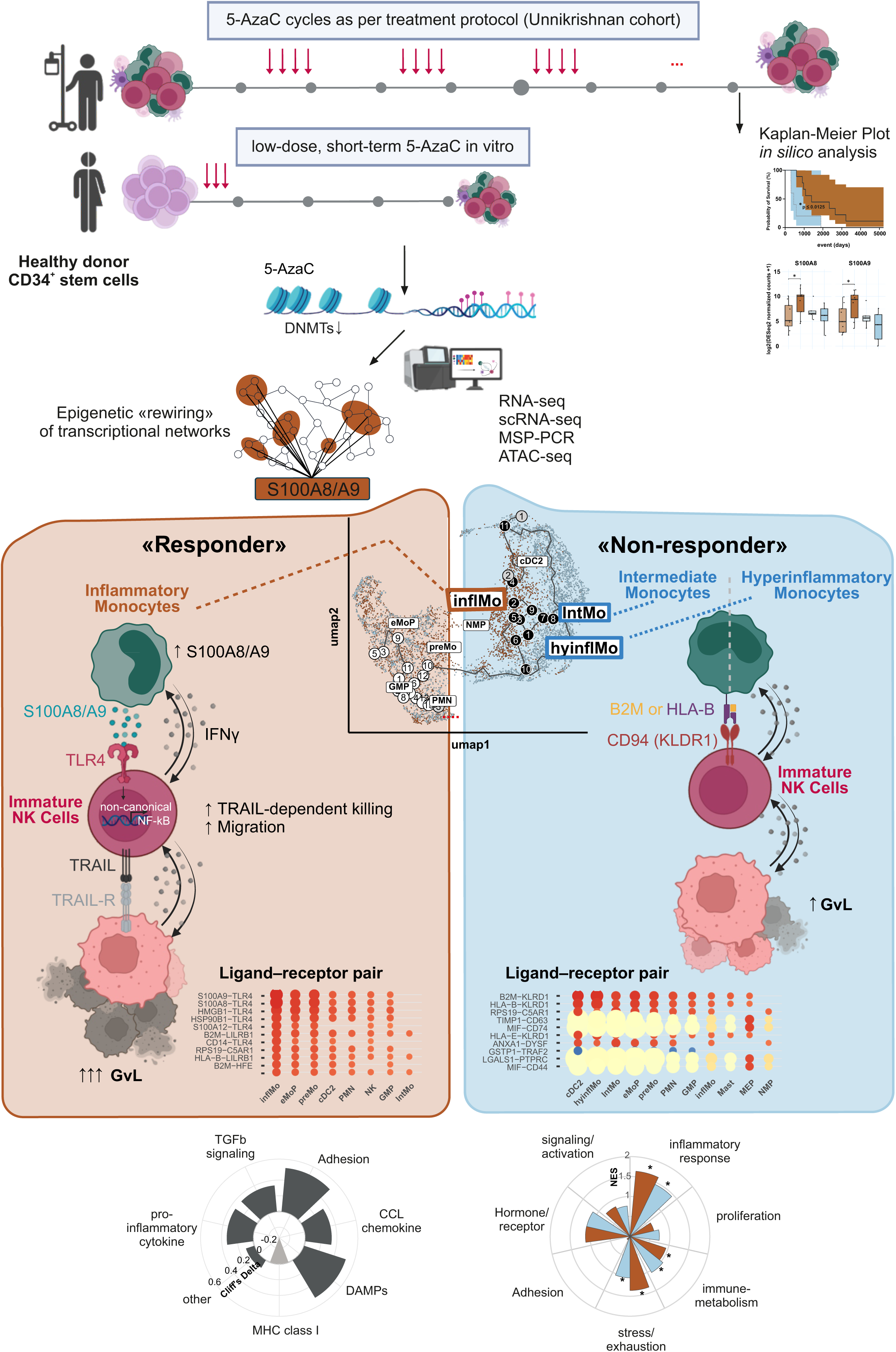
Proposed model. Linking the epigenetic control of inflammation to NK cell-mediated Graft-versus leukemia effects. Shown is the putative interaction between the two key players, inflammatory monocytes and immature NK cells in the context of 5-AzaC exposure.

All-in-all, little is known when and how immature NK cells may be primed to elicit GvL effects, but it is assumed that they rather orchestrate the immune response, than directly exert cytotoxic function. Our findings indicate that the inherent differences in the inflammatory properties of CD34^+^ donor SCs also encompass differences in the NK cell compartment as 5-AzaC-treated immature NK cells of “responders” display a more mature and activated phenotype together with transcriptional programs that are consistent with enhanced cytotoxic readiness and metabolic activation, whereas immature NK cells of “non-responders” express signs of a vivid cell cycling but not of effector cell maturation. In line with previously published data describing that immature NK cells acquire increasing motility as they mature^33^, we document in our human bulk RNA-seq, scRNAseq, ATAC-seq and murine bulk-RNAseq but also in migration assays using human immature NK cells that the ability for migration towards tumor cells is ameliorated upon 5-AzaC-induced up-regulation of S100A8/A9 and is impaired in the absence of S100A9. Lastly, we document that immature human “responder” NK cells are activated when the inflammatory protein S100A8/A9 binds to the TLR4 receptor and elicits non-canonical NF-κB signaling, ultimately leading to what is presumed TRAIL-TRAILR-mediated cytotoxicity.

Notably, this process bypasses the RAGE receptor, which is typically a common binding partner for S100A8/A9. This contrasts with the few existing publications, showing in C57BL/6J mice that the S100A8/A9 complex may have a direct role in mature NK cell activation by eliciting RAGE-p38/MAPK/NF-κB signaling and co-stimulating NKG2D ligand (NKG2DL)-mediated induction of IFN-γ synthesis^34,35^. The molecular determinants that govern the preference for one effector receptor over another remain an unresolved question but could be explained by the quaternary structure of the S100A8/A9 as dimers (and not tetramers) that favor TLR4 over RAGE, by low Ca2^+^ levels as these rather induce dimerization, and by the binding affinity of S100A8/A9 itself which is higher towards TLR4.

To our knowledge, this study is the first to document that low-dose 5-AzaC reprograms healthy donor SCs and subsequently modulates their lineage commitment in the context of S100A8/A9-orchestrated inflammation. The molecular mechanisms that underlie the transcriptional regulation of S100A8/A9 genes remain poorly understood. Our MSP-PCR and ATAC-seq analysis reveal that up-regulation of S100A8/A9 is not a direct consequence of local chromatin opening but rather induced by epigenetic plasticity of upstream-located transcriptional factors that sustain a S100A8/A9-driven inflammation. This process is demonstrable in distinct monocyte subsets but also in NK cells and ultimately determines the specifics of inflammatory monocyte-NK cell crosstalk. Mirroring our findings in inflammatory CD14^+^ monocytes, the upregulation of S100A8/A9 in myeloid-derived suppressor cells (MDSCs) results from a global reconfiguration of transcription factor (TF) networks, rather than a localized increase in chromatin accessibility at the S100A8/A9 locus^36^.

Modern epigenomics suggests that 5-AzaC modulates the myeloid landscape in two ways. First, it may directly inhibit DNA methyltransferases (DNMTs) which demethylate CpG islands in specific promoter regions (e.g., of IL-6, TNF, or IFN genes) or may induce endogenous retroviruses (ERVs) that can elicit a potent innate immune reaction. Second, 5-AzaC may act systemically in the sense of remodeling TF networks by opening distal enhancers and by unmasking binding sites thereby exposing motifs for lineage-determining TFs such as PU.1, CEBP and AP-1. We interpret the data derived from healthy donor SCs within the framework of “systemic network rewiring” that resulted in a coordinated shift in the myeloid cell’s identity and an enhanced inflammatory S100A8/A9-mediated “readiness” that licensed immature NK cell functionality and ultimately controlled target cell recognition.

Our ex vivo analysis of mice with a targeted deletion of S100A9 demonstrates that S100A9 deficiency disrupts the temporal kinetics of healthy hematopoietic SC differentiation. Bulk RNA-seq suggests that this perturbation is characterized by a failure to properly upregulate the myeloid-priming factors CEBPB and Jun during early stem cell commitment. Consequently, the developmental delay extends into late-stage maturation, where a significant reduction in NFIL3 expression halts the acquisition of functionality of both immature and mature NK cells. Crucially, we establish a causal link between these proteins and the observed pathology by showing that pharmacological inhibition of S100A8/A9 with Tasquinimod phenocopies these perturbing effects, thereby confirming that the S100A8/A9 axis is the primary driver of the hematopoietic disruption. Moreover, our in vivo experiments in sublethally irradiated C57BL/6J mice demonstrate that 5-AzaC treatment effectively restores dysregulated hematopoietic homeostasis and improves NK cell functionality.

In MDS patients, the Pimanda group recently linked 5-AzaC *response” with enrichment of immune-related pathways and identified “non-responders” based on their enrichment in extracellular matrix and erythrocyte associated pathways^37^. The same group described that only “responders” were able to elicit an inflammatory response with activation of the complement system, pattern-recognition receptor signaling, ROS production and pro-inflammatory cytokine and NK cell signaling^28^. Our in silico analysis of the Australian MDS/CMML cohort confirms and extends these data by providing evidence that significant inherent differences in the precursor architecture of “responder” and “non-responder” exist that are reflected by different cluster sizes of GMP and MEPs but also by diverging inflammatory immune phenotypes, particularly of Myc and E2F targets. Consistent with our findings in healthy CD34^+^ SCs, we also found upregulated S100A8A/9 mRNA transcripts and signs of interferon gamma, IL6-JAK-STAT3, and TNF alpha signaling in the Australian MDS/CMML cohort of “responders”.

The strengths of our study derived from an extensive epigenetic analysis of healthy differentiating CD34^+^ SCs which permitted a detailed characterization of the functional crosstalk between inflammatory monocytes and immature NK cells that ultimately determines GvL effects. Yet the biological implications of these findings need to be considered alongside certain methodological constraints. Because the scRNA-seq cohort was limited in size due to high costs, statistical power was restricted, especially for analyses of small cell populations. Although we applied pseudobulk DESeq2-based differential expression analysis where feasible, not all comparisons were amenable to pseudobulking and some analyses instead relied on single-cell level models, which are more sensitive to limited cell numbers and sample-to-sample variability. Nevertheless, the high-dimensional nature of our technique and the multi-omics approach provided a detailed landscape of healthy “responder” and “non-responder” donor CD34^+^ SC differentiation that was previously unavailable. While our findings suggest S100A8/A9-TLR4 signaling as a one key regulator of eliciting immature NK cell function, further studies, ideally using targeted demethylation of enhancer regions by introducing a dCas9-Tet1 construct into CD34^+^ hematopoietic stem cells^38–40^ are necessary to provide the missing causal link.

In conclusion, our data provide a valuable framework to guide future development of targeted epigenetic niche therapy in myeloid malignancies as we provide compelling evidence that inherent properties of healthy donor SCs exist that might contribute to a “response” of pre-emptive posttransplant 5-AzaC therapy of high-risk MDS patients. As such, screening of healthy donors SCs for specific inflammatory markers in early developing monocytes could be used as a marker to predict which donor will have the potential of generating a S100A8/A9-driven inflammatory response and to identify MDS but theoretically also AML recipients that might benefit from a low-dose, short-term 5-AzaC therapy as early as day 7 post transplantation.

## Supporting information

Statistical Tests Overview

## Abbreviations

AML: acute myeloid leukemia
5-AzaC: 5-Azacytidine
ATAC-seq: Assay for Transposase-Accessible Chromatin with sequencing
BM: bone marrow
DAMPs: Damage-associated molecular patterns
DNMT: DNA methyltransferase
FDR: False discovery rate
GSEA: Gene set enrichment analysis
GvL: Graft-versus-Leukemia
huNSG: humanized NOD.Cg-*Prkdc^scid^ IL2rg^tmWjl^*/Sz mice
HLA: Human leukocyte antigen
HSCT: Hematopoietic stem cell transplantation
IFN: Interferon
KIR: Killler immunoglobulin-like receptor
MRD: Minimal residual disease
NK cells: Natural Killer cells
PAMPs: pathogen-associated molecular patterns
RAGE: Receptor for Advanced Glycation Endproducts
SC: stem cells
TF: transcription factor
TSS: transcription starting site

## Acknowledgements

We thank S. Bühler, Andre lab, University of Tuebingen, Germany, for yearlong excellent technical assistance. We thank all stem cell donors for allowing us to use a defined number of stem cells for these experiments. We thank Dr. Unnikrishnan, Sydney, Australia, for providing access to the clinical and RNA-seq data from the patients included in his MDS/CMML study^28^ (Unnikrishnan et al.). We thank Prof. M. Lübbert, University of Freiburg, Germany, and Prof. Dr. K. Welte, University of Tuebingen, Germany, for their helpful comments and general advice during the conceptualization of this project.

## Authorship Contributions

Fig. 1. RS performed migration and in vitro cytotoxicity assays, FG and RS performed in vitro killing assays with and w/o Ca depletion and with and w/o prior cell sorting, FG, DS and MCA performed and interpreted RNAseq data from human samples, HW performed human S100A8/A9 ELISAs, FG performed S100A8/A9 PCR, PK analyzed S100A8/A9 mRNA expression in 5-AzaC treated THP-1 cells, MS performed S100A8/A9 blocking experiments. Fig. 2. RS, JHL, AS, DS and BK designed, performed and interpreted scRNAseq analysis of human samples. Fig. 3. BZ performed flow cytometric analysis of genome methylation and performed and interpreted MSP-PCR data. MK, MR and JS contributed to designing and interpreting the MSP-PCR. FCJ contributed to designing ATAC-seq analysis, SL performed ATAQ-seq, and SL, FCJ and DS analyzed ATAQseq, PK performed analysis of NF-κB–associated signaling molecules. Fig. 4. TV provided living S100A9^−/−^ mice and S100A9^−/−^samples of euthanized mice for RNAseq analysis, AN performed and interpreted RNAseq data of mouse samples together with DS. MS and BFB performed and interpreted in vivo experiments in S100A^−/−^ mice, TV performed murine S100A8/A9 ELISAs. Fig. 5. DS performed in silico analyses of patient data.

MOB and NC performed the sequencing, quality control and initial analysis of the RNAseq of human and mouse bone marrow samples, MBO performed data management of omics datasets and submission to public repositories. NP prepared the graphical design of our conceptualization (Fig. 1) and hypothesis (Fig. 6), AK and FG participated in performing initial experiments and in conceptualizing the early stages of this work, RS and DS participated in conceptualizing the work, participated in interpreting all data, performed all statistical analyses and prepared the first draft of manuscript. TV, MK, JS, DA, EMM, NG, NP, RH and UH interpreted data and substantially contributed to critical discussion. MCA conceptualized the work, acquired funding, designed experiments, interpreted data and wrote the manuscript. All authors critically revised the manuscript and approved of that latest version.

## Online Methods

### In vitro differentiation of CD34^+^ human stem cells (SC) to NK cells

CD34^+^ hematopoietic stem cells (SCs) from multiple donors were obtained from the Stem Cell Laboratory of the Departments of Pediatric Hematology and Oncology and Internal Medicine, University Hospital Tuebingen, Germany. Cells were differentiated towards the NK cell lineage for 31 days in RPMI-1640 (Sigma-Aldrich, R0883, Taufkirchen, Germany) supplemented with 10 % heat-inactivated fetal bovine serum (A5256701, Thermo Fisher Scientific, Darmstadt, Germany), 10 % heat-inactivated human serum (Sigma-Aldrich, H4522), 1 % penicillin/streptomycin (15070063, Thermo Fisher Scientific, Darmstadt, Germany), 1 % L-glutamine (A2916801, Thermo Fisher Scientific, Darmstadt, Germany), and a cytokine cocktail consisting of 20 ng/mL SCF (No. 11343327), 20 ng/mL Flt3-L (11343307), 20 ng/mL IL-7 (11340077), and 20 ng/mL IL-15 (11340157, all ImmunoTools, Friesoythe, Germany). An overview of the culturing procedure is shown in Fig. 1A. Half of the cultures were treated with 0.1 µM 5-Azacytidine (5-AzaC; Sigma-Aldrich, A2385, Taufkirchen, Germany) on days 7, 10, and 14. Cells were maintained at 1×10⁶ cells/mL, and medium was refreshed every 3–4 days. Importantly, in any process of medium change, adherent cells were gently detached using a cell scraper (Corning, 3010, Glendale, USA) in PBS (Sigma-Aldrich, D8537, Taufkirchen, Germany) and carefully combined with suspension cells. Both fractions were then pelleted at 350 × g for 10 min. and resuspended in fresh medium. In selected S100A8/A9 blocking experiments, Tasquinimod (SML2489, Sigma Aldrich, Taufkirchen, Germany) was added to the SC differentiation cell culture medium at a concentration of 10, 25 or 50uM daily on day 17-20 of culture. All antibodies used for flow cytometric detection of human cells are listed in **Suppl. Table 35**.

### Differentiation and polarization of the monocytic cell line THP1 to M1 and M2 macrophages

This experiment was performed according to a modified protocol^1^. Cells were cultured in 1640 RPMI medium supplemented with 10 % heat-inactivated FCS, 10 mM Hepes, 1 mM pyruvate and 2.5 g/l D-glucose. Differentiation into macrophages was induced by incubation with 150 nM PMA for 24 h, followed by 24h in normal culture medium. Polarization was achieved by adding 20 ng/ml IFN-γ (R&D Systems, Minneapolis, MN, USA) and 10 pg/ml LPS (0111:B4, Sigma Aldrich, Taufkirchen, Germany) m for M1 macrophages, or 20 ng/ml IL-13 and 20 ng/ml (R&D Systems, Minneapolis, MN, USA) IL-4 for M2 macrophages. An additional group of THP1 cells was cultured for 7 days in the medium described above supplemented with 0.1 µM 5-AzaC on days 1, 3, and 5.

### Isolation of CD14^+^ monocytes and CD14^−^/CD34^−^ immature lymphoid cells

CD14^+^ and CD14^−^/CD34^−^ cell fractions were isolated using magnetic bead–based separation (Miltenyi Biotec, Bergisch Gladbach, Germany). CD14^+^ cells were enriched with CD14 MicroBeads (130-050-201) using LS columns (130-042-401) according to the manufacturer’s instructions. The flow-through was subsequently depleted of CD34^+^ cells using CD34 MicroBeads (130-046-702) and LD columns (130-042-901) to obtain the CD14^−^/CD34^−^population.

### In vitro cytotoxicity assay of immature NK cells against K562 or Jurkat cells

On day 31 of SC differentiation, K562 and Jurkat cells were stained with 4 µM Vybrant CFDA SE Cell Tracker (V12883, Thermo Fisher Scientific, Darmstadt, Germany) for 15 min. at 37 °C. Differentiated SC-derived cells were then co-cultured with stained K562 (50:1) or Jurkat (10:1) target cells for 18 h at 37 °C in the NK cell differentiation medium described above. In selected experiments with Jurkat cells, 2 mM EGTA and 2 mM MgCl₂ were added to induce receptor-mediated, Ca²⁺-independent apoptosis^2^. Following co-culture, cells were stained with 0.02 % Pacific Blue (P10163, Thermo Fisher Scientific) and fixed in 0.5 % formaldehyde in PBS containing 2 % FBS and 2 mM EDTA (15575020, Thermo Fisher Scientific). In selected experiments, Tasquinimod (SML2489, Sigma Aldrich, Taufkirchen, Germay) was added at a concentration of 25 uM to the in vitro cytotoxicity assay (day 31) to block S100A8/A9. All samples were analyzed on a BD FACS Canto II (BD Biosciences, Heidelberg, Germany), and data were processed using FlowJo v10 (BD Biosciences, Heidelberg, Germany). To correct for spontaneous target cell death, monoculture controls were included in each experiment. Specific cytotoxicity was calculated as: (% CFSE⁺PB⁺ dead targets − % CFSE⁺PB⁺ spontaneous death) / (100 − % CFSE⁺PB⁺ spontaneous death) × 100 %.

### Cytotoxicity assays using sorted CD14^−^/CD34^−^ cells

In selected experiments, CD14⁺ monocytes were depleted on day 17 using magnetic bead isolation (see “Isolation of CD14⁺ monocytes and CD14⁻/CD34⁻ immature lymphoid cells”). The remaining CD14⁻/CD34⁻ immature lymphoid fraction was then cultured until day 31. Cytotoxicity against K562 cells was assessed using these CD14⁻/CD34⁻ fractions, differentiated either with or without CD14⁺ monocytes, as effector cells following the procedures detailed in “In vitro cytotoxicity assay of SC against K562 or Jurkat”.

### Human and murine S100A8, A9 and A12 ELISA assays

Supernatant of differentiating SC was collected on day 17 of differentiation by centrifugation for 10 min. at 300 g, followed by additional centrifugation for 10 min. at 2000 g. Supernatants were frozen at -80 °C until measurement. Human samples were analyzed for S100A8/A9 expression with Bühlmann MRP8/14 Calprotectin ELISA (Bühlmann Laboratories AG, Schönenbuch, Germany) according to manufacturer’s instructions, and for S100A12 using an in-house ELISA (normal level <150 ng/mL) as previously reported^3^. Murine serum concentrations of S100A8/A9 complexes were determined using an in-house ELISA as described previously^4^.

### Migration assay

On day 31, differentiating SC-derived cells were harvested, counted, and seeded onto 3 μm transwell inserts (833932300, Sarstedt, Nümbrecht, Germany) at 1×10⁶ cells/mL in the NK differentiation medium described in “In vitro differentiation of CD34⁺ SC to NK cells”. Cells were allowed to migrate for 4 h at 37 °C toward K562-derived tumor, generated by cultivating K562 cells for 24 h at 1×10⁶ cells/mL. Parallel cultures without inserts served as controls for normalization. After migration, cells were collected from the lower chamber, and CD14^+^ monocytes and CD56^+^ NK cells were stained for flow-cytometric identification. Subsequently, migrated cells were quantified using 123 counting beads (01-1234-42, Thermo Fisher Scientific, Darmstadt, Germany) on a BD FACS Canto II, and data were analyzed in FlowJo v10 (BD Biosciences, Heidelberg, Germany). Migration was first normalized to the no-insert control (100%) and subsequently normalized for +5-AzaC vs. untreated control. The resulting migration index was correlated with cytotoxicity determined in the K562 cytotoxicity assay (see “In vitro cytotoxicity assay of human immature day 31 NK cells against K562”).

### RT-PCR

RNA was isolated from differentiating +/- 5-AzaC-treated SCs (day 17 of culture) using the Qiagen RNeasy Mini Kit (74104, Qiagen, Hilden, Germany), followed by cDNA synthesis with the SuperScript™ IV VILO™ Mastermix with ezDNase™-Enzyme (11766050, Thermo Fisher Scientific, Darmstadt, Germany). Reverse transcription was carried out with the TaqMan™ Gene expression assays (FAM) (4331182, Thermo Fisher Scientific, Darmstadt, Germany; Assay IDs: Hs00374264_g1 (S100A8), Hs00610058_m1 (S100A9), Hs04187239_m1 (ID2), Hs00172872_m1 (EOMES), Hs00894392_m1 (TBX21), Hs00427620_m1 (TBP), Hs00995953_m1 (ZBTB20), Hs01049519_m1 (TOX), Hs00705412_s1 (NFIL3), Hs00169473_m1 (Perforin), Hs02786711_m1 (SPI1/PU.1) and Hs030239_g1 (ACTB). The log_2_FC was calculated using the standard 2^−ΔΔCt^ method^5^.

### Methylation-specific PCR analysis of S100A8 and S100A9

Genomic DNA from day 17 differentiating SCs was isolated using the QIAamp DNA Blood Mini Kit (51104, Qiagen, Hilden, Germany), with final elution in nuclease-free water. Bisulfite conversion of 500–800 ng genomic DNA was performed with the EpiTect Bisulfite Kit (59104, Qiagen, Hilden, Germany) according to the manufacturer’s instructions. For methylation analysis of S100A8 and S100A9, three genomic segments per gene (including promoter regions) were selected. Genomic sequences were obtained from Ensembl, and segment design was performed using DBTSS (https://dbtss.hgc.jp/) and MethPrimer 2.0 (http://www.urogene.org/methprimer2/)^6^. Segment 1 of the S100A9 promoter had been previously described^7^. Bisulfite-converted DNA (approx. 30 ng) was then amplified using GoTaq Hot Start Green Master Mix (M5122, Promega, Fitchburg, USA). PCR products were cloned using the NEB PCR Cloning Kit (E1202S, New England Biolabs, Frankfurt, Germany), transformed, and selected on LB–ampicillin plates (100 µg/mL). For each segment, ≥15 colonies were screened by colony PCR to obtain 8–10 positive sequences for methylation analysis. Colony PCR was performed using GoTaq Colorless Master Mix (M7132, Promega, Fitchburg, USA). Bacterial DNA templates were prepared by resuspending individual colonies in 20 µL nuclease-free water, of which 4 µL were used per PCR. PCR products were analyzed on 1.2% agarose gels, and products of expected size were sequenced by Eurofins Genomics (Ebersberg, Germany). Methylation status was evaluated BiQ Analyzer^8^.

### Bulk RNA Sequencing experiments

#### Library preparation (human CD34+ SCs)

Libraries were generated from total RNA after poly(A)^+^ mRNA enrichment using the NEBNext Ultra II Directional RNA Library Prep Kit with NEBNext Multiplex Oligos (NEB) according to the manufacturer’s instructions. Sequencing was performed on an Illumina NovaSeq 6000 using 150 bp paired-end reads.

#### Library preparation (mouse BMC)

Libraries were generated from total RNA after poly(A) mRNA enrichment using the NEBNext Ultra II Directional RNA mRNA UMI Kit (NEB) with 100 ng RNA input, according to the manufacturer’s instructions. Sequencing was performed using 100-base paired-end reads on the Illumina NovaSeq 6000.

#### Preprocessing (human CD34+ SCs)

Raw FASTQ files were quality-controlled using ngs-bits (v2019_11) to assess per-cycle base quality and adapter contamination. Reads were aligned to the GRCh37 reference genome using STAR (v2.7.3a). Alignment quality was evaluated with ngs-bits and by visual inspection in IGV (v2.4.14). Gene-level counts were generated with featureCounts (Subread v2.0.0) using Ensembl release 95 annotations; no additional pre-filtering was applied (analysis performed in R v4.3.0).

#### Preprocessing (mouse BMC)

Quality control, alignment, and quantification were performed within the Nextflow-based nf-core/rnaseq pipeline (v1.4.2). Raw FASTQ files were quality-controlled using FastQC (v0.11.8) and RSeQC (v3.0.1). Reads were aligned to the GRCm38 reference genome using STAR (v2.6.1d). Gene-level counts were generated with featureCounts (v1.6.4).

#### Preprocessing (GSE76203)

Raw data from Unnikrishnan et al.^9^ were accessed via the Gene Expression Omnibus (GEO) under accession GSE76203. Read quality was assessed using FastQC (v0.10.1). TruSeq sequencing adaptors were clipped using cutadapt (v4.0) with Illumina adaptor sequences (Illumina, San Diego, USA). A Phred score cutoff of 20 was applied for per-base quality and reads shorter than 30 bases were discarded. Reads were aligned using HISAT2 (v2.1.0) to GRCh38. Read counting was performed using featureCounts (v2.1.1), generating per-gene read counts for each sample. Count files were merged into a single tab-delimited matrix; no additional pre-filtering was applied (analysis performed in R v4.3.0).

#### Differential expression analysis

Differential expression analysis was performed using DESeq2 (v1.40.2)^10^. Gene counts were modeled with a negative binomial generalized linear model. For paired samples, pairing was accounted for by including subject as a blocking factor together with treatment status (post vs. pre) in the design formula (design = ∼ subject + treatment). For unpaired comparisons, the design formula included treatment only (design = ∼ treatment). Differential expression was tested using the Wald test implemented in DESeq2, and p values were adjusted for multiple testing using the Benjamini–Hochberg method.

#### Gene set enrichment analysis (GSEA)

Pre-ranked and classic GSEA were performed using the Broad Institute GSEA Desktop Application (v4.4.0)^11^. For paired experiments, genes were ranked by DESeq2’s Wald statistic (“stat”) from the paired differential expression model described above, and pre-ranked GSEA was applied. For unpaired experiments, DESeq2 size-factor–normalized counts were used as input for classic GSEA. In all analyses, 10,000 permutations were performed. Gene sets from the MSigDB Hallmark (H) and C5:GO:BP (Gene Ontology Biological Process) collections, as well as the IVANOVA^12^ collection, were tested; for the IVANOVA^12^ collection, all eight gene sets were combined into a single file and analyzed together. For GSE76203, ssGSEA was additionally performed on DESeq2 size-factor–normalized counts using GSVA (v1.48.3) with MSigDB Hallmark (H) gene sets.

#### GO term analysis

GO term enrichment analysis was performed using clusterProfiler (v4.8.3)^13^. All DESeq2 significant genes (cutoff p.adj. < 0.05) were imported into R as Ensembl gene IDs and mapped to gene symbols using org.Hs.eg.db (v3.17.0) and AnnotationDbi (v1.62.2).

#### Cell type deconvolution

Cell-type fractions were estimated from bulk RNA-seq data using CIBERSORTx^14^ with two reference strategies. For healthy d17 stem cell samples, a signature matrix was generated from the CPM-normalized scRNA-seq reference “Fig2e-5PBMCs_scRNAseq_matrix.txt” provided on the CIBERSORTx website (default settings), and fractions were inferred using S-mode batch correction, 1000 permutations, and absolute mode. For patient CD34⁺ HSPC samples, a signature matrix was generated from CPM-normalized sorted HSPC bulk RNA-seq profiles (GEO GSE74246) using default parameters, and fractions were inferred using B-mode batch correction, 1000 permutations, and absolute mode.

Plots were generated in R using ggplot2 (v3.5.0) unless stated otherwise. Radar plots were produced with fmsb (v0.7.6), and Venn diagrams were generated using VennDiagram (v1.7.3)^15^.

### Single cell RNA-seq and CITE-seq

scRNA-seq was performed on cryopreserved differentiating SCs of day 17 of culture. Samples were tagged with Total-Seq HTO antibodies (CITE-seq, **Suppl. Table 36**) and incubated simultaneously with a selection of Total-Seq antigen-specific antibodies according to the manufacturers’ instructions (Biolegend, San Diego, USA) (.protocols.io/view/totalseq-a-antibodies-and-cell-hashing-with-10x-si-261geo6mol47/v1). Following the Cite-Seq staining protocol, 24 subsamples were consolidated into two samples, to be further processed across two 10x Genomics lanes. For each lane, differentiated SC were isolated using an Aria Fusion Sorter (BD Biosciences, Heidelberg, Germany). The cell concentration was standardized to 20.000 cells per 10x lane. Immediately post-sorting, chilled cells were loaded onto the 10x Chromium system for processing, facilitating the generation of droplet-based single-cell RNA sequencing datasets (Chromium platform, 10x Genomics, Pleasanton, USA). The standard protocol for the 10x single-cell kit (V3.1 chemistry) was applied comprehensively for all steps, including cell loading, library preparation, and quality control. For the construction of the Total-Seq libraries (HTO and ADT), the specified, afore-mentioned protocol was employed. The cDNA concentrations were meticulously adjusted to align with the operational specifications of the S2 flow cell and sequenced utilizing an Illumina NextSeq system. The multiplexed sequencing raw data consisting of mRNA library reads as well as labelled ADT and hash tag oligos (HTO) were processed using the qbic-pipelines/cellranger pipeline (version 2.0; commit dd28b16, github.com/qbic-pipelines/cellranger) with default parameters. This pipeline functions as a wrapper for the 10x Genomics cellranger-multi pipeline based on the nf-core framework. As part of this pipeline, the quality of the raw data (FASTQ files) was first assessed usingFastQC (version v0.11.9) and aggregated for visualization using MultiQC (version1.14; http://multiqc.info/)^16^. Subsequently, the raw reads were filtered and aligned to the reference human genome (GRCh38) to generate feature barcode matrices for each sample.

#### Data processing

Gene expression, CITEseq (antibody-derived tag, ADT), and hashtag oligonucleotide (HTO) count matrices were imported into Seurat (v5.2.1) running in R (v4.3.3) for downstream single-cell analysis. Initial quality control was performed on gene expression data, retaining cells with 200–4.500 detected features and less than 20% mitochondrial transcript content. Low-quality cells and doublets were removed at this stage. HTO counts were normalized using centered log-ratio (CLR) transformation and demultiplexed using the MULTIseqDemux function to assign cells to individual samples. Following demultiplexing and quality control, gene expression, ADT, and HTO information were combined into a finalized Seurat object that was used for all downstream analyses. No additional demultiplexing or singlet filtering was applied thereafter. For transcriptomic analyses, samples were initially processed separately by condition (“responder” vs. “non-responder”; 5-AzaC vs. control) to assess potential batch- or condition-specific effects on clustering. Gene expression data were normalized using NormalizeData, highly variable features were identified using FindVariableFeatures, and data were scaled using ScaleData with default parameters. Linear dimensionality reduction was performed by principal component analysis (PCA), followed by construction of a k-nearest neighbour graph (FindNeighbors, dimensions 1–35). Unsupervised clustering was carried out using the Leiden algorithm (FindClusters, resolution = 0.7), and non-linear dimensionality reduction was performed using UMAP (RunUMAP, dimensions 1–35). To facilitate downstream differential expression and integrative analyses, all samples were subsequently merged and reprocessed from normalization onward. During the scaling step, mitochondrial and cell-cycle–associated genes were regressed out to mitigate their influence on clustering and differential expression. Cell type annotation was performed using SingleR, supported by manual inspection of canonical marker expression and interactive visualization with ShinyCell. Clusters were annotated as granulocyte–monocyte progenitors (GMPs), monocyte subsets, polymorphonuclear neutrophils (PMNs), dendritic cells, mast cells, and NK cells. For differential expression analyses, pseudobulk profiles were generated using AggregateExpression to account for within-sample correlations. Differential expression testing was performed on pseudobulk profiles using FindMarkers with DESeq2. In addition, targeted within-monocyte comparisons were performed by subsetting hyinflMo, IntMo, and inflMo cells and running pairwise differential expression with FindMarkers using the likelihood-ratio test (test.use = “LR”) while adjusting for donor, response group, and treatment (latent.vars = c(“donor_id”,“group”,“vidaza”)); results were visualized with volcano plots and exported as full and significant gene tables.

#### ADT analysis

ADT data were analyzed using the finalized Seurat object after transcriptomic quality control. The ADT assay was set as an active assay and normalized using centered log-ratio (CLR) normalization across cells. NK cells were identified based on transcriptomic clustering and sub-setted for downstream protein-level analyses. To enable donor-aware comparisons, ADT expression was aggregated at the donor level by calculating the median CLR-normalized expression per marker across all NK cells from each donor and condition. Only donor–marker combinations supported by a minimum of three NK cells were retained. Comparisons between “responders” and “non-responders” under 5-AzaC treatment were performed using donor-level ADT summaries. Given the limited number of donors (n = 3 per group), statistical testing was considered exploratory, and effect sizes and consistency across donors were prioritized for interpretation. ADT results were used as protein-level support for transcriptomic and pathway-based findings.

#### Gene set enrichment analysis (GSEA)

GSEA was performed using the R package *fgsea* (v1.36.0). GSEA was applied to differential expression results obtained from Seurat. For each contrast, genes were ranked using a composite ranking metric defined as *rank*g = *avg*log2FC,g × [−log10(*p*g)], where *avg*log2FC,g denotes the average log2 fold change and *p*g the corresponding p-value. Gene set enrichment was assessed using the Hallmark gene set collection (MSigDB Hallmark). Only gene sets containing between 10 and 500 genes were included. Enrichment statistics, including normalized enrichment scores (NES), nominal p-values, and Benjamini–Hochberg adjusted p-values (FDR), were estimated using the multilevel algorithm (*fgseaMultilevel*).

#### Pseudotime analysis

For trajectory inference, the Seurat object was converted into a Monocle3-compatible *cell_data_set* (CDS) using Monocle3 (v1.4.26). The CDS was preprocessed by principal component analysis (30 PCs) using *preprocess_cds()* and embedded using UMAP via *reduce_dimension()* with PCA preprocessing. Cells were clustered using *cluster_cells()*, and partitions were computed; only the largest partition was retained for downstream pseudotime analysis. Cell types represented by fewer than five cells within this partition were excluded to improve robustness of trajectory learning. The filtered CDS was reclustered, and a principal graph was learned using *learn_graph()* with *use_partition = FALSE*. To further refine the granulocyte–monocyte progenitor (GMP) compartment, GMP cells were extracted and sub-divided into two sub-clusters using k-means clustering on Monocle3 UMAP coordinates (k = 2; *set.seed*(1); *nstart* = 10). The subcluster occupying the earliest region of the GMP manifold was designated GMP_root, while the adjacent, dendritic cell–biased cluster was labeled GMP_DCbranch and stored as a GMP subtype annotation. Cells were ordered in pseudotime using *order_cells()* on the UMAP embedding, specifying GMP_root cells as the root state. Pseudotime values were extracted using *pseudotime()* and GMP subtype annotations were transferred back to the Seurat object for downstream visualization and integration with transcriptomic analyses. Pseudotime trajectories were visualized on the UMAP embedding with the learned principal graph overlaid. Pseudotime distributions were summarized using violin plots with overlaid boxplots across annotated cell types and, where indicated, stratified by treatment and response group. All figures were generated in R using *ggplot2* (v4.0.1).

#### S100A8/A9 single-cell expression

Per-cell log-normalized expression values (Seurat RNA, data slot) were visualized across clusters using violin plots with overlaid jittered single -cell points (ggplot2).

### Assay for Transposase-Accessible Chromatin with sequencing (ATAC-seq)

#### ATAC library generation and sequencing

ATAC-seq was performed to assess day 17 5-AzaC-treated and untreated SSCs genome-wide chromatin accessibility. ATAC-seq libraries were generated using the Omni-ATAC protocol optimized for cryopreserved lymphocytes^17^. Briefly, nuclei were isolated from 5-AzaC–treated and untreated CD14⁺ and CD14⁻CD34⁻ SSCs from three donors, followed by Tn5 transposase-mediated tagmentation and library preparation. Libraries were pooled and sequenced on an Illumina HiSeq 3000, yielding a mean depth of 46 million reads per sample.

#### Pre-processing

From the raw read data, the adaptors were trimmed (using cutadapt to remove Nextera adapter sequences from both 3′ and 5′ ends, requiring a minimum 5-bp overlap; -O 5) and reads shorter than 30 bp were removed (cutadapt; -m 30). Then the trimmed reads were aligned to the reference genome (GRCh38) using Burrows-Wheeler Alignment tool (BWA)^18^ (Li and Durbin et al., 2009). Mitochondrial genome and possible PCR duplicates, reads which piled up with same start and end coordinates, were subsequently removed from alignment output using SAMtools^19^ (Li and Handsaker et al., 2009) to generate mitochondria-excluded, de-duplicated BAM files. Library quality was assessed using ATACseqQC in R to determine the enrichment of reads around transcription start sites of genes (a good quality library shows a high enrichment score). TSSE scores varied across samples and the one with lowest quality sample was excluded from subsequent pooled peak calling in MACS2. To ensure peaks were called using a balanced design comprising the same number of treated and untreated cells of each cell type, further four libraries were excluded from pooled peak calls. To equalize sequencing depth across libraries prior to pooled peak calling, each remaining library was downsampled to 5 million mapped reads/fragments and then merged for pooled peak calling by MACS2^20^ in paired-end mode (BAMPE) with corresponding pooled data from genomic sequencing libraries as input controls. Coordinates of the called peaks were extracted from the MACS2 narrowPeak output and converted to GFF format, and considering each individual’s entire readset separately, reads falling in the pooled peaks were counted using htseq (htseq-count; intersection-strict, non-unique reads excluded). Peak visualization: BigWig signal tracks were used for region-specific visualization of chromatin accessibility, and plots were generated in R using ggplot2 (v4.0.1). Differentially expressed peaks: This was performed in R using DESeq2 (v1.50.2), with treatment condition and pairing included as design factors. The combined peak count matrix was first separated into CD14^+^ and CD34^−^CD14^−^ cell fractions. Peaks with raw counts <5 in at least 3 samples were filtered out, followed by normalization using DESeq2’s median-of-ratios method. Differential accessibility was assessed with an FDR cutoff of q ≤ 0.05. Volcano plots showing all peaks for CD14^+^ and CD34^−^CD14^−^ cells were generated using ggplot2 (v4.0.1). Gene ontology (GO) term analysis: Functional enrichment analysis of differentially accessible regions was performed using the rGREAT web interface (GREAT) with Gene Ontology terms. Enrichment significance was assessed using the region-based binomial test. For reporting, terms were filtered to retain those with ≥5 observed region hits.

#### Transcription factor (TF) motif enrichment and foot printing

Transcription factor motif enrichment and foot printing in accessible chromatin were performed with TOBIAS v0.17.1. Aligned, de-duplicated and mitochondrial DNA–removed BAM files were down-sampled to 30.000.000 reads per sample using samtools v1.16.1 and then merged by cell type and treatment into four pooled groups (three donors per group; ∼90.000.000 reads per merged BAM): CD14^+^ + 5-AzaC, CD14^+^ - 5-AzaC, CD34^−^CD14^−^ + 5-AzaC, CD34^−^CD14^−^ - 5-AzaC. Peaks were called on each merged BAM using MACS2 v2.2.7.1 in paired-end mode (-f BAMPE) with –nomodel, –shift -100, –extsize 200, –keep-dup 1, and –call-summits, applying a stringent q-value cutoff (q ≤ 0.01). For each lineage, a consensus peak set was generated by concatenating the corresponding group peak files, sorting, and merging overlapping intervals using bedtools v2.31.1 (bedtools sort/merge). Tn5 insertion bias was corrected with TOBIAS ATACorrect (hg38 reference; blacklist filtering enabled), and footprint scores were computed with TOBIAS FootprintScores using the bias-corrected signal tracks and the lineage-specific consensus peak regions. Differential TF binding was assessed with TOBIAS BINDetect using JASPAR CORE non-redundant motifs as reference. Results were visualized in R using ggplot2 (v4.0.1).

### Proteome Profiler Human NFkB Pathway Array

Characterization of the NFκB pathway was performed using the Human NFκB Pathway Array (ARY029, R&D Systems, Minneapolis, USA) according to the manufacturer’s instructions. CD14⁺ and CD14⁻/CD34⁻ fractions were sorted on day 21 (see “Isolation of CD14⁺ monocytes and CD14⁻/CD34⁻ immature lymphoid cells”), cultured overnight and lysed the next day for protein extraction. Protein concentrations were determined using the BCA Protein Assay Kit (Thermo Fisher Scientific, Darmstadt, Germany) and measured on a NanoDrop 2000 (Thermo Fisher Scientific, Darmstadt, Germany). Array membranes were blocked as instructed, and 250 µg protein per sample was applied and incubated overnight at 4 °C on a plate rotator. The next day, membranes were incubated with the supplied antibody cocktail for 1 h, followed by IRDye 800CW Streptavidin (LI-COR Biosciences, Lincoln, USA) for 30 min. in the dark, replacing the standard HRP-Streptavidin step. Signal detection was performed using the LI-COR Odyssey FC imager, and data were analyzed with ImageJ (NIH, Bethesda, MD, USA).

For each protein, fold-change (FC) relative to the untreated control was calculated. Biological triplicates were analyzed for CD14^+^ and CD14^−^/CD34^−^. FC values were log2-transformed prior to statistical analysis. One-sample t-tests were performed for each protein and cell type to assess whether the mean log2 fold-change was significantly different from 0 (corresponding to no change relative to control). P-values were adjusted for multiple comparisons using the false discovery rate (FDR) method. A heatmap was generated using the pheatmap package in R, displaying the mean log2 fold-change for each protein and cell type. Proteins were hierarchically clustered based on similarity of expression profiles across cell types using Euclidean distance and complete linkage.

### Animal experiments

Male and female mice aged 6–14 weeks were used for all in vivo studies. On day 0 of the experiment, animals received a sublethal single dose of total body γ-irradiation (100 cGy) to eradicate the hematopoiesis and to induce regeneration of newly emerging hematopoietic cells. On days 7, 10, and 14 after γ-irradiation, mice were treated intraperitoneally with 5-AzaC at a dose of 5 mg/kg, control animals received PBS alone. In some experiments, mice were subcutaneously injected on day 21 with 1 × 10⁶ RMA-S cells; control animals received PBS only. Animals were monitored regularly, and experiments were terminated at humane endpoints determined by tumor size, between days 28 and 33 post-irradiation. Post euthanasia, spleen and bone marrow cells were subjected to phenotypic analysis of engraftment using flow cytometry or were used as immature NK cell effectors for ex vivo cytotoxicity assays against RMA-S cells as targets (Ratio 100:1). To test mature NK cell cytotoxicity, naïve C57BL/6J and S100A9^−/−^ mice were euthanized likewise used as effectors against RMA-S cells as targets (Ratio 100:1). All antibodies used for flow cytometric detection of murine cells are listed in **Suppl. Table 35**.

### Kaplan–Meier analysis

Publicly available myelodysplastic syndrome (MDS) and chronic myelomonocytic leukemia (CMML) cohort published by Unnikrishnan et al.^9^ and additionally provided unpublished data by the same authors comprising of 28 matched samples from 14 patients across three sequencing batches was used to assess the impact of 5-AzaC response on overall survival of MDS/CJMML patients. Overall survival was defined as time from treatment initiation to death; patients alive at their last follow-up were censored. Survival curves were estimated using the survfit function and plotted using ggsurvplot. Statistical comparisons were performed using the log-rank (Mantel–Cox) test. A Cox proportional hazards model was fitted using coxph to estimate hazard ratios (HR) with 95% confidence intervals (CI). Model validation used likelihood ratio and Wald tests.

### Ethical statement on the use of CD34+ SCs and the performance of animal experiments

This study on human stem cells was approved by the Local Institutional Ethic’s Review Board (23/2007, 199/2010BO1, 213/2014BO2, 285/2021BO2). Informed consent was obtained from all enrolled stem cell donors. Animal experiments were approved by the Local Review Board (Regierungspräsidium Tuebingen, Germany, K2/16, K03/19G, K06/20G, K03/21G, K 03/22 G and § 4 notification 2021.02.01) and were performed according to Institutional Guidelines.

### Statistics

Statistical analyses were performed using R (version 4.x) and GraphPad Prism. Unless not stated otherwise, statistical testing was conducted at the level of independent biological replicates (human donors or mice, respectively). Sample sizes (n) are reported in the corresponding figure legends and denote the number of independent donors or animals. No statistical methods were used to predetermine sample size. For comparisons between two independent groups (e.g., “responders” vs. “non-responders”), two-sided Welch’s t-tests were used when appropriate; otherwise, two-sided Wilcoxon rank-sum (Mann–Whitney U) tests were applied. For paired comparisons within the same donor (e.g., 5-AzaC treatment vs. control), two-sided paired t-tests or Wilcoxon signed-rank tests were used, as indicated. One-sided tests were applied only in predefined directional comparisons and are explicitly stated in the corresponding figure legends. For experiments with repeated measurements across treatment conditions within donors, two-way repeated-measures ANOVA or mixed-effects models were used (GraphPad Prism), with treatment condition and response group included as fixed effects and donor status included as a repeated/random factor. Mixed-effects analyses were fitted using Prism’s restricted maximum likelihood (REML) approach with donor as a random intercept. Post-hoc multiple-comparison testing was performed using Sidak’s or Tukey’s correction, as indicated. Differential gene expression analyses of bulk RNA-seq data were performed using DESeq2 with Wald testing and Benjamini–Hochberg false discovery rate (FDR) correction. For scRNA-seq analyses, donor-level pseudobulk profiles were used to avoid pseudo-replication from single-cell measurements, except for targeted differential expression between monocyte states as described above. Gene Ontology enrichment was assessed by one-sided hypergeometric (Fisher’s exact) over-representation testing with Benjamini–Hochberg correction. GREAT-based enrichment of differentially accessible ATAC-seq regions was evaluated using GREAT’s region-based binomial test, reported as Binomial FDR q values. GSEA was performed using permutation-based testing (10.000 permutations), reporting enrichment scores (ES), normalized enrichment scores (NES), nominal p values, and FDR q values. For donor-level cluster proportion comparisons across multiple clusters, p values were computed per cluster; multiple-testing correction (Benjamini–Hochberg) was applied where specified. Where unadjusted p values are shown, these represent exploratory statistics for predefined comparisons and adjusted values are provided in the **Supplementary Statistics Table**. Correlation analyses were performed using Pearson’s correlation coefficient (r) with two-sided testing. Effect sizes are reported in the Fig. Legends where applicable (e.g., Cliff’s delta for distributional differences). Normality was assessed visually, and non-parametric tests were used when sample sizes were small, or distributions deviated from normality. Data are presented as mean ± standard deviation unless specified otherwise. Box plots indicate the median and interquartile range, with whiskers representing 1.5× IQR. Exact p values are reported where appropriate; significance thresholds are denoted as *p ≤ 0.05, **p ≤ 0.01, ***p ≤ 0.001, ****p ≤ 0.0001. All tests were two-sided unless explicitly stated otherwise.

### Data availability statement

The data discussed in this publication have been deposited in NCBI’s Gene Expression Omnibus^21^ (Edgar et al.,) and are accessible through GEO Series accession number GSEXXXX.

## Supplemental Figure Legends

**Suppl. Fig. S1. Proliferation rate in 5-AzaC treated CD34^+^ SC cell cultures.** Cell numbers were quantified at the indicated time points during SC differentiation. Proliferation is shown as fold change in 5-AzaC–treated cultures relative to untreated controls (dashed line). Statistical significance was assessed by unpaired Student’s t-test (*p ≤ 0.05). n=7 “responders”, n=27 “non-responders”.

**Suppl. Fig. S2. Flow cytometric assessment of cell composition during SC differentiation.** Relative frequencies of major cell populations were quantified by flow cytometry on day 17 **(A)** and day 31 **(B)** in cultures treated ±5-AzaC. n = 14 per condition; paired Student’s t-test.

**Suppl. Fig. S3. Expression of NK cell lineage-specific transcription factors during early SC differentiation.** qRT-PCR analysis of selected transcription factors on day 10 and d14 of CD34^+^ differentiation. Expression changes are shown as log fold change (5-AzaC/control) calculated using the 2^−ΔΔCt^ method. n = 4 donors; two-way ANOVA with Dunnett’s multiple-comparisons test.

**Suppl. Fig. S4. Induction of S100A8 and A9 transcripts in 5-AzaC treated “responder” cultures.** qRT-PCR analysis of S100A8 and A9 expression on day 17 of SC differentiation. Data are shown as log fold change (5-AzaC/control) using the 2^−ΔΔCt^ method. n = 5 donors; one-way ANOVA with Dunnett’s multiple-comparisons test.

**Suppl. Fig. S5. Cell-type–specific methylation levels of S100A8 and S100A9 in human hematopoietic populations.** In silico analysis of DNA methylation levels across CD34^+^ stem cells and mature blood cell types using the EWAS (Epigenome-Wide Association Study) Atlas database^22^. Methylation values are displayed as box-and-whisker plots (subject numbers as indicated).

**Suppl. Fig. S6. Tasquinimod effect on viability and endogenous S100A8/A9 levels. (A)** Percentage (%) of cell death of the K562 cell line upon incubation with 25 uM of Tasquinimod for 18 hours as determined by flow cytometry (n = 5, unpaired t-test). **(B, C)** Endogenous S100A8/A9 concentrations in untreated **(B)** and 5-AzaC-treated **(C)** SC cultures in the presence of increasing Tasquinimod concentrations (10–50 μM), quantified by ELISA and shown as fold change relative to vehicle control (n = 5; two-way ANOVA with Tukey’s multiple-comparisons test).

**Suppl. Fig. S7. scRNA-seq: Identification of cell-type identities.** Cell-type identities were assigned based on canonical marker gene programs, i.e. NK cells were defined by coordinated expression of *NCAM1*, *PRF1* and *GZMB*. Monocyte subsets expressed *CD14*, *CSF1R* and *FCGR1A*, PMN were characterized by high levels of granule-associated transcripts such as *MPO* and *ELANE,* cDC2 expressed *CLEC10A*, *CD1C* and *FCER1A*; mregDC were distinguished by additional upregulation of *LAMP3*, *EBI3* and *CCR7*, pDC selectively expressed *LILRA4*, *CLEC4C* and *IL3RA*. Among immature myeloid populations, eMoP and preMo co-expressed *FCGR1A*, *CEBPA* and low levels of *MPO*. The NMP cluster exhibited a mixed monocytic–neutrophil pattern with concomitant expression of *CD14*, *CSF1R*, *MPO* and *ELANE*. The most immature progenitors - GMP and MEP - showed expected stem cell (SC)-like signatures, including *CD34* and *HBB*, with GMP additionally expressing *CD38*, *ELANE* and *MPO,* and MEP expressing erythro-megakaryocytic markers such as *KLF1*, *ITGA2B* and *PF4.* Bubble plot of canonical marker gene expression across clusters. Dot size represents the fraction of expressing cells and the dot color shows the average normalized expression.

**Suppl. Fig. S8. scRNA-seq: Donor-level distribution of inflammatory monocyte subsets.** Proportions of inflMo, IntMo, and hyinflMo subsets are shown for individual “responder” and “non-responder” (n=6) donors. Control and 5-AzaC–treated samples are indicated by point shape. Two-sided Wilcoxon rank-sum test; Cliff’s delta is reported as effect size.

**Suppl. Fig. S9. scRNA-seq: Monocle3 pseudotime distributions across refined myeloid subsets.** Violin plots showing pseudotime distributions across annotated myeloid cell states, with embedded boxplots indicating median and interquartile range.

**Suppl. Fig. S10. scRNA-seq: Hallmark pathway enrichment across inflMo, IntMo and hyinflMo clusters.** Heatmap showing the normalized enrichment scores (NES) across curated Hallmark pathways in n=6 donors. Pathways are grouped by functional categories (Wilcoxon signed-rank test with Benjamini–Hochberg correction; *FDR < 0.05. The corresponding terms are written in bold letters.

**Suppl. Fig. S11. scRNA-seq: Treatment-associated Hallmark pathway changes in „responder” inflMo and “non-responder” IntMo and hyinflMo clusters (treated vs. untreated).** Heatmap shows the normalized enrichment scores (NES) across curated Hallmark pathways in n=6 donors (n=3 per group). Pathways are grouped by functional categories. Colors indicate NES, and asterisks denote (Wilcoxon signed-rank test with Benjamini–Hochberg correction; *FDR < 0.05). Corresponding terms are written in bold letters.

**Suppl. Fig. S12. scRNA-seq: DGEA of inflammatory monocytes.** Volcano plots of differential gene expression between 5-AzaC–treated and control samples within inflammatory monocyte subsets. Red dots indicate significantly up-regulated genes (p < 0.05 and |log2FC| ≥ 0.5), blue dots indicate significantly down-regulated genes, grey dots denote non-significant genes. Identification of the top 15 most significant genes (ranked by p-value) per subset.

**Suppl. Fig. S13. scRNA-seq: Single-cell expression of S100A8 and S100A9 across various clusters.** Violin plots with overlaid jittered points show log-normalized single-cell expression of S100A8 and S100A9 across annotated various hematopoietic progenitor clusters, stratified by donor (n=6) response group (“responders” vs. “non-responders”) and treatment condition (5-AzaC vs. untreated control). Each dot represents one cell, and violins indicate the distribution of expression values within each cluster–condition group. Expression is shown at the single-cell level without pseudobulk aggregation. Cell numbers per cluster and condition are provided in the accompanying panel. The fraction of S100A8- or S100A9-expressing cells (>0) was calculated directly from single-cell data. Highlighted are monocytic clusters (dark lila) and the NK cell cluster (light lila).

**Suppl. Fig. S14. scRNA-seq: Treatment-associated Hallmark pathway changes in „responder” and “non-responder” NK cells (treated vs. untreated).** Heatmap shows the normalized enrichment scores (NES) across curated Hallmark pathways in n=6 donors (n=3 per group). Pathways are grouped by functional categories. Colors indicate NES, and asterisks denote (Wilcoxon signed-rank test with Benjamini–Hochberg correction; *FDR < 0.05). Corresponding terms are written in bold letters.

**Suppl. Fig. S15. scRNA-seq: DGEA of NK cells.** Volcano plots of differential gene expression between 5-AzaC–treated and control samples within NK cells (|log2FC| ≥ 0.5, adjusted p < 0.05). Red dots indicate significantly up-regulated genes (p < 0.05 and |log2FC| ≥ 0.5), blue dots indicate significantly down-regulated genes, grey dots denote non-significant genes. Identification of the top 15 most significant genes (ranked by p-value) per subset.

**Suppl. Fig. S16. scRNA-seq: Differences in NK cell-directed intercellular communication between “responders” and “non-responders”. (A)** Summed ligand–receptor interaction scores from all sender clusters to NK cells under 5-AzaC treatment relative to control. **(B)** Changes in weighted interaction load (Δ = treatment − control) across sender clusters in “responders” and “non-responders”. Only active LR pairs and clusters meeting minimum cell number of 10 were included. n=6 donors, n=3 per group.

**Suppl. Fig. S17. Methylation status of S100A8 and S100A9 in “responders” and “non-responder”. (A**) Flow cytometric analysis of global 5-methylcytosine (5-mC) levels during SC differentiation in the presence or absence of low-dose 5-AzaC. Shown is the fold change (FC) in MFI of 5-mC expression. **(B - G)** Methylation-sensitive PCR (MSP-PCR) analysis of CpG methylation across the S100A8 (B–D) and S100A9 (E–G) loci in day 17 CD34⁺ SC-derived cultures from “responders” and “non-responders”. Schematics indicate analyzed CpG regions relative to the transcription start site (TSS). Methylation percentages were calculated from sequencing of 9–10 bacterial clones per region (n = 5 donors; two-way ANOVA with Šídák’s multiple-comparisons test).

**Suppl. Fig. S18. ATAC-seq: Chromatin accessibility profiles at the S100A8/A9 loci.** Genome browser tracks showing chromatin accessibility profiles from the ATAC-seq at the indicated loci in sorted day 17 CD14^+^ fractions from +5-AzaC and untreated control samples. (n = 4 “responder” donors).

**Suppl. Fig. S19. Heatmap of differentially expressed transcription factor genes in C57BL/6J and S100A9^−/−^ mice.** Values represent row-wise Z-scores of DESeq2-normalized expressions with hierarchical clustering and dendrograms. Differential expression was assessed using DESeq2 (Wald test with Benjamini–Hochberg FDR correction). n= 4 mice per group.

**Suppl. Fig. S20. Experimental design and gating strategy for murine in vivo studies. (A)** Schematic overview of irradiation, 5-AzaC treatment and tumor priming. **(B)** Flow cytometric gating strategy for identification of murine hematopoietic precursor populations.

**Suppl.** Fig. 21**. Hematopoietic reconstitution in irradiated mice. (A, B)** Bone marrow reconstitution in irradiated WT ±5-AzaC and untreated S100A9^−/−^ mice under (A) PBS or (B) RMA-S priming conditions. (n = 4-8; two-way ANOVA with Šídák’s multiple-comparisons test).

**Suppl. Fig. S22. Serum S100A8/A9 levels in 5-AzaC–treated mice.** S100A8/A9 concentrations were measured by ELISA in serum from naïve or tumor-bearing WT mice treated ±5-AzaC (days 28–33 post irradiation). n = 6–9; two-way ANOVA with Šídák’s multiple-comparisons test.

**Suppl. Fig. S23. Contribution of CD8⁺ T cells to immature NK cell cytotoxicity in vivo.** Quantification of immature NK cell cytotoxicity in the presence or absence of CD8⁺ T cells in WT ±5-AzaC and S100A9^−/−^ mice (n =1-6) (details as indicated). T cell depletion of > 90% was confirmed with flow cytometry.

**Suppl. Fig. S24. Transcriptional inflammatory signatures in 5-AzaC treated MDS/CMML CD34⁺ cells from the Unnikrishnan cohort. (A, B)** Bubble plot of the top 25 enriched (A) cellular component (CC) and (B) molecular function (MF) GO terms among genes upregulated in treated vs. untreated “responders”. Over-representation was tested using a one-sided Fisher’s exact (hypergeometric) test with Benjamini–Hochberg correction; dot size indicates gene count, color indicates −log10(p-adj.), and the x-axis shows gene ratio.

**Suppl. Fig. S25. ssGSEA Hallmark pathway activity for MYC targets and E2F targets.** Boxplots show ssGSEA scores stratified by response (“responder” vs. “non-responder”) and treatment (untreated vs. treated), with individual patient values overlaid. Statistical differences were evaluated using a hybrid Wilcoxon testing framework across all six pairwise comparisons among the four groups (“responders” n=9, “non-responders” n=5): treated vs. untreated within each response class was tested using a paired Wilcoxon signed-rank test (paired by patient), whereas all other comparisons were tested using an unpaired Wilcoxon rank-sum test. P-values were Benjamini–Hochberg corrected within each pathway across the 6 comparisons. Adjusted significance is indicated as *FDR ≤ 0.05, **≤ 0.01), ***≤ 0.001), or ns. (not significant).

## Supplemental Tables and Table Legends

**Suppl. Table 1.** Differential expression of d17 bulk RNA.

**Suppl. Table 2.** GO terms BP of d17 bulk RNA.

**Suppl. Table 3.** GO terms MF of d17 bulk RNA.

**Suppl. Table 4.** GO terms CC of d17 bulk RNA.

**Suppl. Table 5.** GSEA of d17 single cell inflMo vs. hyinflMo.

**Suppl. Table 6.** GSEA of d17 single cell inflMo vs. IntMo.

**Suppl. Table 7.** GSEA of d17 single cell hyinflMo vs. IntMo.

**Suppl. Table 8.** Differential expression of d17 single cell hyinflMo vs. inflMo.

**Suppl. Table 9.** Differential expression of d17 single cell hyinflMo vs. IntMo.

**Suppl. Table 10.** Differential expression of d17 single cell IntMo vs. inflMo.

**Suppl. Table 11.** GSEA of d17 single cell hyinflMo (responders) treated vs. untreated.

**Suppl. Table 12.** GSEA of d17 single cell inflMo (non-responders) treated vs. untreated.

**Suppl. Table 13.** GSEA of d17 single cell IntMo (non-responders) treated vs. untreated.

**Suppl. Table 14.** Differential expression of d17 single cell hyinflMo (non-responders) treated vs. untreated.

**Suppl. Table 15.** Differential expression of d17 single cell inflMo (responders) treated vs. untreated.

**Suppl. Table 16.** Differential expression of d17 single cell IntMo (non-responders) treated vs. untreated.

**Suppl. Table 17.** GSEA of d17 single cell NK cells (non-responders) treated vs. untreated.

**Suppl. Table 18.** GSEA of d17 single cell NK cells (responders) treated vs. untreated

**Suppl. Table 19.** Differential expression of d17 single cell NK cells (non-responders) treated vs. untreated.

**Suppl. Table 20.** Differential expression of d17 single cell NK cells (responders) treated vs. untreated.

**Suppl. Table 21.** Differentially accessible regions d17 CD14^+^.

**Suppl. Table 22.** Differentially accessible regions d17 CD34^−^CD14^−^.

**Suppl. Table 23.** GO BP of d17 CD14^+^ (region based).

**Suppl. Table 24.** GO MF of d17 CD14^+^ (region based).

**Suppl. Table 25.** GO CC of d17 CD14^+^ (region based).

**Suppl. Table 26.** Differential expression of mouse BM cells WT vs. S100A9^−/−^.

**Suppl. Table 27.** GO terms BP of mouse BM cells WT vs. S100A9^−/−^.

**Suppl. Table 28.** GO terms MF of mouse BM cells WT vs. S100A9^−/−^.

**Suppl. Table 29.** GO terms CC of mouse BM cells WT vs. S100A9^−/−^.

**Suppl. Table 30.** Differential expression of GSE76203 responders 6 cycles vs. pre-treatment.

**Suppl. Table 31.** Differential expression of GSE76203 non-responders 6 cycles vs. pre-treatment.

**Suppl. Table 32.** GO terms BP of GSE76203 responders 6 cycles vs. pre-treatment.

**Suppl. Table 33.** GO terms MF of GSE76203 responders 6 cycles vs. pre-treatment.

**Suppl. Table 34.** GO terms CC of GSE76203 responders 6 cycles vs. pre-treatment.

**Suppl. Table 35.** Antibodies for FACS analysis.

**Suppl. Table 36.** Primers used for the MSP-PCR analysis of S100A8, S100A9 and S100A12. Primers were designed using the MethPrimer 2.0 Webtool (http://www.urogene.org/methprimer2/)^6^ F= forward primer; R= reverse primer. * published primer, ^7^.

**Suppl. Table 37.** ADT sequences for CITEseq

## Notes

**Grant support:** This work was supported by grants from the Deutsche Kinderkrebsstiftung (2014.03 and 2021.09), the Sander Stiftung (2018.112.1), the Deutsche Forschungsgemeinschaft (AN 849/1-2 and 849/3-1), Else Kröner Fresenius Stiftung (2102_A296), the Jose Carreras Leukämie Stiftung (SP 02/2020), the Faculty of Medicine of Tuebingen FO_2013_01_14, and the „Stiftung des Fördervereins für krebskranke Kinder Tübingen” all to MCA, the Fortune program (3038-0-0) to RS and (2392-0-0) to AK (Kübler), and the Faculty of Medicine of Tuebingen (2023-0-01) to DS, (2298-0-0) to AK (Kuru), (2020-2-26) to BFB, (2022-1-14) to PK, (2024-2-26) to NG. NC was financially supported by the DFG-funded NGS Competence Center Tübingen (INST 37/1049-1).

### Competing Interest Statement

The authors have declared no competing interest.

