## Supplemental Figures for "Epigenetic control of S100A8/A9-driven monocytic inflammation licenses anti-leukemic functionality of immature NK cells during hematopoietic stem cell differentiation"

### Suppl. Fig. S1.

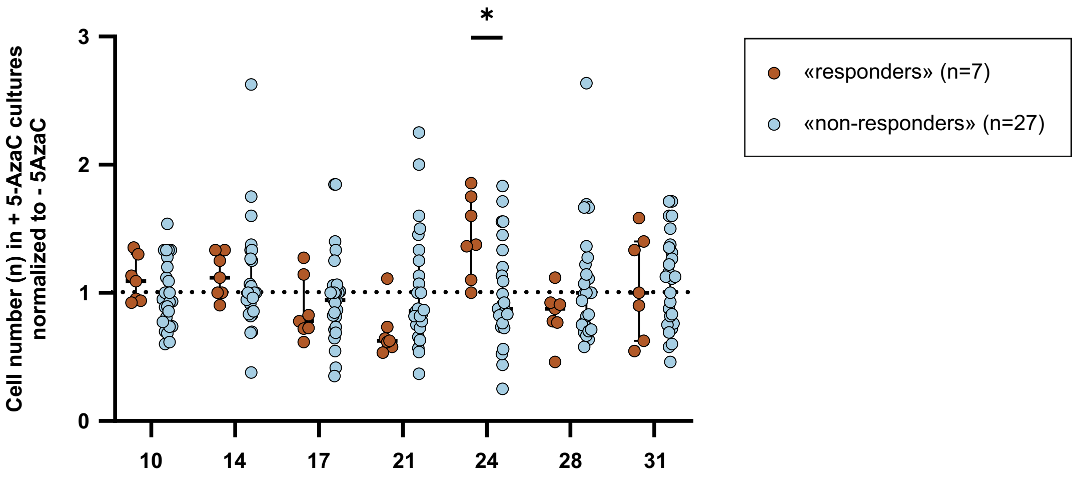

*
Cell number (n) in + 5-AzaC cultures normalized to - 5AzaC
3
2
1
0
10
14
17
21
24
28
31
«responders» (n=7)
«non-responders» (n=27)

### Suppl. Fig. S2.

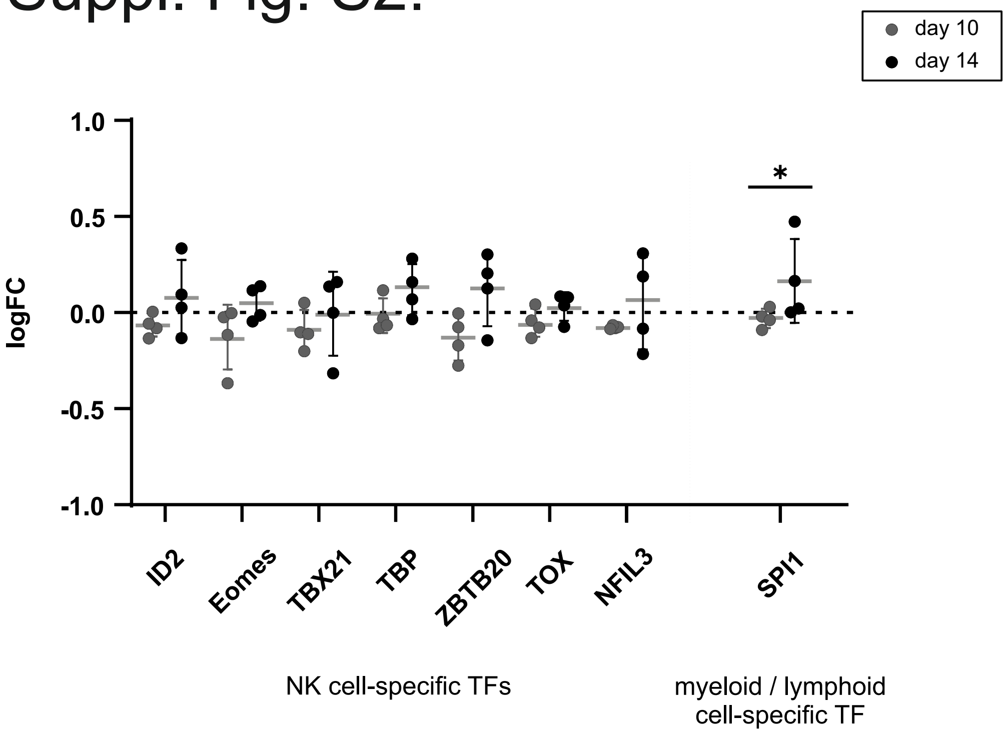
day 10
day 14
1.0
0.5
0.0
-0.5
-1.0
logFC
*
ID2
Eomes
TBX21
TBP
ZBTB20
TOX
NFIL3
SPI1
NK cell-specific TFs
myeloid / lymphoid
cell-specific TF

### Suppl. Fig. S3.

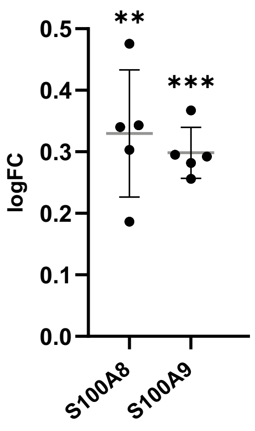

### Suppl. Fig. S4.

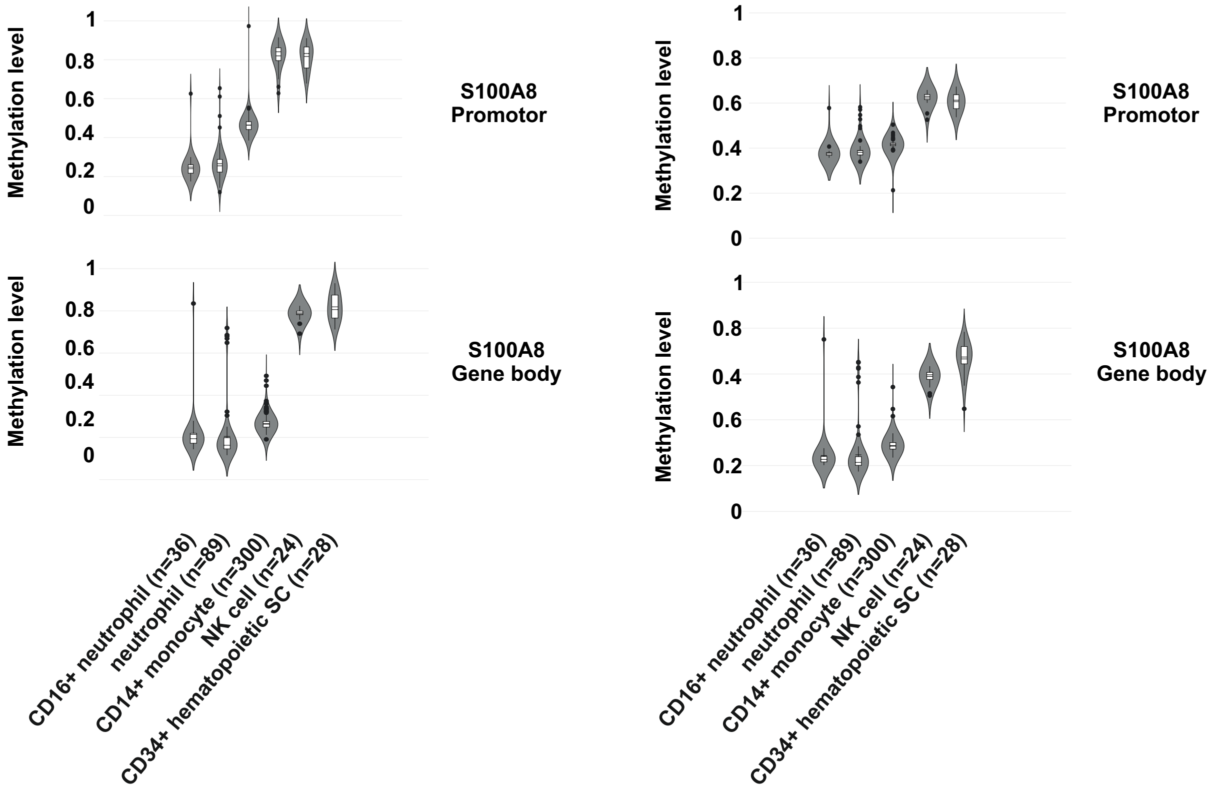
Methylation level
1
0.8
0.6
0.4
0.2
0
S100A8
Promotor
S100A8
Gene body
CD16+ neutrophil (n=36)
neutrophil (n=89)
CD14+ monocyte (n=300)
NK cell (n=24)
CD34+ hematopoietic SC (n=28)

### Suppl. Fig. S5.

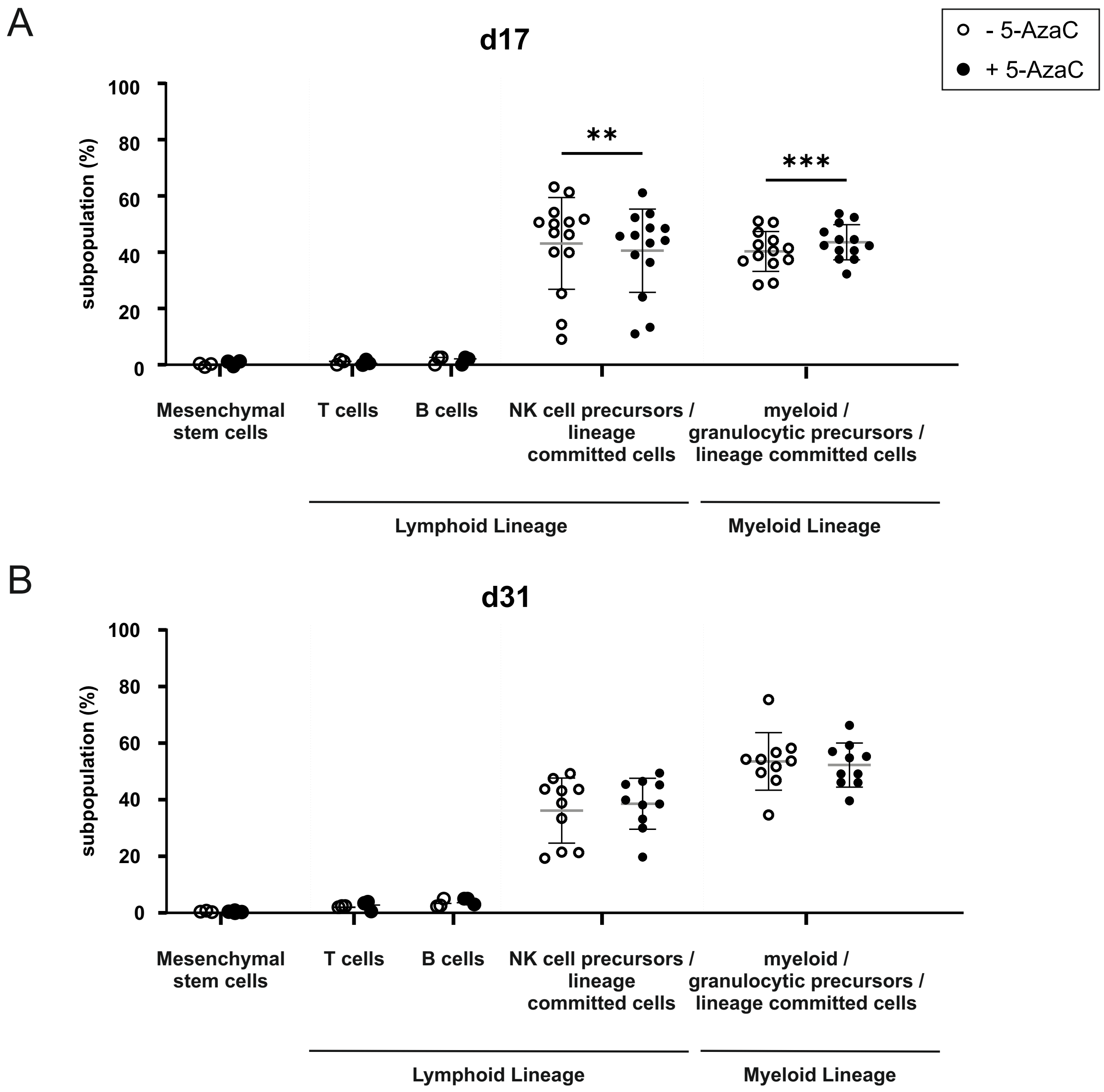
A
d17
- 5-AzaC
+ 5-AzaC
subpopulation (%)
100
80
60
40
20
0
**
***
Mesenchymal stem cells
T cells
B cells
NK cell precursors / lineage committed cells
myeloid / granulocytic precursors / lineage committed cells
Lymphoid Lineage
Myeloid Lineage
B
d31
subpopulation (%)
100
80
60
40
20
0
Mesenchymal stem cells
T cells
B cells
NK cell precursors / lineage committed cells
myeloid / granulocytic precursors / lineage committed cells
Lymphoid Lineage
Myeloid Lineage

### Suppl. Fig. S6.

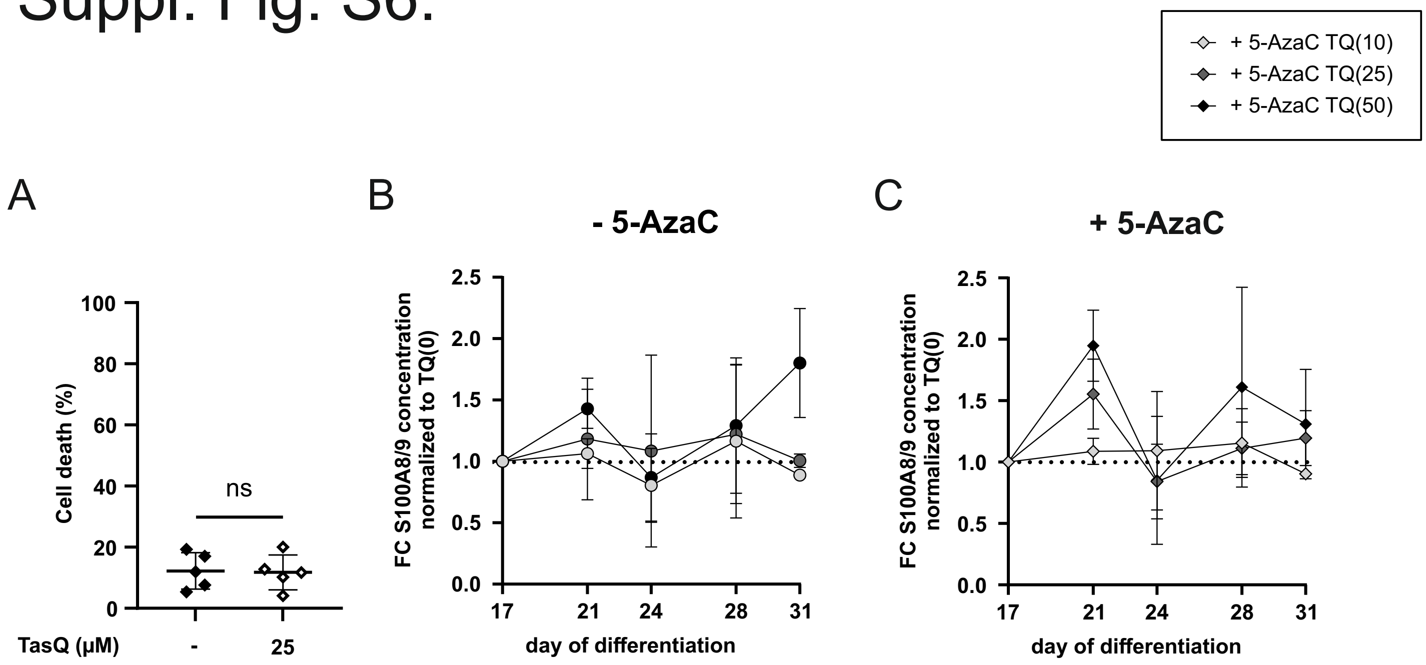
+ 5-AzaC TQ(10)
+ 5-AzaC TQ(25)
+ 5-AzaC TQ(50)
A
Cell death (%)
100
80
60
40
20
0
ns
TasQ (µM)
-
25
B
- 5-AzaC
FC S100A8/9 concentration normalized to TQ(0)
2.5
2.0
1.5
1.0
0.5
0.0
17
21
24
28
31
day of differentiation
C
+ 5-AzaC
FC S100A8/9 concentration normalized to TQ(0)
2.5
2.0
1.5
1.0
0.5
0.0
17
21
24
28
31
day of differentiation

### Suppl. Fig. S7.

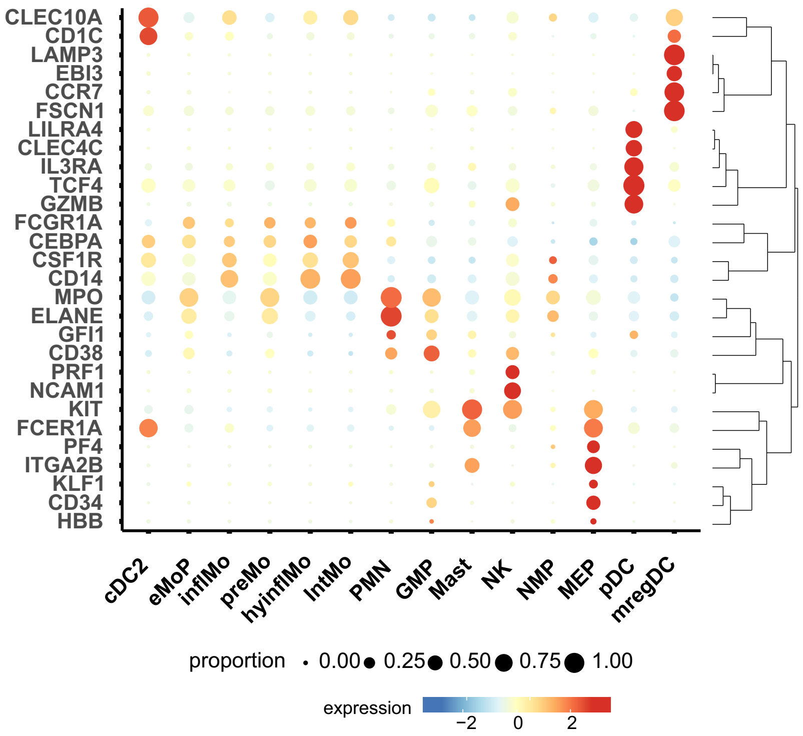
CLEC10A
CD1C
LAMP3
EBI3
CCR7
FSCN1
LILRA4
CLEC4C
IL3RA
TCF4
GZMB
FCGR1A
CEBPA
CSF1R
CD14
MPO
ELANE
GFI1
CD38
PRF1
NCAM1
KIT
FCER1A
PF4
ITGA2B
KLF1
CD34
HBB
cDC2
eMoP
inflMo
preMo
hyinflMo
IntMo
PMN
GMP
Mast
NK
NMP
MEP
pDC
mregDC
proportion
0.00
0.25
0.50
0.75
1.00
expression
−2
0
2

### Suppl. Fig. S8.

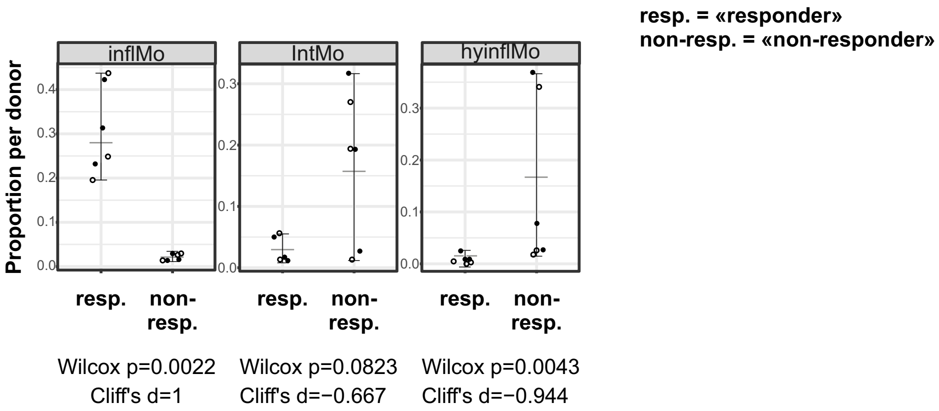

resp. = «responder»
non-resp. = «non-responder»
inflMo
IntMo
hyinflMo
Proportion per donor
resp.
non-resp.
Wilcox p=0.0022
Cliff's d=1
Wilcox p=0.0823
Cliff's d=−0.667
Wilcox p=0.0043
Cliff's d=−0.944

### Suppl. Fig. S9.

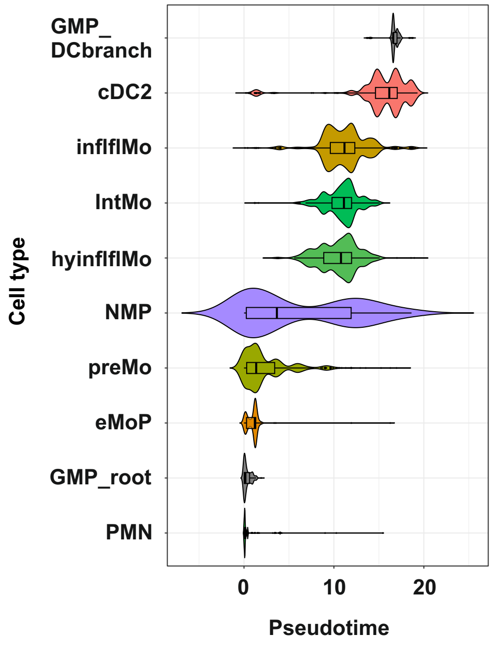
GMP_
DCbranch
cDC2
inflflMo
IntMo
hyinflflMo
NMP
preMo
eMoP
GMP_root
PMN
Cell type
0
10
20
Pseudotime

### Suppl. Fig. S10.

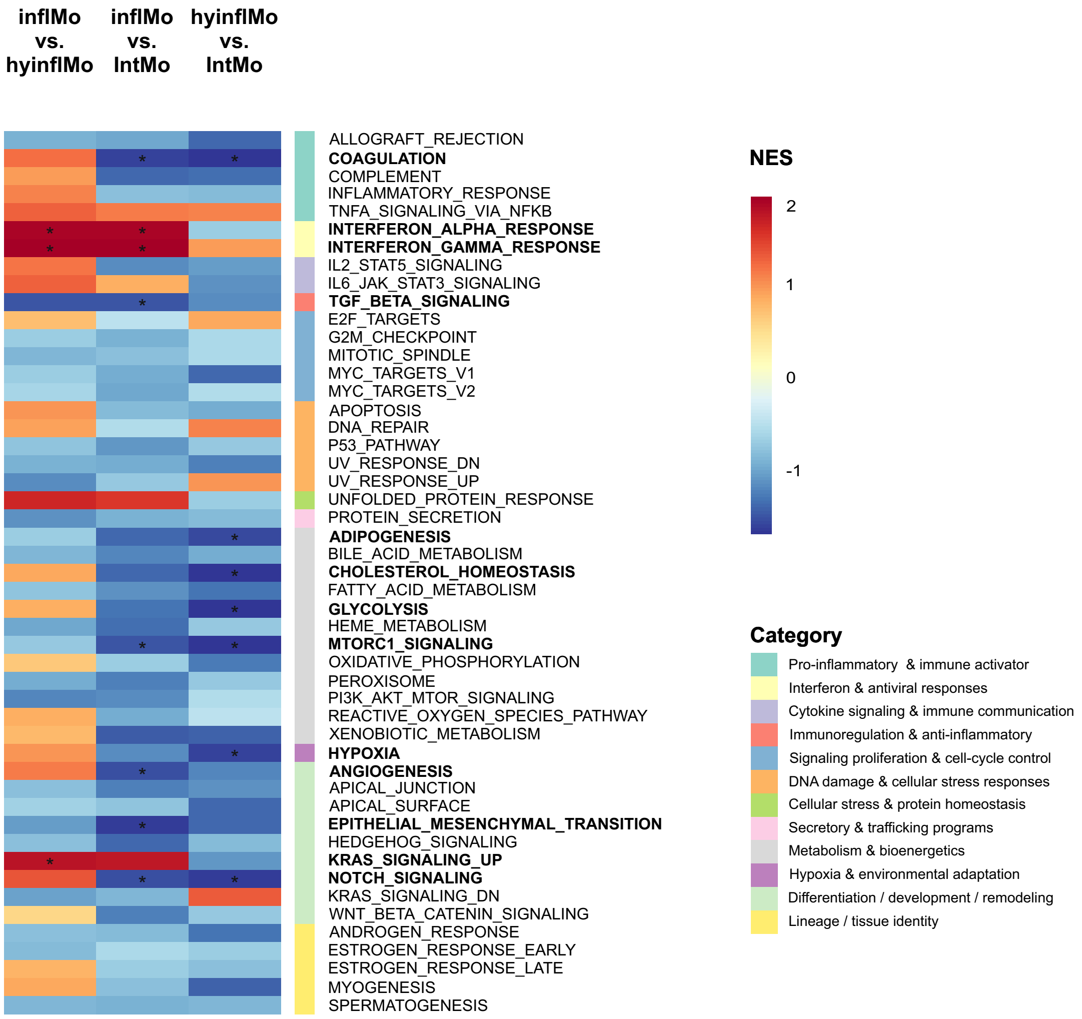

### Suppl. Fig. S11.

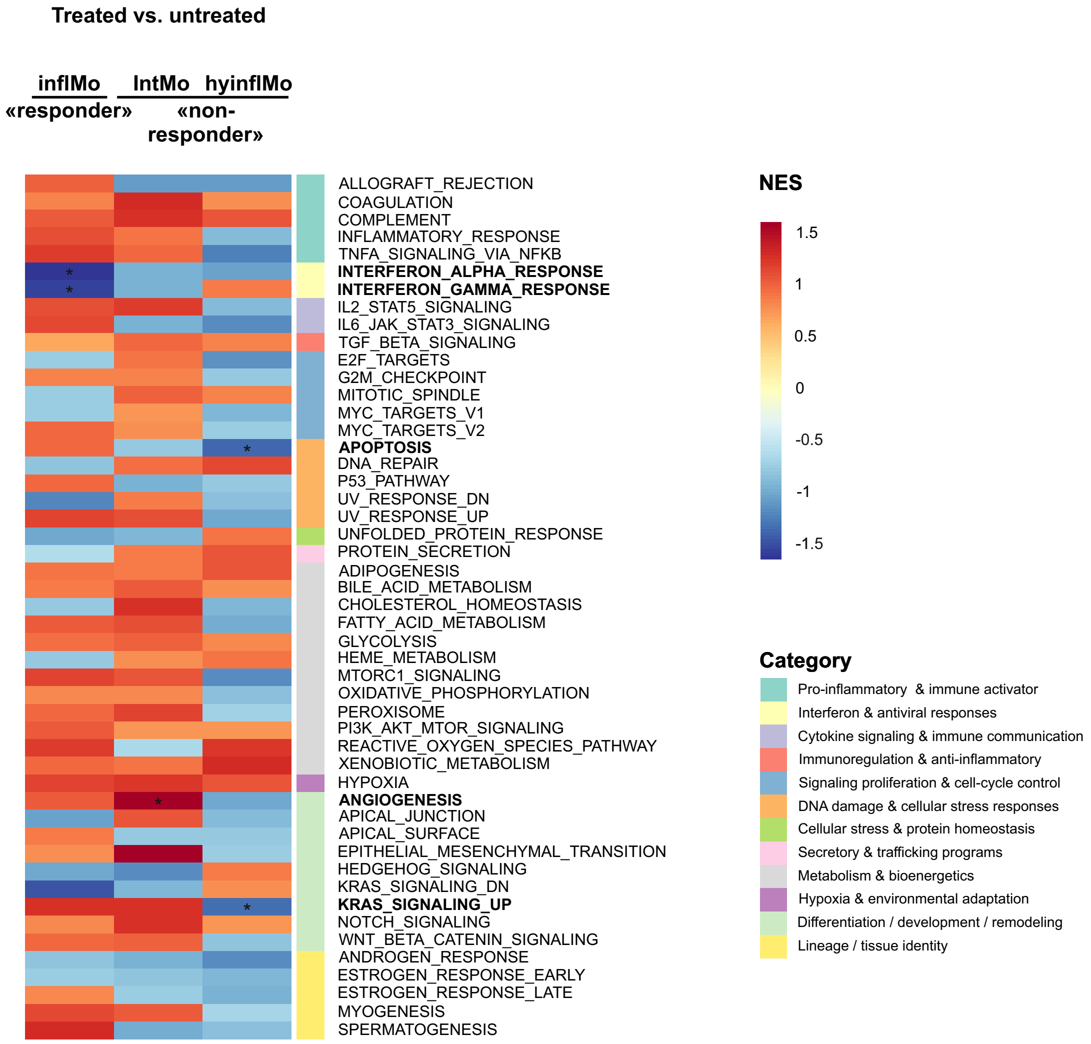

### Suppl. Fig. S12.

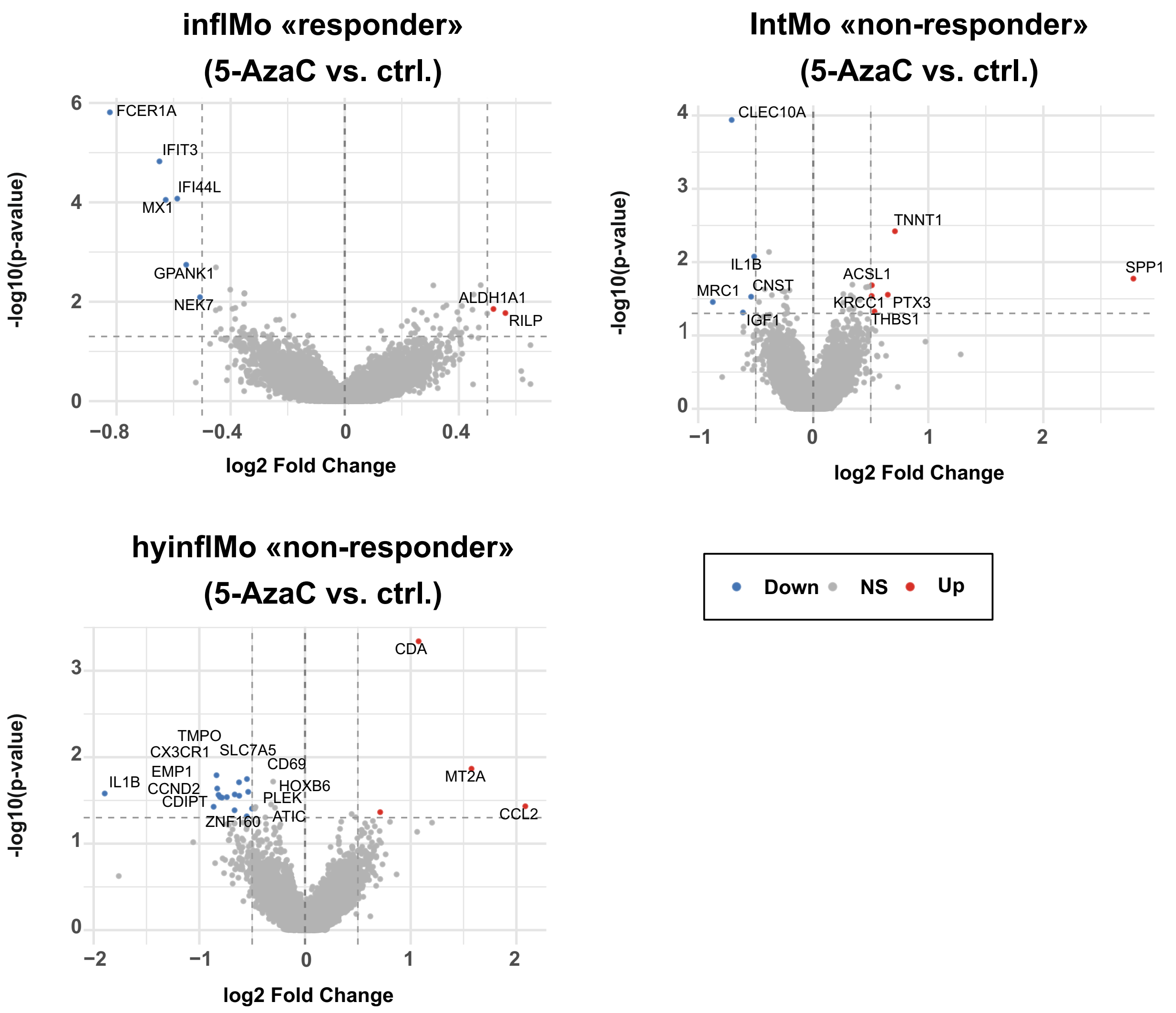

### Suppl. Fig. S13.

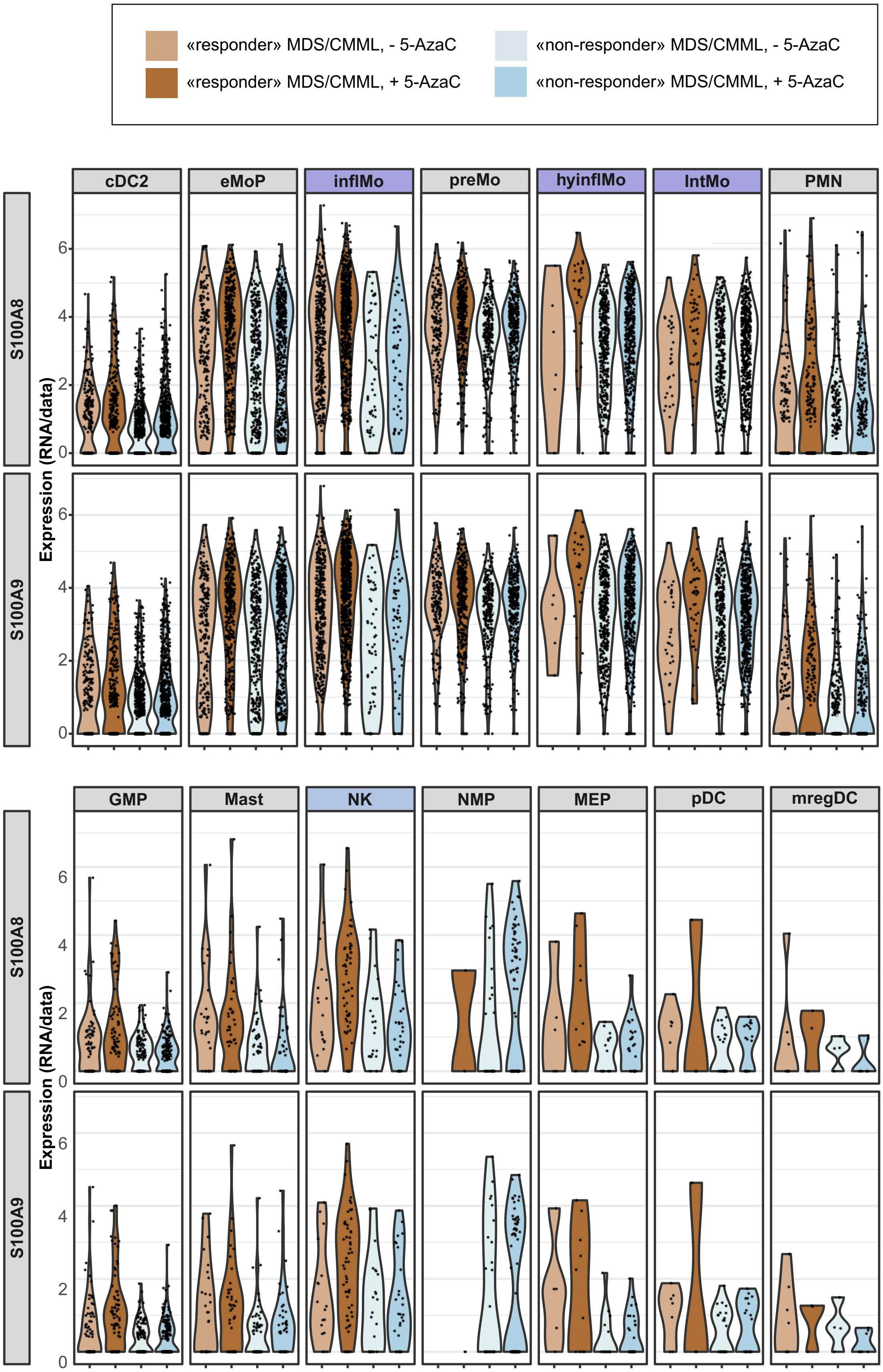

### Suppl. Fig. S14.

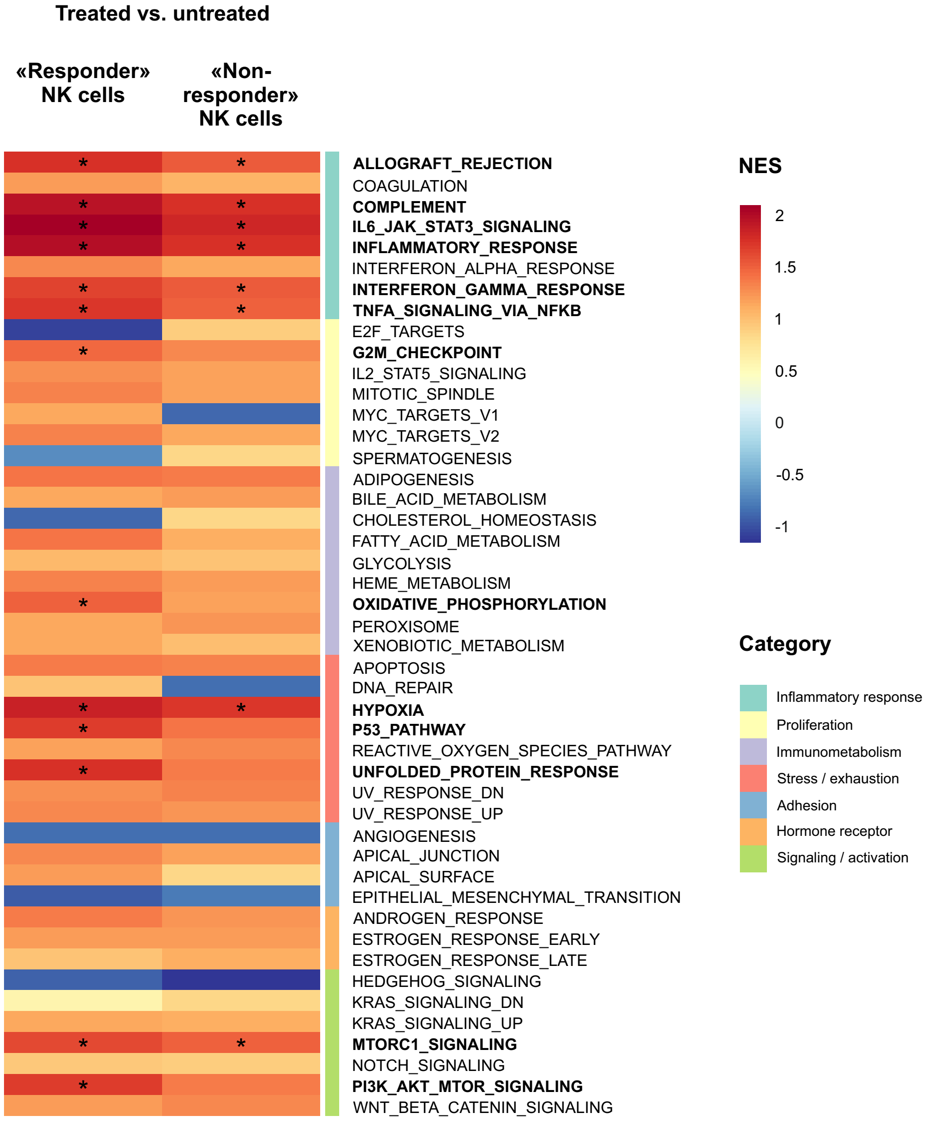

### Suppl. Fig. S15.

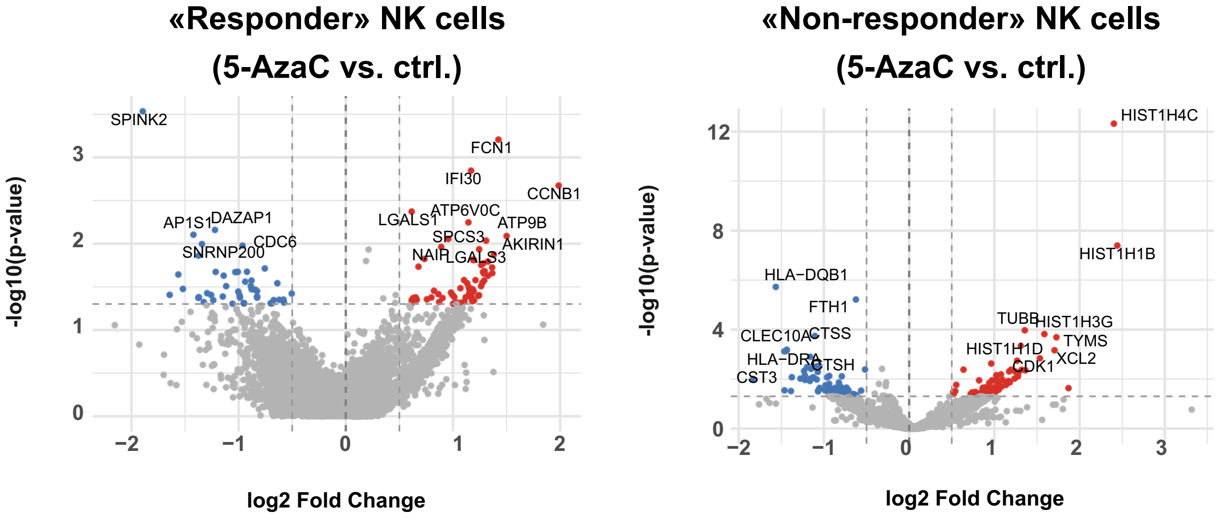

### Suppl. Fig. S16.

A

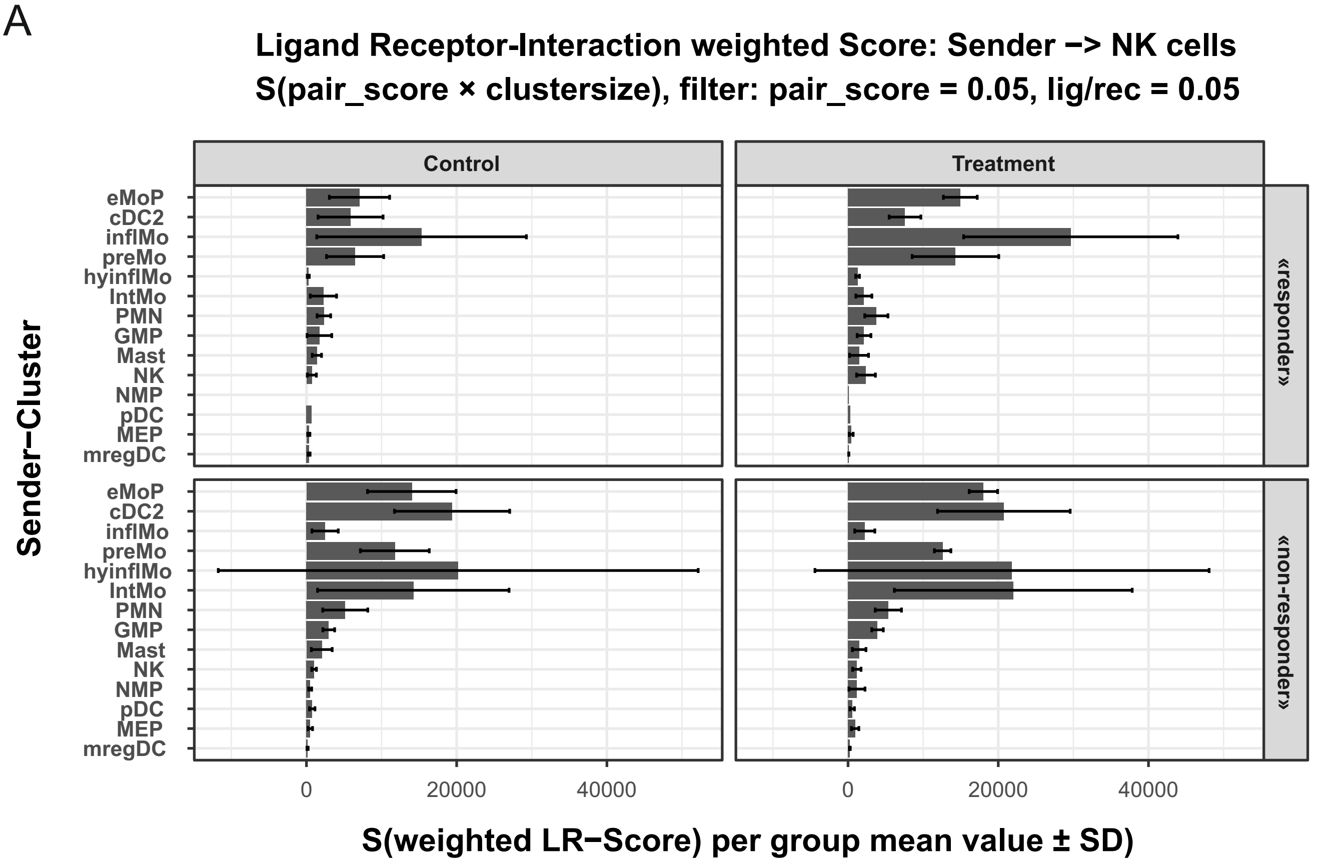


B

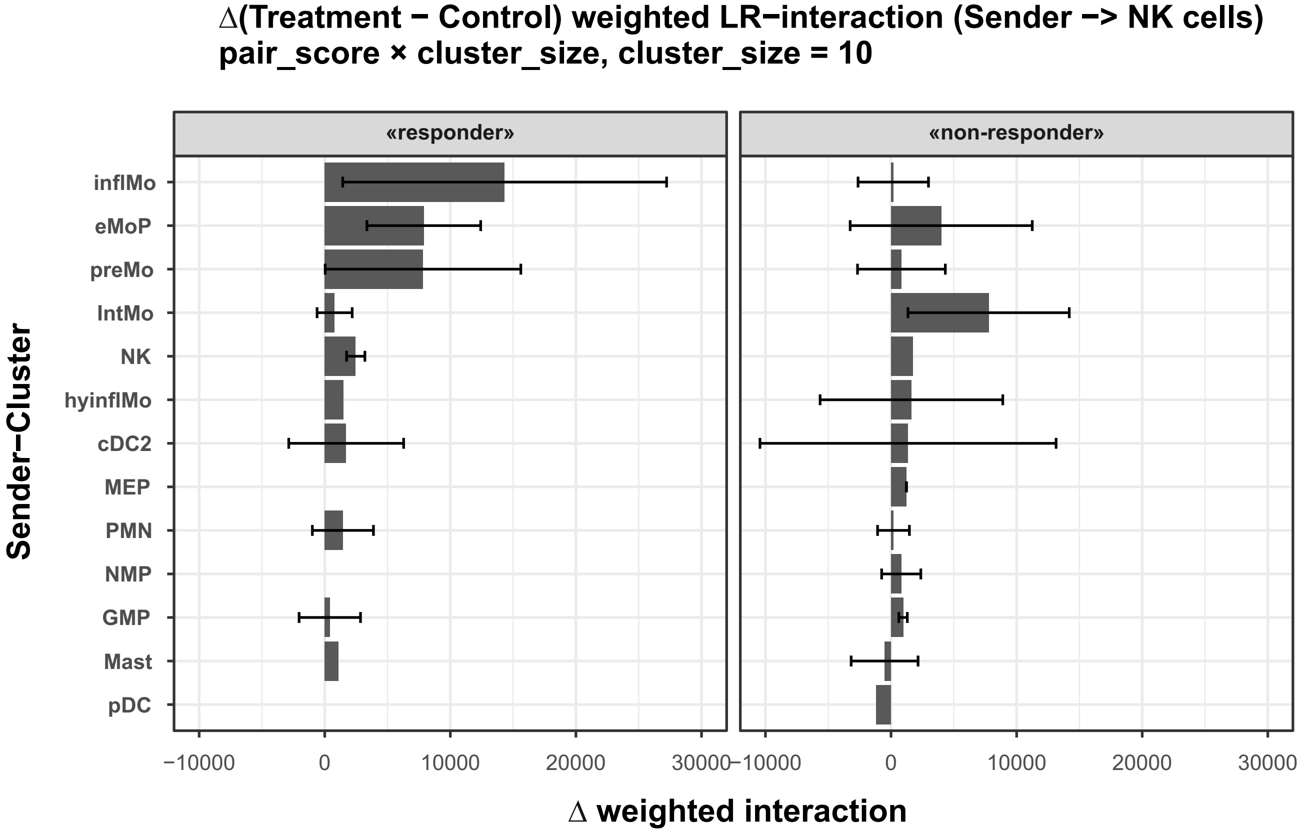

### Suppl. Fig. S17.

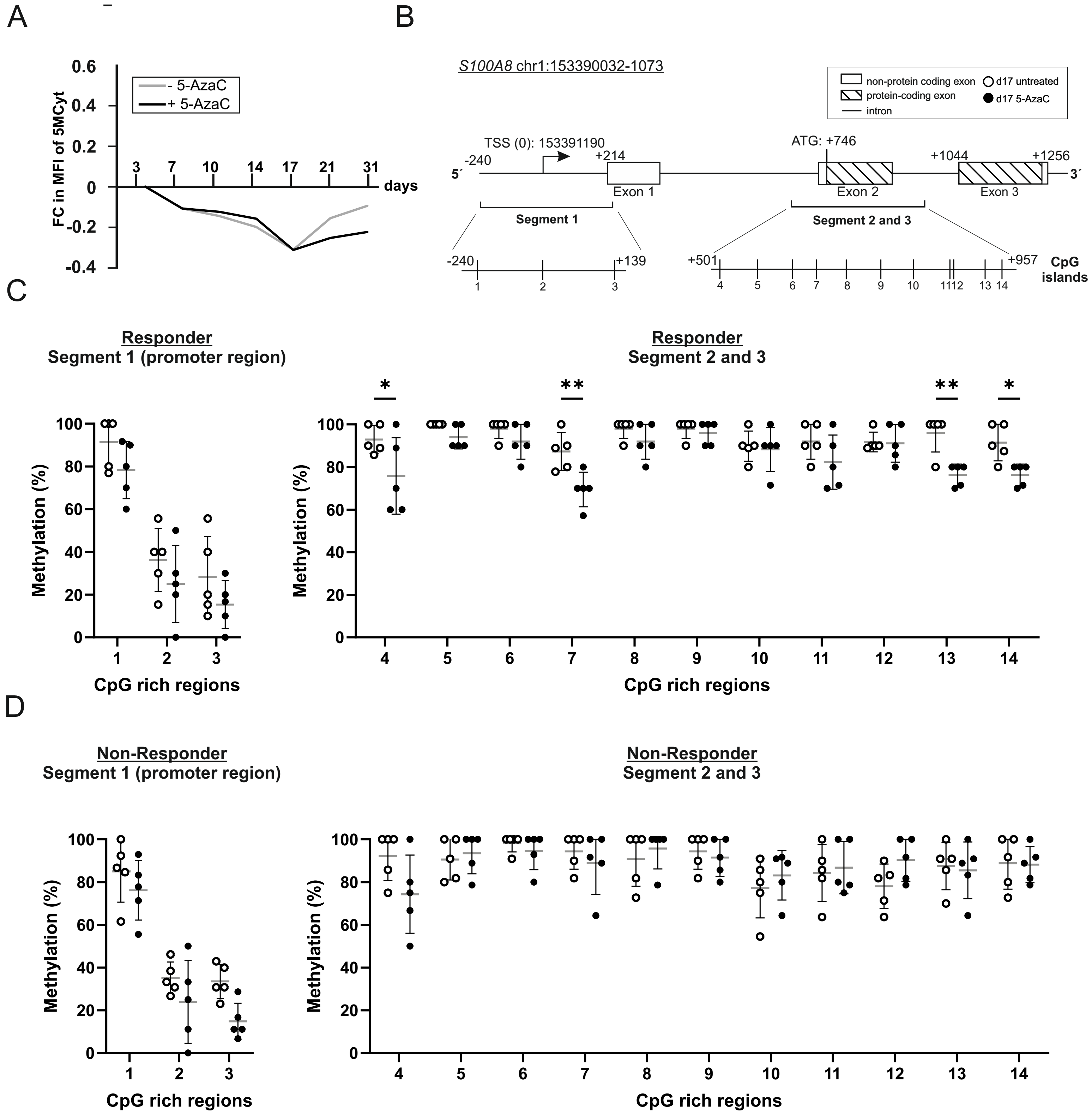

### Suppl. Fig. S17, cont.

E

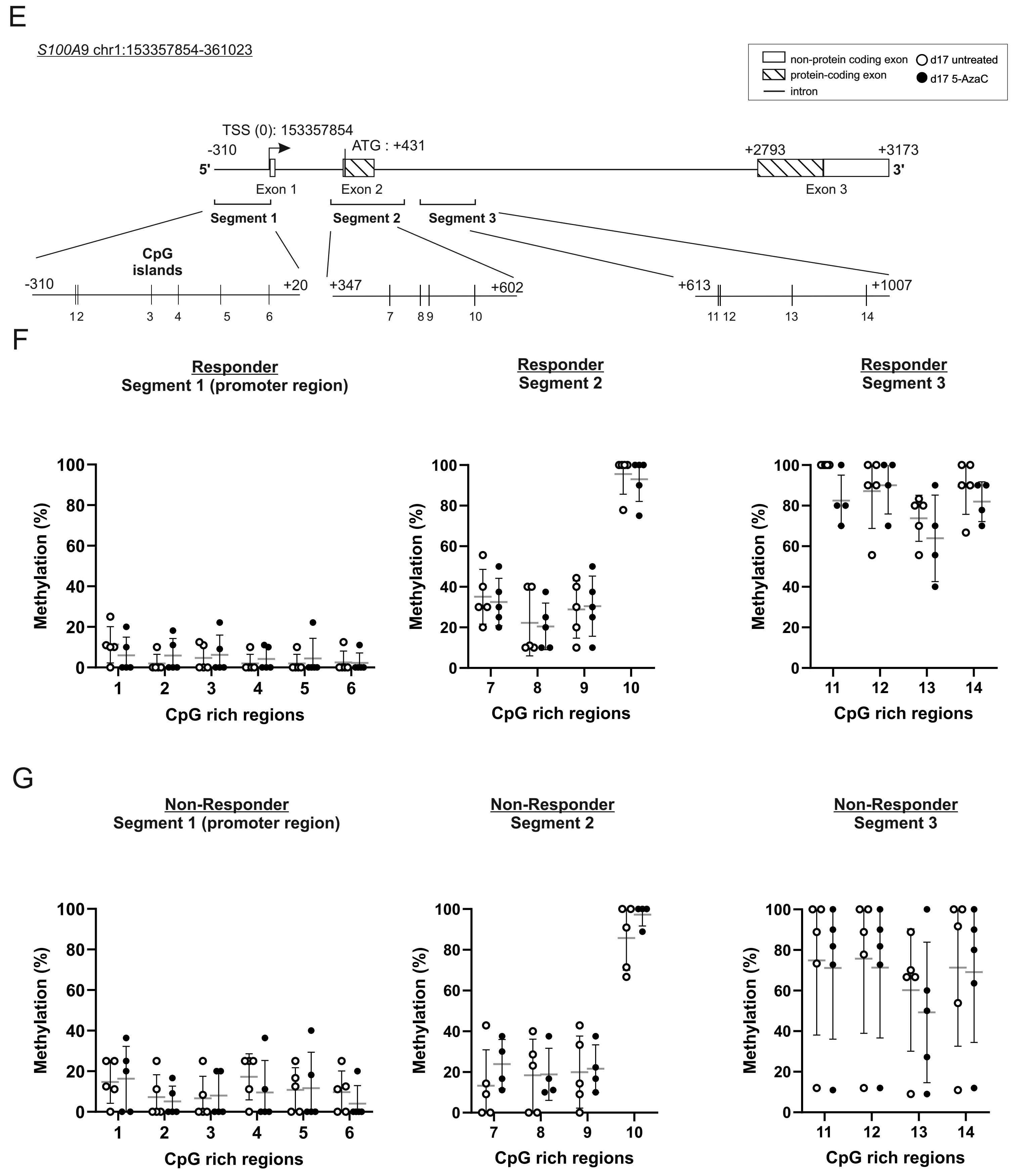

### Suppl. Fig. S18.

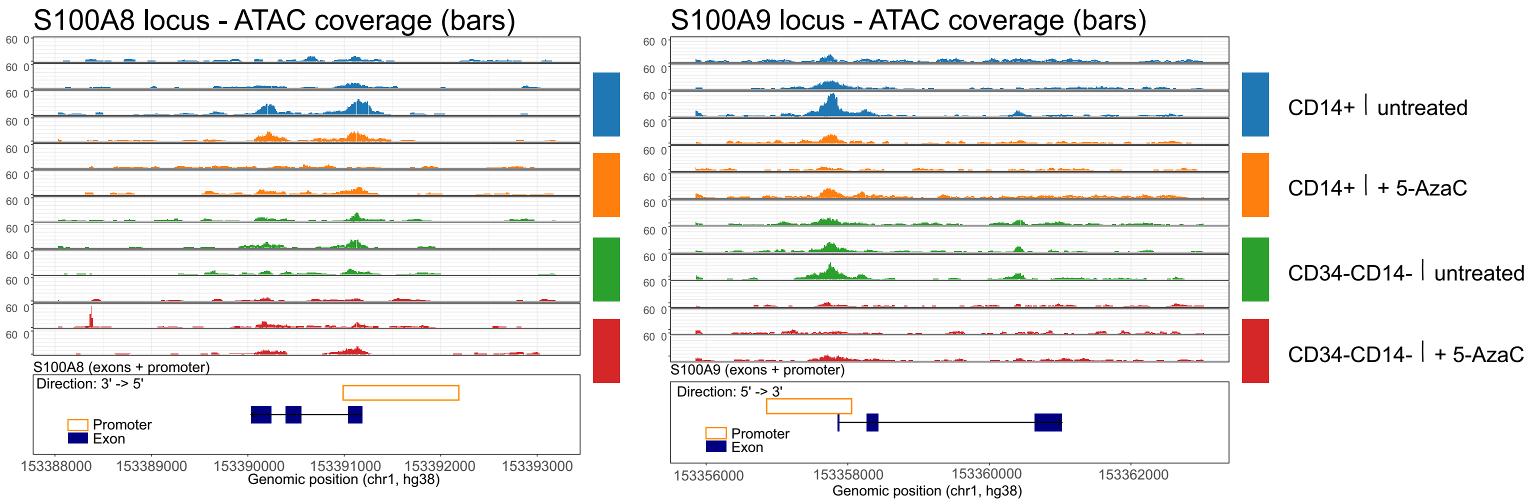

### Suppl. Fig. S19.

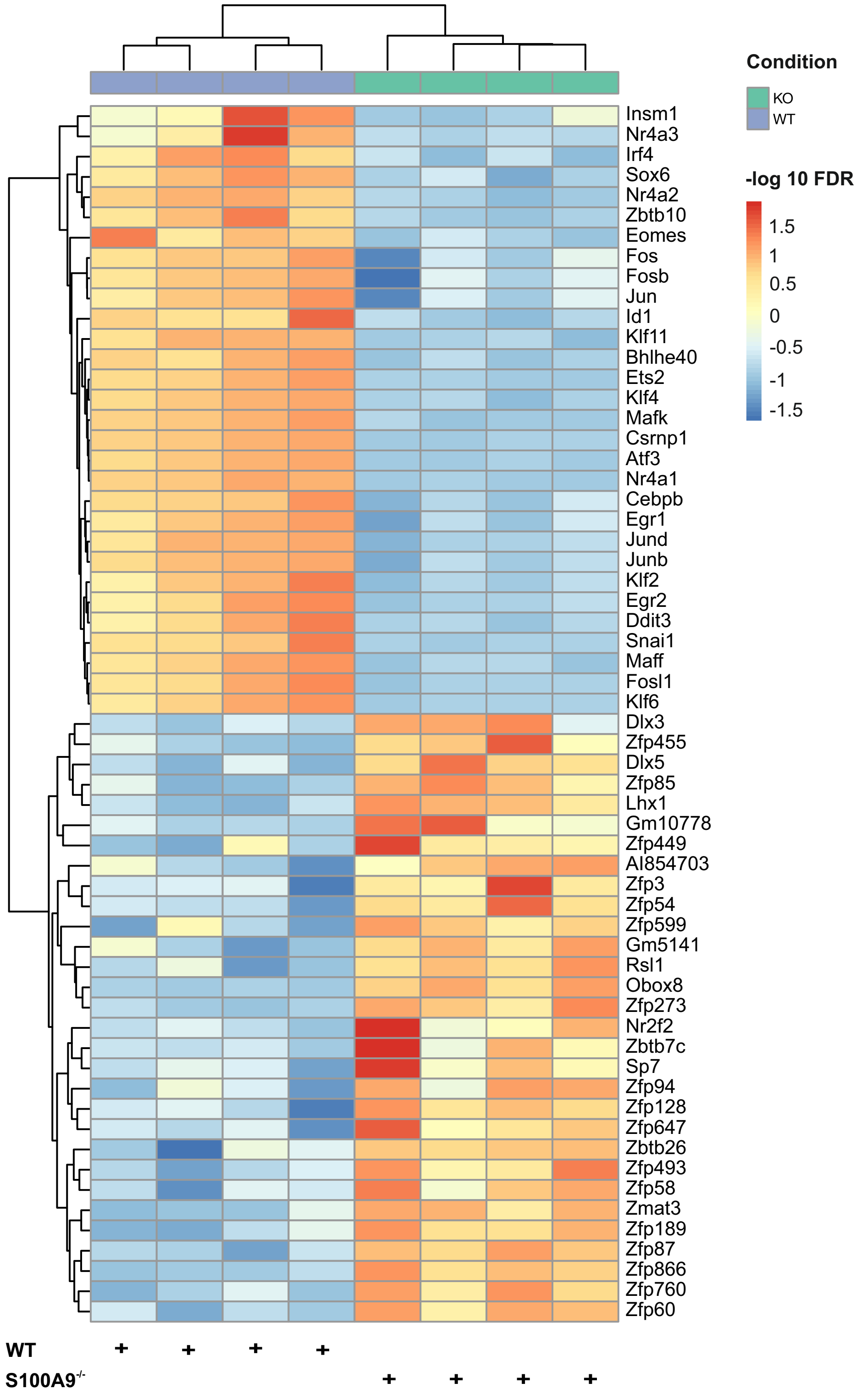

### Suppl. Fig. S20.

A

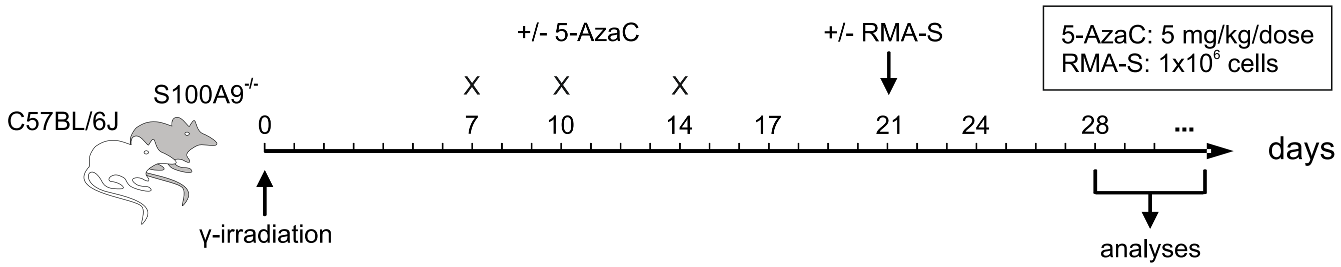


B

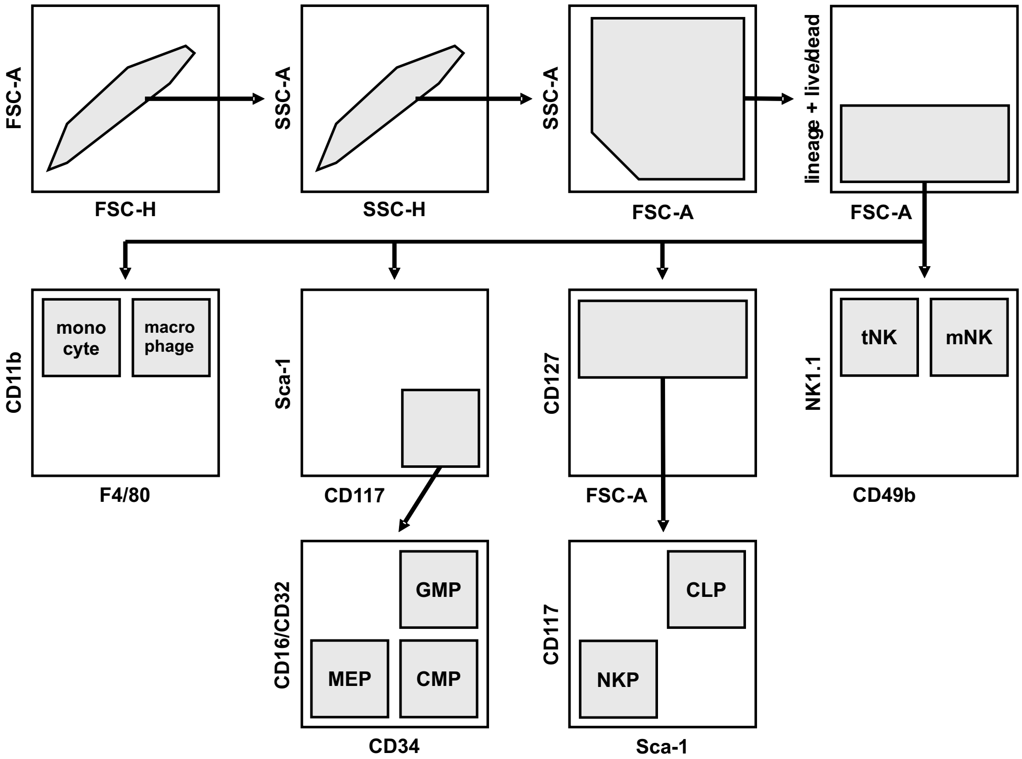

### Suppl. Fig. S21.

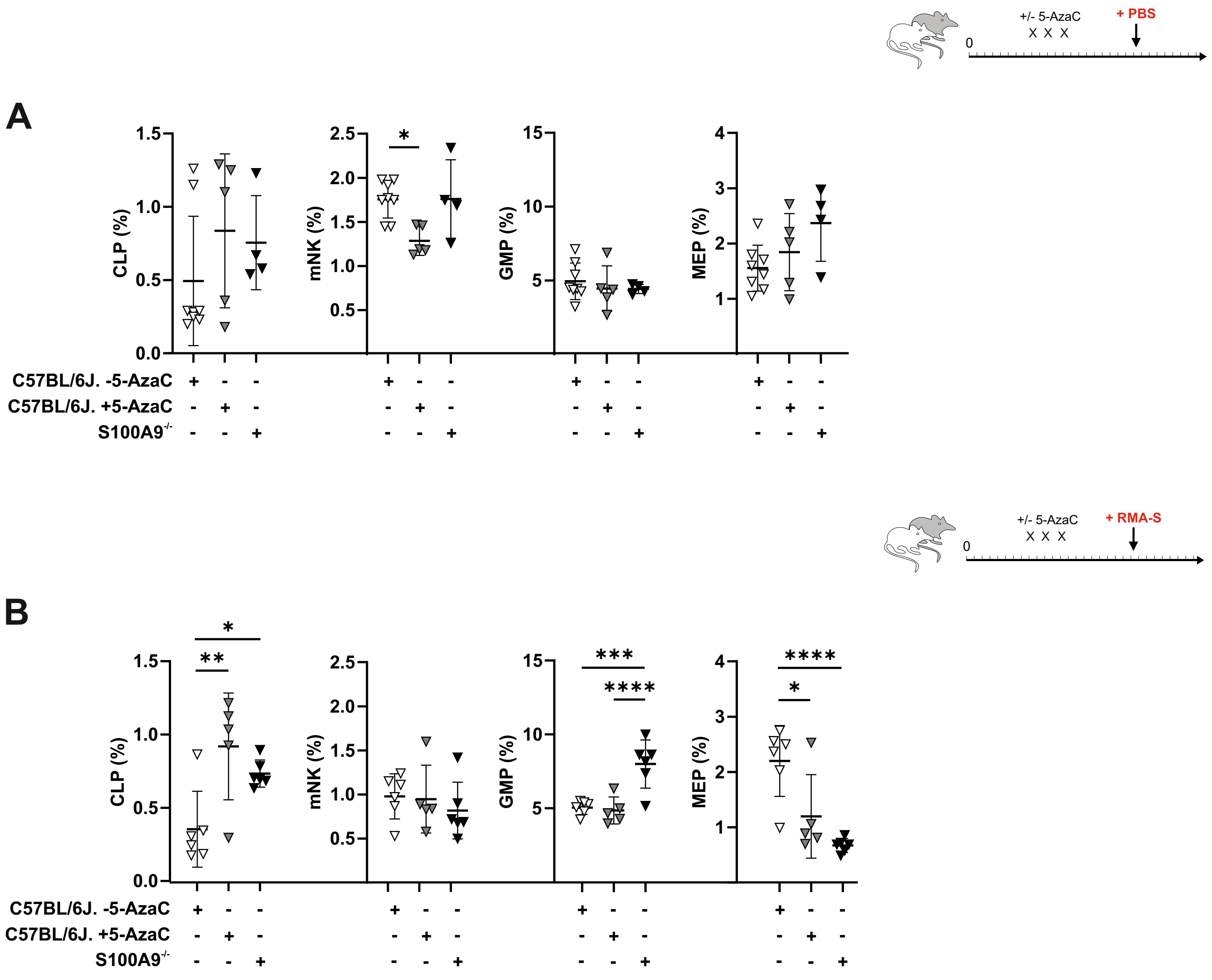

### Suppl. Fig. S22.

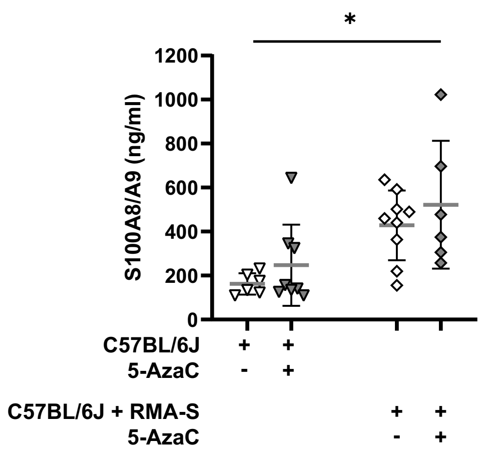

### Suppl. Fig. S23.

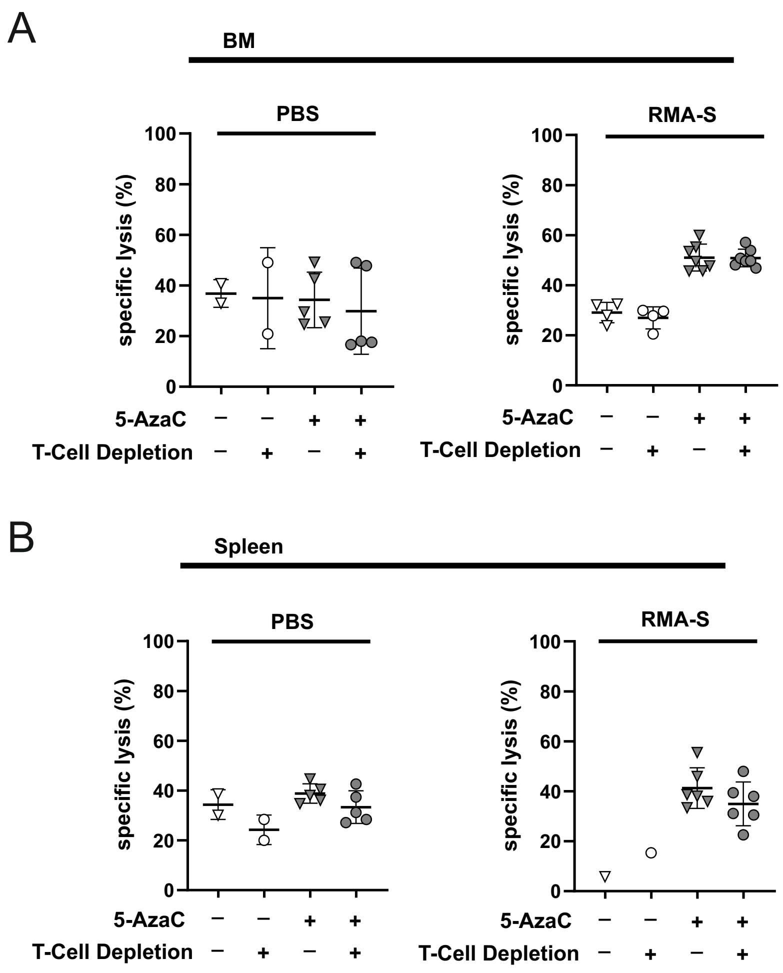

### Suppl. Fig. S24.

A

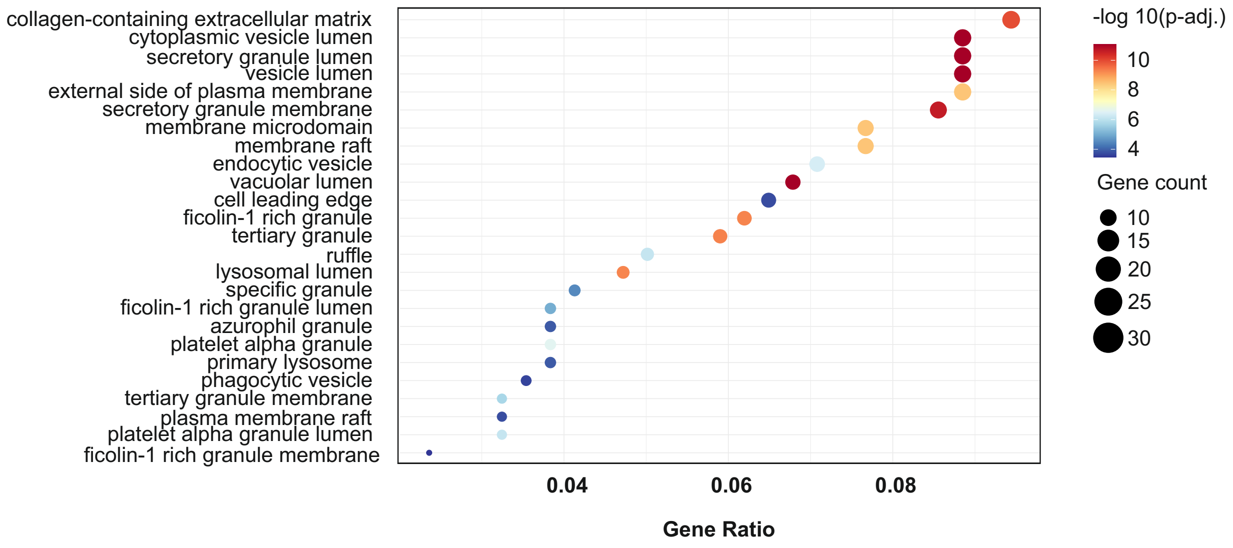


B

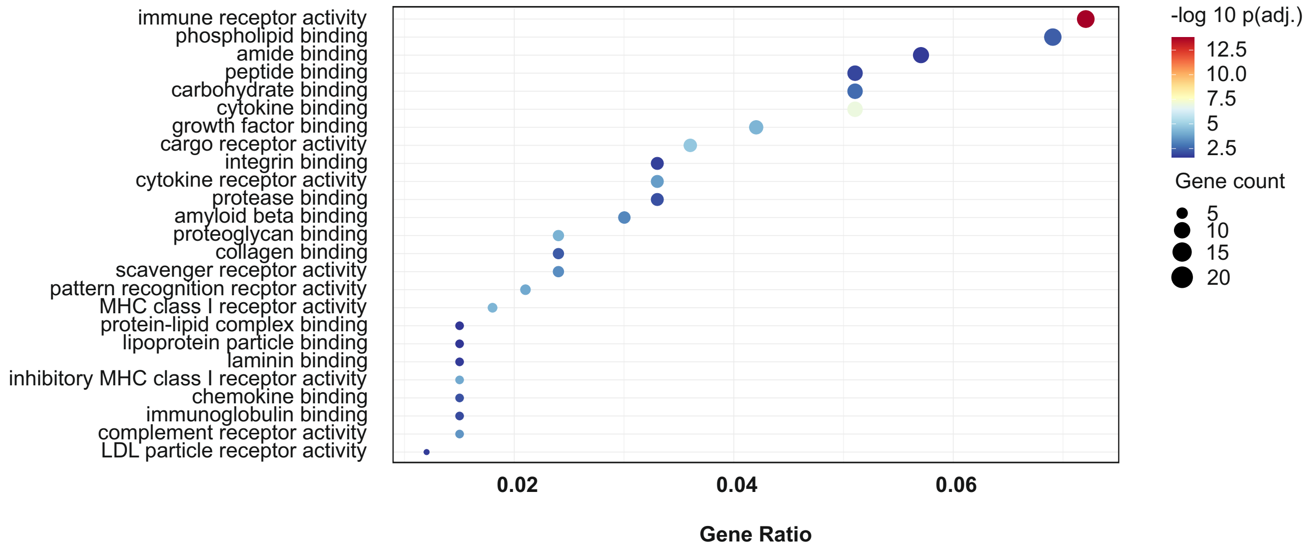

### Suppl. Fig. S25.

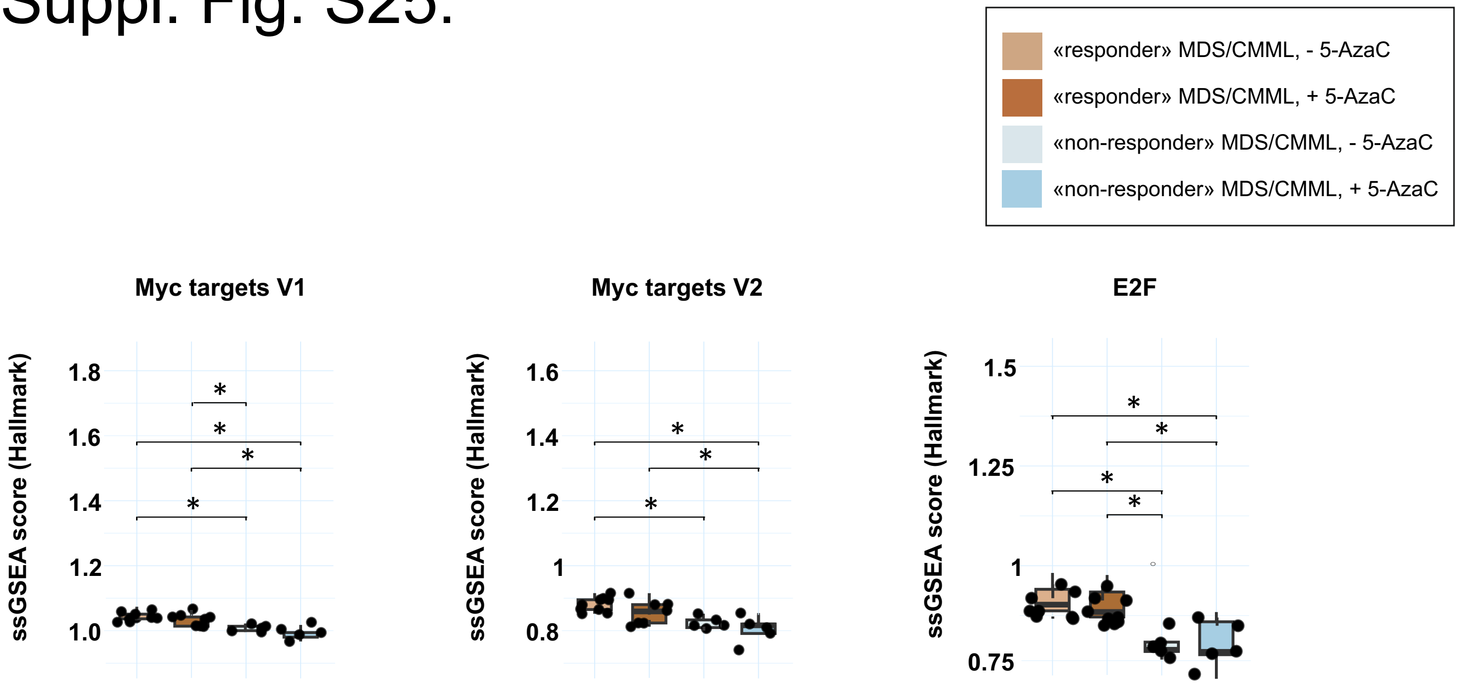
